# Ancient human genomes from the Altai region reveal population continuity and shifts in the 4th-12th centuries

**DOI:** 10.64898/2026.03.24.713889

**Authors:** Yusuf Can Özdemir, Balázs Gyuris, Kristóf Jakab, Tamás Szeniczey, Botond Heltai, Melinda Megyes, Balázs G. Mende, István Major, Nikolai Seregin, Vadim V. Gorbunov, Sergey Grushin, Petr K. Dashkovskiy, Mikhail A. Demin, Kirill Yu. Kiryushin, Yury T. Mamadakov, Nadezhda F. Stepanova, Svetlana S. Tur, Alexey V. Fribus, Marina P. Rykun, Attila Türk, Alexey A. Tishkin, Anna Szécsényi-Nagy

## Abstract

The Altai region is a crossroads of the Asian steppes. However, the population history of the region remains understudied. We analyse ancient human genomes from the Altai and Ob regions, creating a ca. 1400-year-long time transect with 91 new data. We demonstrate an Iron Age genetic variety that continued into the Medieval era, with additional large-scale spread of East Asian genetic ancestry coinciding with the rise and spread of the Turkic cultural customs. Furthermore, we find a unique lineage in the Early Medieval Altai with elevated Ancient North Eurasian ancestry, providing a missing link between the North Eurasian hunter-gatherers and modern North Asian people. We identify distinct genetic patterns and connections among populations of the Mountainous and the Forest-Steppe Altai in the 4th-8th centuries.

## Introduction

The Altai Mountains span 2,000 kilometers in the territories of Russia, Mongolia, China and Kazakhstan in Inner Asia. To the north of this range, the Forest-Steppe Altai region marks the easternmost end of the Central Steppe, and the Ob River plateau stretches farther north (Fig. 1). The Altai region encompasses both the Mountainous and Forest-Steppe Altai geographical areas, and is a crossroads between the Eastern and Central Steppes of Eurasia. Although a key area for Inner Asian population histories, the understanding of the Altai remains incomplete, partially due to the limited number of published ancient genomes from the narrower geographical area. Considering previous genetic research on the Bronze Age (BA) and post-BA Inner Asia is essential to improve our understanding. The Iron Age (IA) populations of Inner Asia can be traced back to impactful basal genetic components detailed in Box 1, after which Inner Asia continuously experienced gene flow from East Asia, and from South-Central Asia (*1–4*). The post-IA Inner Asians, namely the Xiongnu period and Medieval peoples exhibited substantial genetic heterogeneity throughout the region (*1–5*).

**Figure 1:**
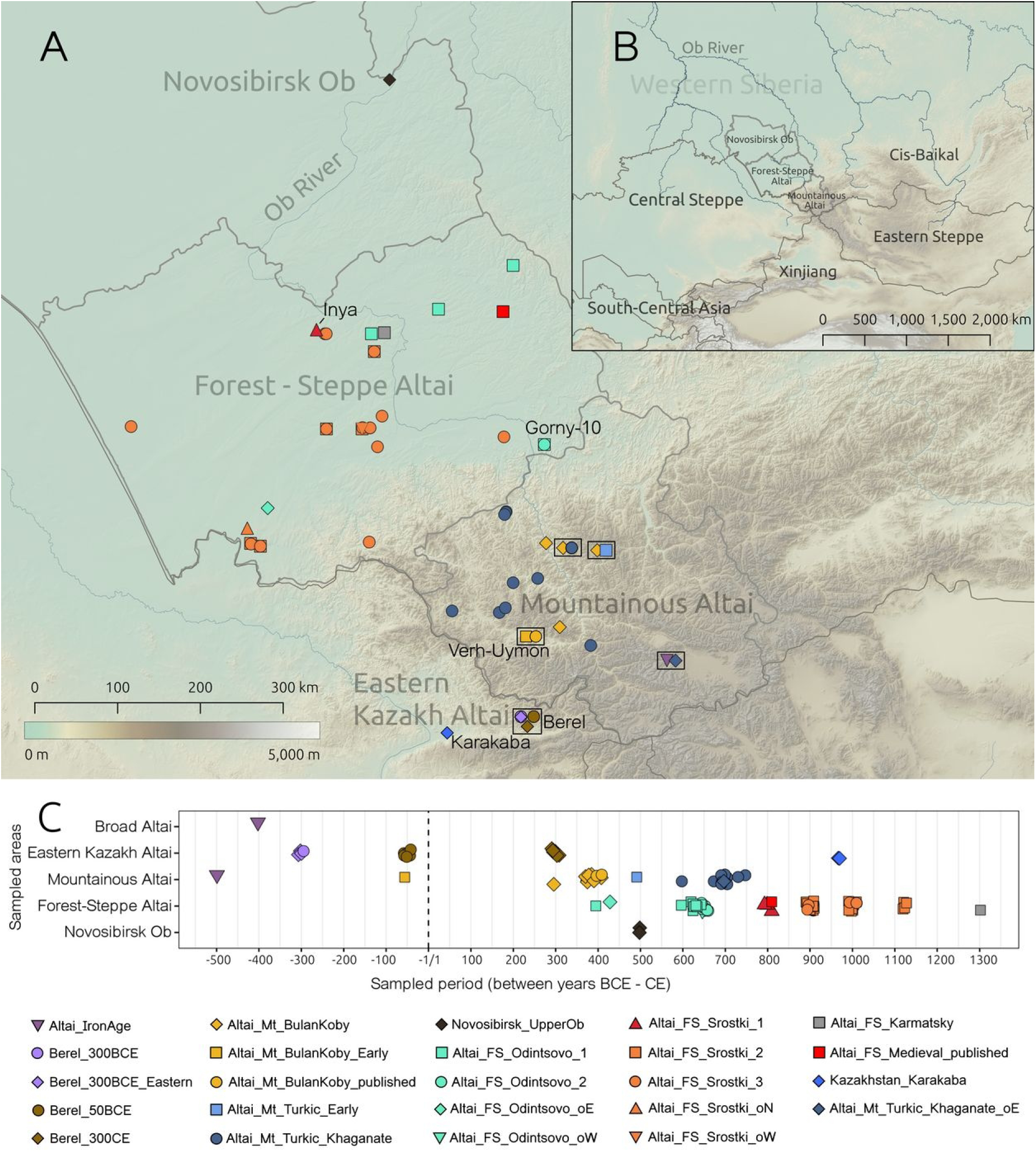
Sampled burial sites and chronology of the sampled individuals. (A) Map of the archaeological sites where newly-reported and previously-published Altaian individuals were excavated. Different periods sampled from the same archaeological sites are jittered and shown in boxes. Sites with multiple genetic profiles from the presented periods have overlapping symbols. The symbols represented on the map are the same as those used in the legend at the bottom of the figure. Borders of internal regions of modern Russia are shown in addition to the country borders. (B) A broad-scale map of the country borders and names, and contextually-important regions. (C) Chronology of the newly reported and published ancient groups from the Altai and Upper Ob Plateau. The symbols and color scheme used are the same as those used in Fig. 2.

### Box 1

Genetic background of Inner Asia

The Iron Age (IA) Inner Asian gene pool was shaped by admixture events of distinct Neolithic – Bronze Age (BA) Asian ancestries: Ancient Northeast Asian (ANA) forms the core of the East Asian ancestries in the IA Asian steppes (*3*). The Ancient North Eurasian (ANE) component is first described in Palaeolithic Siberians (*6*), then found in the West Siberian hunter-gatherers and Eneolithic Botai culture’s people (WSHG_Botai) and in the Altaian Neolithic hunter-gatherers (Altai_HG) (*7–11*). Altai_HG and WSHG_Botai represent pre-Bronze Age ancestries of the Altai region and of more westerly parts of Southern Siberia, respectively, and belong to the recently-defined North Eurasian hunter-gatherer cline (NEAHG) (*12*) (Fig. S3). This cline was later disrupted by the expansions of ANA-related Neolithic – Bronze Age Yakutians and Cis-Baikalians (*12*), as well as by the Early Bronze Age Western Steppe Herders associated with the Yamnaya culture (Steppe_EBA) and related Bronze Age groups (*9*). Finally, the South-Central Asian components are linked to the people of the Bactria-Margiana Archaeological Complex (BMAC) and their descendants (*1–4*, *9*, *13*).

During the IA, most of the Asian steppe region and the Altai were dominated by the broad ‘Eastern Scythian - Saka’ cultural sphere, and the Altai area attests to the Scythian phenomenon’s earliest emergence (Alekseev et al., 2001). Communities within this horizon were mobile (nomadic) pastoralists whose subsistence strategies remain characteristic in the steppe region even today (*14–16*). Within this context, the Pazyryk culture (6th-3rd c. BCE) prevailed in the Mountainous Altai, distinguished by a complex hierarchical system, numerous elite kurgans, and extensive contacts with distant cultural centres of Asia (*14–16*). The Pazyryk culture’s collapse was simultaneous with the military campaigns of the Xiongnu Empire in the 2nd c. BCE, when the Bulan-Koby archaeological culture in the Mountainous Altai emerged (*17–19*). This period has been associated with the destruction of the Scythian-type cultures and the ‘displacement and cultural assimilation’ of their inhabitants (*17–19*). In correlation, the Bulan-Koby exhibits major cultural influences from the Xiongnu and later steppe polities, introducing innovative elements to the Mountainous Altai’s material culture and funerary sphere (*17–19*) (Text S1a).

In the Forest-Steppe Altai, however, the Scythian cultures were succeeded by the yet-unsampled Kulay culture in the 2nd-1st c. BCE (*20*), followed by the rise of the Odintsovo and Medieval Upper Ob cultures around the 2nd-4th c. CE (*21–24*). These cultures introduced novel traits and traditions comparable to the West Siberian taiga belt (boreal forest), and are attributed to early Samoyedic peoples (*20*, *21*, *24*). The Odintsovo culture also incorporated additional influences from other societies of Inner and South-Central Asia (*21*, *23*, *24*) (Text S1b). The Odintsovo culture’s population inhabited the region before the Turkic culture emerged, and persisted on the periphery of the Turkic Khaganate without adopting Turkic cultural customs (*21*, *24*).

The 5th c. CE saw the arrival of the Ashina clan in the Altai area, recorded as consisting of five hundred families (*17*, *25*, *26*) (Text S1c). They united the local Bulan-Koby-associated groups and adopted the name ‘Türk’, and these mark the time when the Turkic material culture first emerged in the Mountainous Altai (*17*, *25*). The primary features of this culture reflect the rise of a new, distinct worldview during the Early Medieval period, yet several of their cultural elements were continuous from the earlier Pazyryk and Bulan-Koby materials and funerary traditions (*17*, *27*). They later founded the Turkic Khaganate in the Eastern Steppe in 551-552 CE and expanded their rule and cultural influence over a vast region (*17*, *25*). The resulting cultural sphere is polymorphic, with regional differences likely shaped by the ‘polyethnic’ groups under the realm (*28*).

The collapse of the Eastern and Western Turkic Khaganates—which had split in 603 CE—led to the rise of new political formations across the steppe during the 8th c. CE (*25*, *29*). During this period, the Turkic-type traditions reached the Forest-Steppe Altai in successive waves, giving rise to the Srostki culture through a mixture of the Turkic and Odintsovo elements (*30–33*) (Text S1d). The region subsequently came under the control of the Turkic-speaking Kimak and Kipchak groups, and the archaeological findings suggest that the Odintsovo-remnants became subordinate to the newly-arriving communities (*30–32*). The Srostki culture persisted until the Mongol Empire’s conquests in the 13th c. CE, when new archaeological entities such as the Karmatsky culture emerged (*30*) (Text S1d). Further details on the archaeological cultures and burials are provided in Supplementary Text S1.

In this study, we aimed to investigate whether the well-documented historical and archaeological transformations in the Altai area between the 4th century BCE and the 12th century CE were reflected in the region’s genetic history. To do this, we constructed a genomic time transect spanning more than a millennium, focusing on key archaeological sites in the region. We explored the genetic compositions and interactions of the ancient inhabitants of the distinct mountain and forest-steppe ecosystems. This approach also enabled us to explore the extent to which genetic continuity, admixture, and population shifts correlated with the spread of novel cultural traditions in the multiethnic Altai setting.

## Results

### Overview of the dataset and the analyses

We collected a total of 180 human remains excavated from 39 burial sites associated with seven archaeological cultures spanning the 6th-4th centuries BCE and the 3rd-14th centuries CE in the Mountainous Altai, Forest-Steppe Altai and the Novosibirsk Upper Ob Plateau (Fig. 1; Table S1; Text S1). The time-table plot of the cultures and the analysis groups can be followed in Figure 1C, and further context on the sampling and these cultures can be found in Box 2. After initial screening of the samples, we generated genome-wide data from 91 individuals through shotgun sequencing and 1,4 million SNP target enrichment (*34*) of these samples, where respectively 510,184 and 395,601 were the average SNP hits of the shotgun sequenced and the target enriched samples on the 1240k SNP panel (*35*)) (Methods, Table S1). For population genetic analyses, we combined the new dataset with previously published ancient and modern pseudo-haploid genotypes from the v54.1 AADR dataset (*36*) and other relevant sample sets (*5*, *10*, *13*, *37*, *38*) (Methods; Table S2; Text S2).

#### Box 2

Sample selection and the definition of cultural groups

Our analyses are based on samples from representative burial sites and inhumed individuals, since cremation burials—also present in the study area—are unsuitable for DNA analysis. We analysed samples from burials in the Altai, assigned to cultures that emerged before the rise of the Early Medieval Turkic culture, alongside those associated with the rise and spread of the Turkic traditions (Turkic and Srostki cultures). We note that we treat these archaeological cultures not as ethnic designations but interpret them with reference to previously described material cultures and funerary rites (Text S1). We refer to the Mountainous Altaian Bulan-Koby, Forest-Steppe Altaian Odintsovo, Novosibirsk Upper Ob groups and the published data from Berel kurgans in Eastern Kazakhstan as ‘non-Turkic’ type cultures. This terminology is used for clarity and brevity, to distinguish between groups with and without Turkic traditions that emerged and spread in the Early Medieval era, and to describe cultural changes across the Altai’s geographical zones at different stages in time.

We first performed Eurasia-wide principal component analysis (PCA) (*39*, *40*) and supervised ADMIXTURE analysis (*41*) (Methods; Fig. 2, S2-S4). We detected genetic outlier individuals on the PC1-PC2 and PC1-PC3 spaces of the PCA as described in the Methods and Supplementary Text S3 (Table S3). We verified our observations with subsequent outgroup-*f*_3_ and *f*_4_/*D* tests (*42*), and used qpAdm (*42*, *43*) to test the fitting deep and proximal ancestry admixture models for the target groups and samples (Fig. 3, S5, S6; Table S4-S6; Text S4). Using the qpAdm models presented in Fig. 3a and Table S6, we inferred pairwise admixture times with DATES (*9*) for our analysis groups (Fig. 3b, Table S7). We analysed runs of homozygosity (ROH) signals and inferred the effective population sizes (*N_e_*) from samples grouped by subsequent periods (*44*) (Fig. S7; Table S8). Through genetic relatedness analyses (*45*, *46*) and identity-by-descent (IBD) analysis of shared autosomal haplotypes (*47*), we aimed to capture the close and distant genetic links among the individuals in our dataset (Methods; Text S5, S6). We report two pedigrees spanning three generations, both consisting of samples from elite burial contexts (Fig. S8; Table S9; Text S5). The IBD dataset prepared with ancIBD (*47*) had 846 ancient individuals, and represented at least 12 cM shared segment per pair indicative of common descent (Methods; Fig. S4, S9, S10; Text S6). We repeated the network-building on a dataset filtered for at least one stretch of 20 cM shared IBD segment to observe stronger genetic links, since such segments identify more recent connections of around five meiosis events (*47*) (Fig. S11; Text S6). Findings from the customary population genetic analyses, as well as objective metrics on the IBD network and detailed analyses and evaluations conducted on it, are presented in Supplementary Texts S2-S6.

**Figure 2:**
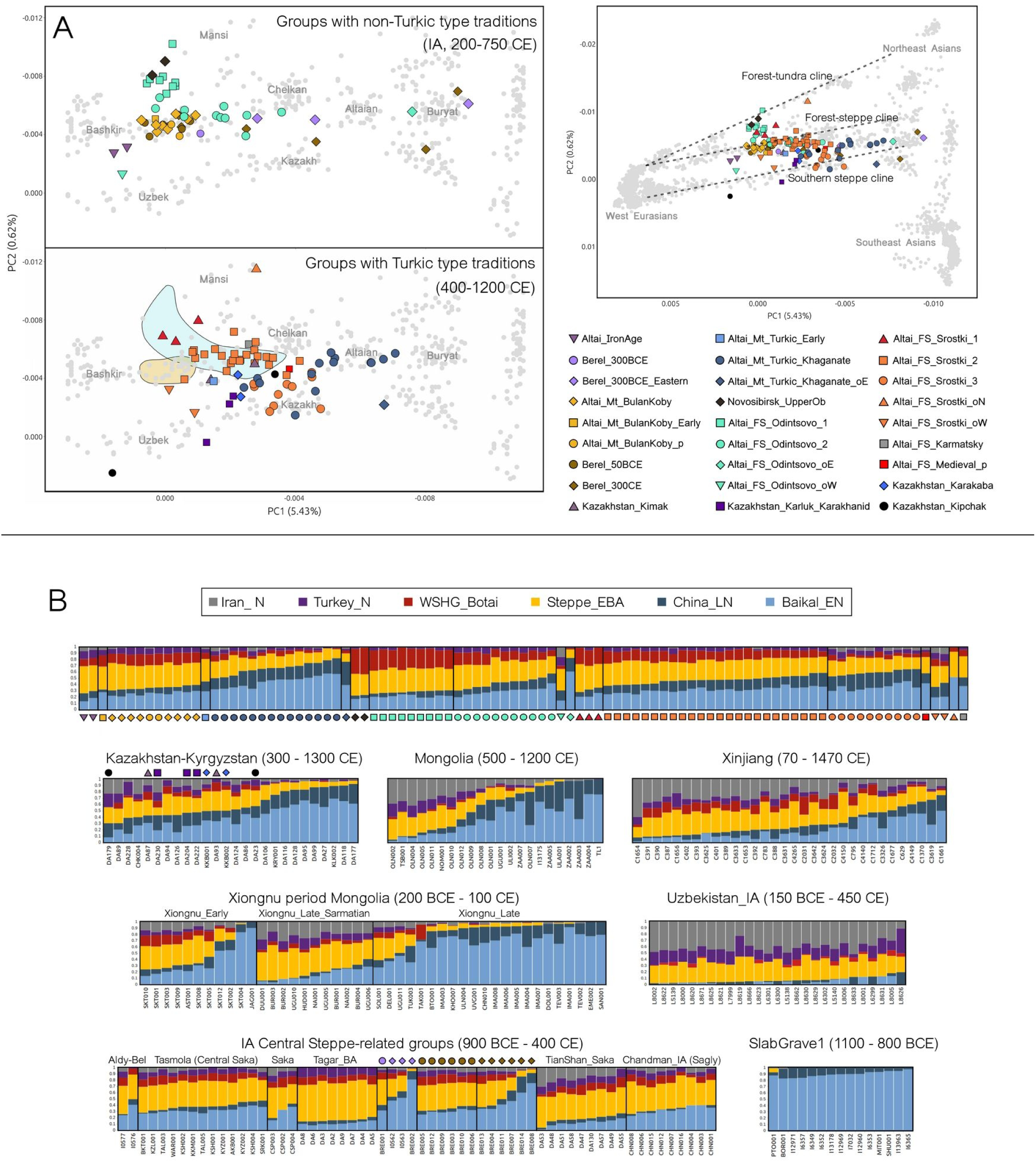
PCA and ADMIXTURE analysis. (A) Principal Component Analysis (PCA) plot of Eurasia on the PC1-PC2 space, based on modern Human Origin dataset (8), including newly reported and published genomes. The right panel shows the complete version of the plot, and the left panels focus on the contextually-important groups from the full Eurasian plot. Ancient genotypes were projected onto the modern genomic variation (gray points). The labels on the PCA plots show broad geographical regions and modern groups. The genetic clines of modern Eurasia are respectively shown in dashed lines on the right panel. (B) K6 supervised ADMIXTURE results of the newly reported genomes, and a subset of published ancient individuals. Inner Asian groups mentioned in the main text are presented here. Individuals per each analysis group are in order according to their increasing total East Asian ancestries (Baikal_EN + China_LN). The symbols in the PCA plot legend also represent the individuals plotted in the upper row of (B).

**Figure 3:**
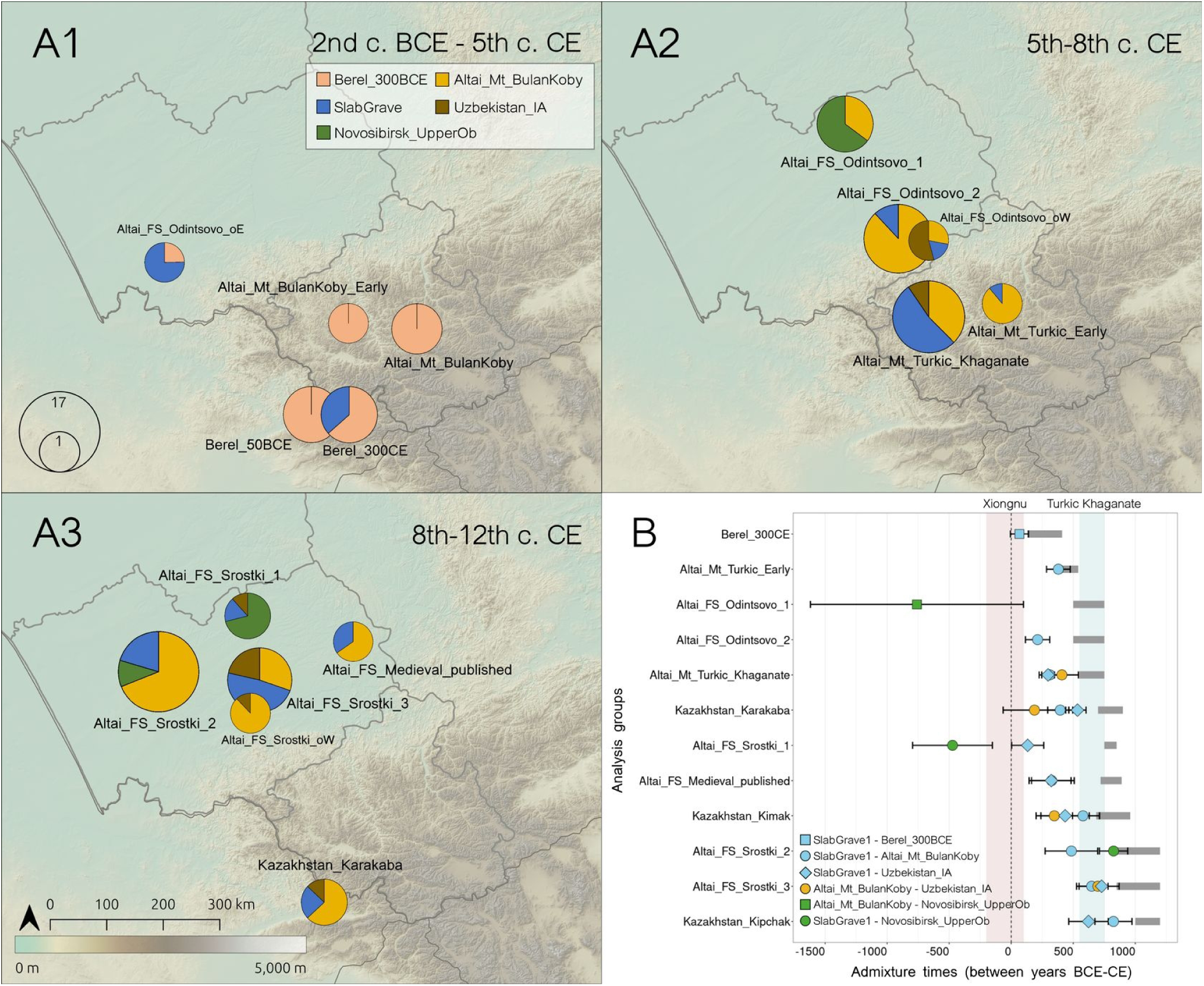
Ancestry modeling and admixture times. (A1-3) Fitting (P≥0.05) lowest rank group-based qpAdm ancestry models of the newly-analysed Altai groups and outliers. The legend shows the designated color for the plotted source qpAdm groups. The right groups used in qpAdm analyses are named in the methods section. The circle sizes indicate the total individual number at a given analysis group. (B) Admixture times of the main analysis groups using DATES. The horizontal black bars show the mean admixture times ± standard errors (considering 29 years per generation), and the mean values are in coloured symbols which are based on the used source combinations (see legend). The horizontal gray bars show the archaeologically assigned dates of the analysis group. The results from DATES can be found in Supplementary Table S9.

### Initial observations from the Altai transect

Across two- and three-dimensional PCA spaces, the Altai inhabitants from both the Iron Age and the non-Turkic type Migration Period and Medieval cultures primarily align with the modern forest-tundra (taiga biome) and forest-steppe –or steppe-forest– genetic diversities of the Asian continent, whereas the Turkic type groups plot closer to the modern southern steppe genetic variation (terminology following Jeong et al., 2019 (*8*)) (Fig. 2a, S2, S3). These differences are evident in ADMIXTURE as well, which displays multiple genetic profiles and varying contributions from the ancestral source populations (Fig. 2b, S4). According to these, some Forest-Steppe Altai and Berel individuals from the same cultural context and period are grouped into multiple, genetically distinguished analysis groups at each phase of the time transect. We label these with numbers or suffixes in the following sections (*e.g.* Altai_FS_Odintsovo and Altai_FS_Srostki analysis groups 1 and 2). We detect such coherent groups and additional PCA outliers in the Odintsovo, Turkic, and Srostki contexts as well (see Methods), with further details provided in Supplementary Text S3 and Supplementary Table S3. Our primary observations are further supported by outgroup-*f*_3_ and *f*_4_/*D*-statistics, where relevant findings are detailed in Supplementary Text S4, and listed in Supplementary Tables S4 and S5.

Our dataset includes two new Altai genomes dated to 6th-4th c. BCE, along with previously published 4th-2nd c. BCE genomes from the Berel kurgans in Eastern Kazakh Altai (*2*, *48*) (Table S1, S2). They are collectively associated with the IA Eastern Scythian cultures, particularly the Pazyryk culture of the Mountainous and Eastern Kazakh Altai (Text S1, Text S2). We observe a heterogeneous genetic profile spanning East-West Eurasian ancestries, and discuss these differences below. This heterogeneity ceases after the 2nd c. BCE, but reappears in the 2nd-4th c. CE at Berel, with individuals showing elevated East Asian affinities (Fig. 2, S2-S4, S6; Table S5; Text S4). The newly reported 6th–4th century BCE Altaians differ from a Berel individual of the 4th–2nd c. BCE, whose genetic profile, in turn, resembles that of the 2nd c. BCE – 1st c. CE and later 3rd–5th c. CE individuals from the Mountainous Altai (Bulan-Koby context) (Fig. 2, S2–S6; Table S5; Text S4). After the 5th c. CE, the Turkic culture emerges in the region, coinciding with a stronger eastern genetic shift in the Mountainous Altai (Fig. 2, S2-S6; Table S4, S5; Text S1c, S4). Meanwhile, contemporaneous Forest-Steppe Altaians associated with the Odintsovo culture (4th-8th c. CE) show more varying ancestries, where we detect at least two genetically distinct groups. Odintsovo group 1, similarly with the Novosibirsk Upper Ob group, shows high levels of allele sharing with the Neolithic-Eneolithic ANE-related peoples (Box 1; Fig. 2, S2-S4, S6; Table S5; Text S4). Odintsovo group 2, however, is distributed along a cline between Odintsovo group 1 and East Asian genomes (Fig. 2, S2-S6; Table S4, S5; Text S4). The 8th-12th c. CE Forest-Steppe Altaians (Srostki groups 1 and 2) mainly overlap with the genetic diversity seen in the earlier regional Odintsovo period (Fig. 2, S2-S6; Table S4, S5; Text S4). Srostki group 1 represents the early stage (750-850 CE), includes individuals from the Inya-1 burial site in the northern Forest-Steppe Altai (Fig. 1; Text S1d), and carries the elevated ANE-related genetic components further in time. Srostki group 2, on the other hand overlaps with the variety of Odintsovo group 2, unlike Srostki 1. These two Srostki groups, therefore suggest continuity from the earlier 4th-8th c. CE Forest-Steppe Altai. We also distinguish a third genetic group (Srostki group 3, also from the classic Srostki stage) which has no detectable precursor in the Odintsovo period (Fig. 2, S2-S6; Table S4, S5; Text S4). These regionally-continuous and incoming varieties occur side-by-side at Srostki-period burial sites (Fig. 1; Table S1). We describe the Y-chromosomal haplogroups and their relationships to prehistoric and Medieval populations in Supplementary Text S7.

In line with the observed continuity of regional ancestries, we find high ratios of IBD sharing between the groups (modules) within each geographical region, depicting biological continuity between succeeding groups within the respective mountainous and forest-steppe areas (with the exception of Altai_FS_Srostki_3) (Fig. 4, S11; Table S10, S11; Text S6). This is evidenced in the high-fraction of ≥12 cM links between the modules in these territories, while the same network also shows a connectedness of the ancient Altaians to some other groups around the study region (Fig. 4; Text S6). Most of the newly-analysed groups belong to a Altai IBD cluster (cluster 0_0_0), which is the largest subcluster in the Leiden analyses of the IBD network, corresponding to our densely-sampled regional dataset. This cluster in turn is a part of the post-IA Eurasian Steppe cluster (cluster 0) (Fig. S9, S10; Table S10; Text S6). The observed ≥20 cM IBD links also support these findings, as such strong links rarely occur between unrelated groups (Fig. S11; Text S6). Additional interpretation of the IBD results and related network metrics is provided below and in Supplementary Text S6.

**Figure 4:**
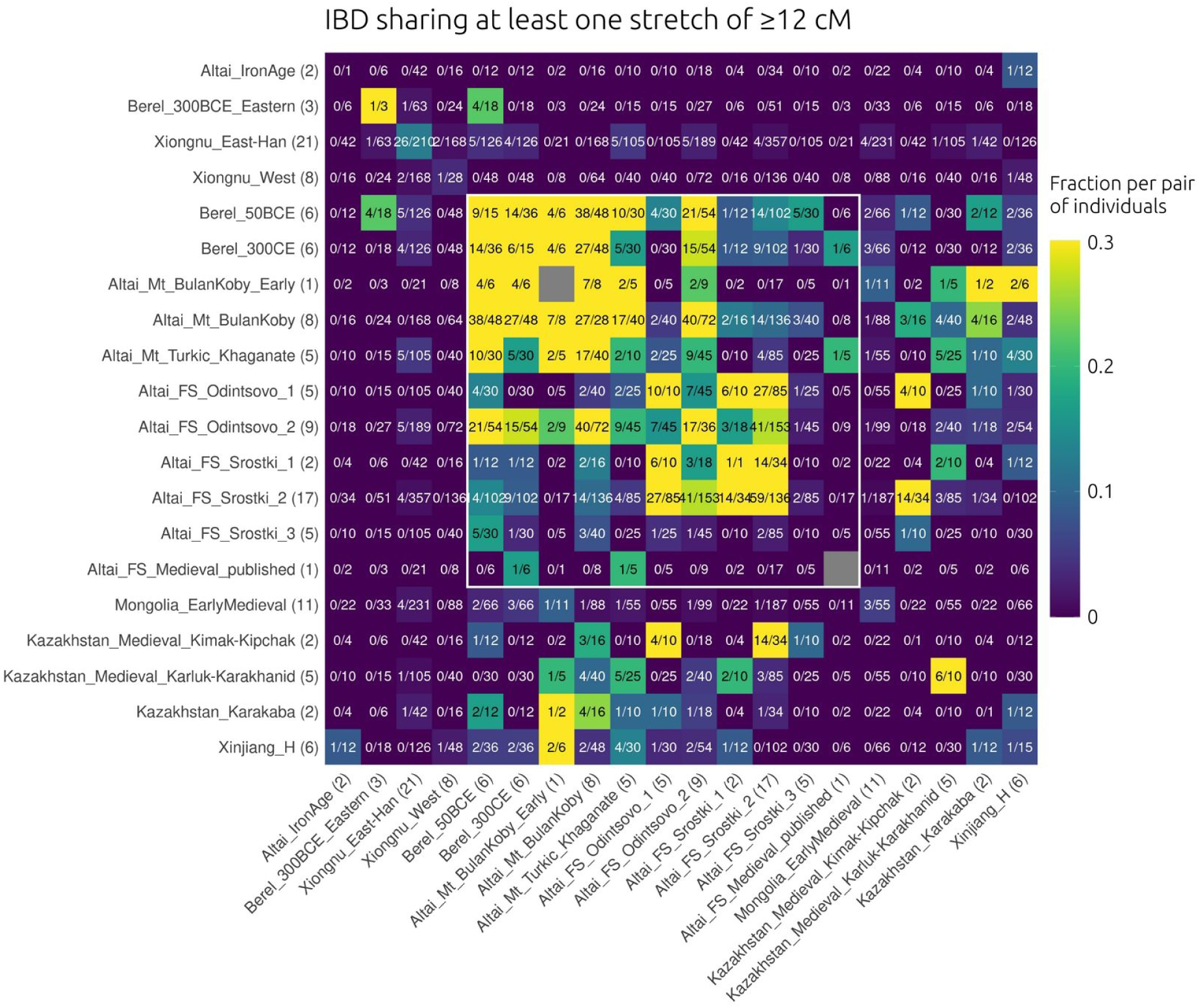
IBD heatmap of the newly-analysed Altaians and other contextually-related groups. IBD sharing between chosen analysis groups in the 12 cM IBD dataset. Parentheses next to the group labels indicate the group sizes. The numbers in the cells represent (total detected IBD links)/(total possible pairwise links) and the colouring is based on normalization with the same fraction similar to Ringbauer et al., 2024 (47). A higher fraction means closer connectedness between the analysis groups. Groups of the post-IA Altai context are framed in white. The IBD analysis groups differ from the customary population genetic analysis groups (see Methods, Table S10).

### Genetic variety of the Iron Age Altai populations

We first establish the genetic profile of the two earliest individuals in our new dataset (Altai_IronAge, 6th-4th c. BCE), who are associated with the Early IA –Scythian– period in the Altai (Fig. 1; Table S1; Text S1). We also incorporate previously published individuals from the Eastern Kazakh Altai’s Berel kurgans into the dataset, dated to this period and ascribed to the Pazyryk horizon (Fig. 1). These are labeled as Berel_300BCE (*2*) and Berel_300BCE_Eastern (*2*, *48*) (Table S2; Text S2). Their genetic composition, based on PCA, ADMIXTURE and *f*_4_/*D* tests, falls into a variety between the East-West Eurasian ancestries (Fig. 2, S2-S6; Table S5; Text S4). Altai_IronAge and Berel_300BCE align with the gene pool of other IA groups largely associated with the Eastern Scythian - Saka cultures (IA Central Steppe) (*2*), whereas Berel_300BCE_Eastern is significantly more shifted towards the East Asian ancestries (Fig. 2b, S4, S6; Table S5; Text S4).

In qpAdm, Berel_300BCE can be modeled with both Early BA Cis-Baikalians or the Late BA Mongolian Khövsgöl (51-55%), with additional Middle Late BA Central Steppe (39-41%) and BA Gonur ancestries (6-8%) (Table S6a). The composition of Berel_300BCE is therefore similar to other groups of the IA Central Steppe, and is cladal with the 8th-6th c. BCE Tasmola group (*2*) from the Kazakh Central Steppe in proximal qpAdm models (Table S6b). In contrast, the newly sampled 6th-4th c. BCE Iron Age Altaians differ in deep and proximal qpAdm models (Table S6a, S6b). This group has a tendency towards Southwest and Southeast Asian ancestries in ADMIXTURE, differing from Berel_300BCE (Fig. 2b). Therefore, although Altai_IronAge shows a profile comparable to Berel_300BCE and other contemporaneous IA Central Steppe groups, it is not cladal with any of those in qpAdm, and needs additional components to represent more southern ancestries (Table S6b). We also show that Berel_300BCE_Eastern requires the BA Ulaanzuukh (*3*) or IA SlabGrave1 (*5*) sources of the Eastern Steppe in most plausible models unlike other groups, a region which could be the source of such an eastern shift in this group when considered together with the other analyses (Fig. S6; Table S5, S6a; Text S4). Consequently, we detect regional differentiation in the Altai during the Pazyryk period (6th-3rd c. BCE), even among sites <200 km apart, coinciding with the comparatively large effective population sizes (*N_e_*) estimated for this interval—the highest in our dataset (Fig. 1; Table S8). In IBD analyses we find that the newly sampled 6th-4th c. BCE (Altai_IronAge) individuals do not have any detectable links with the later Altaian main analysis groups, which, together with the rest of the results, suggest that this population did not survive the 2nd c. BCE Xiongnu expansion in the region (Fig. 4, S11; Table S10, S11). Conversely, the 300 BCE Berel individuals have detectable ≥12 cM links to three individuals of Berel_50BCE (*2*), a later group described below (Fig. 4; Table S10, S11).

### Population dynamics of the Mountainous Altai and post-Iron Age East Asian influxes

The Bulan-Koby culture burial sites of the Mountainous Altai are contemporaries and adjacents of the Eastern Kazakh Altai Berel site’s post-Pazyryk burials (Fig. 1). Contrary to the Pazyryk period, Bulan-Koby-associated individuals from the Xiongnu period (Altai_Mt_BulanKoby_Early of the 1^st^ c. BCE (*49*)) and post-Xiongnu period (Altai_Mt_BulanKoby presented here, 3rd-5th c. CE) show ancestries akin to the Xiongnu-period Berel group (Berel_50BCE (*2*)), and to two out of six individuals of the post-Xiongnu era Berel group of 2nd-4th c. CE (Berel_300CE (*2*), different from Berel_300BCE) (Fig. 2, S2). Correspondingly, this profile is different from the newly sampled 6th-4th c. BCE Iron Age Altaians, yet similar to the ancestry of the Pazyryk-period Berel_300BCE, as seen in PCA, ADMIXTURE and *f*_4_/*D* results (Fig. 2, S2-S6; Table S5; Text S4). These are also reflected in qpAdm, where the Bulan-Koby and Berel_50BCE ancestries are best explained with Berel_300BCE as a single source (Fig. 3; Table S6c). Overall, the Bulan-Koby and Berel groups of the Xiongnu era display high rates of allele sharing with other IA Central Steppe profile groups (Table S6c). However, in the qpAdm admixture models, the newly sampled 6th-4th c. BCE Iron Age Altaians as a source can only explain 64-74% of the Bulan-Koby groups’ ancestry, and require additional genetic contributions from other groups (Table S6c). We therefore interpret that the post-Pazyryk Altaians are continuous from a group related to Berel_300BCE, surviving up to the 5th century CE in the Mountainous Altai. Unlike Mountainous Altai, the 2nd-4th c. CE Berel in the Kazakh Altai has an extra East Asian ancestry contribution (36.6±2.2%), whose admixture dates indicate the Xiongnu period for the East Asian influx (Fig. 3b; Table S6c, S7). This observed heterogeneity is similar to the previous Pazyryk period, when an earlier East Asian influx is recorded in the Berel region but not in the Mountainous Altai, yet more sampling is needed from the latter.

We observe an increase in shorter segments of ROH and a corresponding *N_e_* decrease in the Mountainous Altai and Eastern Kazakh Altai region through the 1st c. BCE – 5th c. CE (Fig S7; Table S8). IBD analyses provide further evidence for possible population declines, as in both 12 cM and 20 cM datasets, the populations of the 1st c. BCE – 5th c. CE Mountainous Altai and Berel are interconnected, and collectively form a biological group with strong IBD connections as explained in detail in Supplementary Text S6 (Fig 4, S11). This biological group also has links to the post-5th c. CE groups in the Mountainous Altai, as well as to the adjacent areas, indicating them also as a source population with detectable biological continuity in the region (Fig. 4, S11; Text S6).

The *N_e_* increases and the ROH signals diminish again in the Mountainous Altai after the 5th c. CE, coinciding with the significant archaeological and historical changes in the region (*17*, *25*, *26*), as well as with our genetic observations, suggesting an increase in the population of the region (Fig S7; Table S8). We present the first genome from an individual associated with the Early Turkic culture, recovered from a burial of low social status, and radiocarbon dated to 430-580 CE (95.4% CI), providing the earliest known evidence for the Turkic-period genetic transitions in Inner Asia (Fig. 1; Table S1; Text S1c). During and after the 5th c. CE, the Mountainous Altaians experience a deviation towards the East Asian genetic variety as observed on the PCA, and the individuals show between 45-80% East Asian ancestry in ADMIXTURE (Fig. 2, S2-S4). The *f*_4_/*D* analyses confirm an influx from a likely BA Eastern Steppe-related group, similar to which contributed to the Berel groups in the Pazyryk period and 2nd-4th c. CE (Fig. S6; Table S5; Text S4). The Turkic-period groups of Mountainous Altai (labelled with Altai_Mt_Turkic prefix) both have increased Eastern Steppe-related ancestry in the fitted qpAdm models (11.1±3.4% and 52.9±1.6% respectively), in addition to their IA Central Steppe component (Fig. 3a, Table S6c). This profile persists in the region up to the 8th-9th centuries CE, as we detect it in the Karakaba group of the Eastern Kazakh Altai (*2*) (Fig. 2, 3, S2-S6; Table S5, S6; Text S2). While the East Asian gene flow is similar to our observations for 2nd-4th c. CE Berel, we interpret this as having happened in another wave from the Eastern Steppe and not from Eastern Kazakhstan, where the Berel site is located (Fig. 1). This is due to the differences in the admixture dates, which correspond to the post-Xiongnu era for the Turkic-period Mountainous Altaians, and the scarcity of the IBD links between the Turkic-period group and the 2nd-4th c. CE Berel (Fig. 3b, 4, S11; Table S7, S10, S11; Text S6).

The Turkic-period Mountainous Altaians exhibit significantly higher IBD sharing with the 3rd-5th c. CE (late) Bulan-Koby dataset in shorter (≥12 cM) segments than with 2nd-4th c. CE Berel (Fisher’s exact test, p = 0.036, odds ratio= 3.63, 95% CI≈ 1.06–14.68) (Fig. 4; Table S10-S12; Text S6). In addition, longer (≥20 cM) segment sharing is present between the Turkic-period group and the late Bulan-Koby (three detected links), but not with 2nd-4th c. CE Berel (Fig. S11; Table S11; Text S6). These findings support a substantial contribution from the local Mountainous Altai baseline population to the formation of the Turkic-period Altai population. Notably, the ≥12 cM IBD links within the Turkic-period Mountainous Altaians are fewer than the inner links of the late Bulan-Koby and Berel_50BCE each, which is statistically significant against the late Bulan-Koby, and observed together with decreased ROH and increased *N_e_*in the area (Fisher’s exact test, p= 7.791e-06, odds ratio= 0.012, 95% CI≈ 0.00–0.15 and p= 0.099, odds ratio= 0.18, 95% CI≈ 0.014–1.36) (Fig. 4; Table S10-S12).

### Traces of Ancient North Eurasian genetic ancestry in the Forest-Steppe Altai and a post-Turkic Khaganate East Asian influx

The 4th-8th c. CE Novosibirsk Upper Ob (Medieval Upper Ob culture) and Forest-Steppe Altai (Odintsovo culture) groups carry some genetic components notably different from their contemporaries in the Mountainous Altai and Inner Asia (Fig. 2, S2-S4). A distinctive feature of these groups is the elevated rates of allele sharing with WSHG_Botai in ADMIXTURE (up to ∼40%) (Fig. 2b). On the three-dimensional PCA space, the Upper Ob group and Odintsovo group 1 also plot closely with WSHG_Botai, but not with Altai_HG, suggesting a difference in ANE-related sources of the North Eurasian hunter-gatherer cline (Box 1; Fig. S3). *f_4_*/*D*-statistics verify their shared alleles with WSHG_Botai, instead of Altai_HG or other ancient Siberian groups, showing an exceptional phenomenon of groups as late as the Medieval period expressing this ancestry, the first case in published and available post-BA datasets to our knowledge (Fig. S6.3; Table S5; Text S4). Outgroup-*f*_3_ statistics also indicate Northern Eurasian affinities for these groups, especially with the modern Samoyedic Selkups and Yeniseian Kets (Fig. S5; Table S4; Text S4).

In qpAdm, the Upper Ob group can be modeled with 26-42% contribution from WSHG_Botai in 3-way mixture models (p≥0.05), where it still requires 26.3±5.8% of this source when the ANE-rich Early BA Cis-Baikalians are in the sources (Table S6a). The Upper Ob group is the only available candidate to represent the ANE-related ancestry in the 4th-8th c. CE Forest-Steppe Altai in the proximal qpAdm models, since possible IA precursors such as the Kulay culture’s population remain unsampled (Fig. 3, Table S6d). Odintsovo group 1 necessitates the Upper Ob source in qpAdm (64.8±5.1%), a feature present across all feasible models of this group, with additional Bulan-Koby-related ancestry (35.2±5.1%) in the regional model (see Methods, Fig. 3, Table S6d). Contrarily, two-way models for Odintsovo group 2 do not require Upper Ob, but fit with Bulan-Koby-period Mountainous Altaians (84-88%) + the Eastern Steppe sources (12-16%) (Fig. 3, Table S6d). The individuals of Odintsovo group 1 come from the more northern parts of the Forest-Steppe Altai, near the Novosibirsk Ob region, whereas every individual of Odintsovo group 2 originates from the Gorny-10 burial site, located in an intermediate zone between the Forest-Steppe and Mountainous Altai (Fig. 1; Table S1; Text S1b). This reflects notable genetic variability in the Odintsovo culture’s people and coincides with landscape differences (Fig. 2, 3a).

The admixture times of the two Odintsovo groups also differ, with the dates for group 1 being earlier, whereas group 2 aligns with a later formation (Fig. 3b). Notably, these groups have varying, but converging *N_e_* and IBD patterns. Odintsovo 1 individuals have ROH signals in both short and long segments, showing both consanguinity and the smallest *N_e_* in the Altai (Fig. S7; Table S8). This is accompanied by dense IBD links, with all five Odintsovo group 1 individuals in the IBD dataset sharing ≥12 cM links to each other, and also showing elevated ≥20 cM within-group segment sharing fractions compared to most other groups of the IBD dataset (Fig. 4, S11; Table S10, S11). On the other hand, Odintsovo group 2 has three times higher *N_e_* compared to Odintsovo 1 and does not have signals for parental relatedness (Fig. S7; Table S8). While group 2 also exhibits a high fraction of within-group IBD links, the proportions of these are significantly lower than that of group 1 (Fisher’s exact test, p= 0.0027, odds ratio= 0, 95% CI≈ 0–0.49) (Fig. 4; Table S10-S12). Between groups 1 and 2, we detect even lower rates of IBD sharing than within-group links, suggesting two communities with limited interactions with each other in different areas of the Forest-Steppe Altai (Fisher’s exact test for each group, respectively p= 6.649e-07, odds ratio= 0, 95% CI≈ 0–0.11 and p= 0.003, odds ratio= 0.21, 95% CI≈ 0.06–0.64) (Fig. 4; Table S10-S12; Text S6). We acknowledge that a wider sampling from the area may change the observations. Yet, the Odintsovo group 2 from Gorny-10 exhibits more than half of the IBD links between the Mountainous and Forest-Steppe Altai regions as an intermediary group. Notably, their links are stronger with the pre-6th c. CE late Bulan-Koby group than with the 6th-8th c. CE Turkic period in the Mountainous Altai in both 12 cM and 20 cM datasets (Fisher’s exact test for 12 cM, p= 0.0002, odds ratio= 4.93, 95% CI≈ 1.97–13.42) (Fig. 4, S11; Table S10-S12; Text S6). This being noted, the 4th-8th c. CE Forest-Steppe Altai mating pool is altogether smaller than that of the 5th-8th c. CE Mountainous Altai region (Table S8).

In the 8th-12th c. CE Forest-Steppe Altai, we considered the Srostki period in its early and classic stages. Within this period, we describe three genetic groups (Altai_FS_Srostki groups 1, 2 and 3). Srostki groups 1 and 2 overlap with the sampled genetic variety of the previous Odintsovo period in the Forest-Steppe Altai as observed in the PCA and ADMIXTURE, while group 3 is different (Fig. 2, S2-S4). Groups 1 and 2 also exhibit comparable tendencies to their regional precursors in the outgroup-*f*_3_ and *f*_4_/*D* tests, and the regional continuity is backed up by the group-based qpAdm analyses (Fig. 3, S5, S6; Table S4-S6; Text S4). However, using the Odintsovo-period groups as regional sources, we also observe highly variable additional East Asian ancestry in both Srostki-period groups (Table S6d). The outgroup-*f*_3_, *f*_4_/*D* and fitting qpAdm results for Srostki group 3 differ significantly from those for the Odintsovo-period groups, also from Srostki 1 and 2, and demonstrate a profile close to the 5th-8th c. CE Mountainous Altaians’ composition with an extra Southwest Asian component (Fig. 3, S5, S6; Table S4-S6; Text S4). The admixture times also vary; for Srostki group 1, the dates overlap with the results for Odintsovo group 1, whereas for Srostki groups 2 and 3, a recent admixture of East Asian ancestry with the local profile is estimated (Fig. 3b; Table S7).

The *N_e_* of Srostki groups 1 and 2 show minor increases, and Srostki group 2 also exhibits ROH signals in longer segments, which are in line with a major regional continuity of the Odintsovo period groups (Fig S7; Table S8). The combined *N_e_* in the forest-steppe Altai however increases with the newly appeared Srostki group 3, where ROH signals are absent (Fig S7; Table S8). The IBD links of the Srostki-period groups reflect the presented genetic ancestries. Srostki group 1 has a higher rate of ≥12 cM links to Odintsovo group 1 –with whom they overlap in genetic ancestries– than to Odintsovo group 2 (Fisher’s exact test, p= 0.035, odds ratio= 6.86, 95% CI≈ 0.97–64.3), while Srostki group 2 has high fractions of ≥12 cM links with each Odintsovo-period group without any statistical difference, suggesting a continuity from both (Fisher’s exact test, p= 0.46, odds ratio= 1.27, 95% CI≈ 0.68–2.36) (Fig. 4; Table S10, S11; Text S6). In the case of longer segment (≥20 cM) sharing, however, the Srostki-period groups have stronger links to their respective Odintsovo-period groups, aligning with the broader separation of the ancestries in the previous era (Fig. 3, S11; Text S6). Srostki group 2 also has longer segment sharing with the contemporaneous Kimak-Kipchak individuals at a rate that is not present in the Odintsovo period or even in other Srostki-period groups, indicating higher levels of interaction between them (Fig. S11; Table S10, S11).

## Discussion

Here we present 91 ancient human genomes dated between the 6th c. BCE and 14th c. CE from the Mountainous Altai, Forest-Steppe Altai and Novosibirsk Upper Ob Plateau in Southern Siberia (Fig. 1; Table S1; Text S1). Using a combination of ancient DNA-adapted population genetic methods, we find evidence that East Asian genetic influences characterised population shifts within the Altaian time transect (Fig. 2, 3; Table S4-S7; Text S4). These genetic events are accompanied by an increase in effective population sizes, potentially indicating external contacts (Table S8). While both the Mountainous and Forest-Steppe Altai populations are affected by these, our analyses indicate a relative genetic differentiation between these groups aligning with the landscape differences of the region (Fig. 2, 3; Table S4-S7; Text S4). This is observed primarily during the 5th-8th c. CE, and ceases to exist with new population movements to the Forest-Steppe Altai region thereafter (Fig. 3; Table S6, S7). Yet, the patterns of homozygosity (ROH) remain distinct between the regions, which could indicate enduring cultural differences (Fig. S7).

Our two new genomes from the Iron Age represent the IA Eastern Scythian (Saka) cultures in the Altai, one of them associated with the Pazyryk culture of this Saka horizon (Fig. 1; Table S1). Their ancestries are similar to other IA inhabitants of the Central Steppe and Western Mongolia, but they have slightly higher Southwest and Southeast Asian genetic components, suggesting additional interactions not seen in other published Eastern Scythian-associated groups (Fig. 2, 3, S2, S4). This niche genetic profile is one of several present around the Altai during the IA and appears to have disappeared after the 2nd c. BCE (Fig. 2-4, S2-S6; Table S4-S6). In contrast, other local variants persist as seen for Berel_300BCE, whose similar but distinguishable IA Central Steppe-related ancestry is continuous in the area. These patterns suggest that the Berel groups may have prevailed over other communities at that time. Having the highest *N_e_* in our dataset (Table S8), the IA-period Altai-Berel peoples’ genetic heterogeneity suggests distant interactions, which is considered to be a characteristic feature of the Pazyryk culture (*15*, *16*). A possible route could be through the Inner Asian Mountain Corridor that served as a facilitator of human contact between the Altai region and more southern areas such as Xinjiang (*50*) (Fig. 2b, S4). In particular, the downfall of the Scythian cultures in the Altai (∼2nd c. BCE) may have resulted in a power and population shift in the region (*17–19*), and coincides with a decrease in effective population sizes as inferred in our analyses (Table S8). Nevertheless, the IA Central Steppe component remained dominant in the Altai region between the 2nd c. BCE – 5th c. CE as observed in the Bulan-Koby culture’s population.

The Turkic cultural traditions emerged around the Mountainous Altai with the blending of local Bulan-Koby cultural and novel practices (*17*, *26*) (Text S1c). The rise and spread of the Turkic cultural traditions in the 5th-8th c. CE were significant for the demise of other Inner Asian cultures, such as the Bulan-Koby itself, and eventually the Odintsovo in the Forest-Steppe Altai, where the new traditions spread and gave rise to the Srostki culture (*30–33*). These cultural transitions coincide with a significant increase of East Asian ancestry in the regions mentioned, while some groups associated with non-Turkic traditions (2-4th c. CE Berel and 6th-8th c. CE Odintsovo group 2) also show detectable East Asian ancestral contributions (Box 2; Fig. 2, 3; Table S4-S6; Text S4). Nevertheless, the shared longer IBD segments of 2nd-4th CE Berel are only sporadic with both Odintsovo group 2 and the Turkic type groups (Fig. 4, S11; Table S10, S11; Text S6). The inferred admixture times of the latter also correspond to later periods than found in 2nd-4th CE Berel, suggesting that there may have been at least two waves of East Asian influx in each period separately, despite originating from a shared BA-IA Eastern Steppe base population (Fig. 3b, S6; Table S4, S5, S7; Text S4). Therefore, according to our findings, the detected East Asian influx from Turkic-period individuals –associated with low-status burials– in the Altai refutes an *elite dominance* scenario (*1*, *51*, *52*) for the spread of the Turkic culture post-5th c. CE (Table S1; Text S1). Instead, it coincides with the appearance of larger groups with predominant East Asian ancestries. This is supported by minimal signs of homozygous runs (ROH) and increased effective population sizes in Turkic-type groups that exhibit the detectable non-local ancestries; mediated probably by an increasing population size, connection system or certain marriage practices, such as exogamy (*44*), which is seen also in other steppe-related communities (*53*, *54*), as well as other factors (Fig. S7; Table S8). The expansion observed for the Turkic-era East Asian signal likely reflects several interacting elements in Inner Asia. For instance, the transition from the Bulan-Koby culture and the Rouran Khaganate to the Turkic culture and the First Turkic Khaganate occurred during a period of Eurasian cooling that climate studies associated with subsistence disruption, famine, and depopulation in the Altai (*55*, *56*), which is consistent with our findings. Meanwhile, the strengthening of the Turkic Khaganate may have facilitated the spread of the politically advantageous groups in the realm. Whereas previous genetic studies reported continuous but varying levels of westward Eurasian spread of East Asian ancestry from the IA to the Medieval era (*1*, *2*, *51*, *52*, *57*), our findings provide finer resolution for this process during the Medieval Period.

Some post-IA groups in the Altai and the Novosibirsk Upper Ob Plateau have additional genetic components, such as the ANE-related ancestry. We show that a population with elevated relations to the Neolithic-Eneolithic ANE-related groups survived in the Siberian mosaic of peoples up until the 9th c. CE in the Forest-Steppe Altai (Fig. 2, 3; Table S5, S6; Text S4). Three newly-presented groups (Novosibirsk_UpperOb, Altai_FS_Odintsovo_1, Altai_FS_Srostki_1) bear significant ANE-related genetic elements, exhibiting a profile closely related to WSHG_Botai, rather than to the Neolithic Altai hunter-gatherers (Altai_HG), who represent the North Eurasian hunter gatherer (NEAHG)-related ancient inhabitants of the Forest-Steppe Altai (*10*, *12*), or than to the BA Okunevo culture’s inhabitants from the Minusinsk Basin in close proximity (*7*) (Box 1; Fig. S6.3; Table S5; Text S4). To our knowledge, this represents the first post-Bronze Age evidence for elevated ANE-related ancestry in the Northern Asian region. Moreover, the Upper Ob and Odintsovo group 1, and their Early Srostki descendants share the most drift with the modern Samoyedic Selkups and Yeniseian Kets among present-day Siberian populations, who inhabit more northern areas and also have affinities with ANE-related reference groups (*8*, *58*, *59*) (Fig. S5; Table S4; Text S4). These findings are consistent with the archaeological links between the Upper Ob and Odintsovo cultures and their connections to the Siberian taiga belt (*21–24*), and the newly analysed samples provide a missing link between the ancient and modern indigenous Siberian populations. This could be further assessed through sampling of the inhabitants of earlier and contemporaneous archaeological cultures in Western Siberia.

Demonstrating the distinct and varying genetic ancestries of the non-Turkic type groups, and their contributions to the Turkic-type cultures’ populations, our findings exemplify substrate groups in the formation of the latter (*28*) (Box 2; Fig. 1-4; Table S5, S6, S10, S11; Text S4, S6, S7). These conform with the previously described affinities between the IA Central Steppe inhabitants and modern Inner Asians, who are mostly Turkic speakers (*48*), as well as with the relations of other distinct groups, such as the Samoyedic- and Yeniseian-speaking peoples who were assimilated and admixed into these modern Inner Asian communities (*12*). However, further research is needed to better understand the Medieval groups’ connections to these modern people. For instance, a frequently-discovered feature of the Early Medieval Mountainous Altaians and Inner Asians is the Y-chromosomal haplogroup J-M172 (J2)—whose sublineage J-PH1795 (PH358) we discuss in Supplementary Text S7—which is still common among some modern Inner Asian groups such as the Uzbeks and Uyghurs (*60*, *61*), yet is reported rare among the modern Altaian populations (0-4%), indicating post-Medieval population events in the studied area with the increase of other paternal lineages such as R, C and Q (*62–65*) (Fig. S12; Table S1).

In conclusion, we present the first large-scale ancient sample set from the Altai region, spanning the Iron Age and Early Medieval periods. We identify a new Iron Age Central Steppe-related group whose genetic legacy vanishes after the 2nd c. BCE, coincident with archaeological and political transformations in the area. Our findings related to the end of the Pazyryk, and the Bulan-Koby periods could shed light on some complex dynamics between the tribally-organised groups and expanding political entities during this period. We also report a unique and previously undescribed genetic profile in the Medieval Upper Ob Plateau and the Forest-Steppe Altai region, which is closely associated with the ANE-related profile of WSHG_Botai in Northern Asia. We provide further evidence that the archaeological cultures do not represent concrete biological groups (*54*, *57*).

However, we also observe that the archaeological developments in the Altai occurred along gene flows, and did not result solely from local advancements or cultural contacts. Our findings support fourfold transformations in the Altai including drastic political changes, archaeological novelties, and genetic events that coincide with climatic events. Nevertheless, regional genetic components also persist in the Altai, carried further by groups present at each stage of transition. Overall, through qpAdm analyses and IBD networks, we demonstrate ∼1400 years of biological continuity despite the genetic shifts among successive pre-Medieval and Early Medieval Altaians.

## Acknowledgments

We are thankful for the computational possibilities the HUN-REN Cloud (https://science-cloud.hu/) provided, which was a key factor to complete resource-intensive processes. We thank Zsófia Rácz, Choongwon Jeong and Juhyeon Lee for their feedback on the research development in the early stage of the study. We thank Leonid Vyazov for his constructive comments at the ending stage of the study. We thank Taylor Mateyka for the proofreading of the Main and Supplementary Texts, as well as the translation of the supplementary archaeological documents. We thank István Zimonyi for his comments on the contexts of culture, ethnicity and language.

## Author Contributions

Conceptualization: AT, AAT, ASN;

Methodology: YCO, BG, ASN;

Collecting the samples: TS, BGM, AT;

Performed the DNA extractions and library preparation: BH, MM;

Conducted the bioinformatics analyses: YCO, KJ;

Performed the statistical analyses and data interpretation: YCO, BG, IM, ASN;

Resources: NS, VVG, SG, PKD, MAD, KYK, YTM, NFS, SST, AVF, MPR, AAT;

Visualization: YCO;

Writing – original main draft of the paper: YCO, BG, NS, AAT, ASN;

Writing – review & editing: YCO, BG, TS, IM, AT, ASN;

Funding acquisition: BGM, ASN; Supervision: AAT, ASN.

## Ethics Declarations

The authors declare no conflicts of interest. The authors had requested and acquired permission from the stakeholders (Altai Museum of Archeology and Ethnography of Altai State University [Музея археологии и этнографии Алтая Алтайского государственного университета] and Anthropology Department of Tomsk State University [Кабинета антропологии Томского госуниверситета]), excavators, and processor archaeologists and anthropologists for analysing the human remains published along the study, which included destructive ancient DNA analyses. These human remains were processed through minimally destructive methods, ensuring treatment with dignity and respect. The funders had no role in any stage of the study, starting from its design and sample collection until the publication of the study.

## Data Availability

All of the data needed for the evaluation of the conclusions are present in the paper and/or the Supplementary Materials. Newly published sequence data can be accessed in the European Nucleotide Archive (ENA) with the accession number xxx (which will be available upon acceptance). Samples without sufficient ancient DNA are reported in Supplementary Table S1.

## Materials and Methods

### Laboratory Work on Ancient DNA

Each step of the laboratory work was performed in the dedicated ancient DNA laboratory premises at the Institute of Archaeogenomics, HUN-REN (ELTE) Research Centre for the Humanities (Budapest, Hungary) under sterile conditions, in protective clothing. During the laboratory work, we applied irradiated UV-C light, DNA-ExitusPlus (AppliChem) and/or bleach for cleaning. We used blank controls for each step of the workflow. A total of 180 samples (petrous bones (n=150), teeth (n=28) and bone powders (n=2)) were taken for genetic investigation (Table S1). Cleaning of sample surfaces was done by mechanical cleaning and UV irradiation. Bone powder was drilled from the petrous bones (*66*) and teeth (*67*). DNA extraction was carried out with well-established protocols (*68*). Success of DNA extraction was verified with polymerase chain reaction (PCR) by the use of mtDNA primer pairs (*69*). Double stranded DNA libraries were prepared with half-UDG treatment as described previously (*70*), with minor modifications for automation. Each library was prepared with unique double internal barcode combinations. Libraries were processed for amplification with TwistAmp Basic (Twist DX Ltd) and purification with AMPure XP beads (Agilent), then were dual indexed with unique iP5 and iP7 indices for shotgun sequencing (*71*). The purification step was repeated with AMPure XP beads (Agilent). DNA concentrations were measured on Qubit 2.0 fluorometer, and fragment size was inspected with Agilent 4200 TapeStation System (Agilent High Sensitivity D1000 ScreenTape Assay). The samples were then screened with shallow shotgun sequencing using Illumina MiSeq System (2×75 bp) for endogenous DNA contents and duplication rates. Samples with high endogenous DNA content and low duplication rate (n=69) were directly sent to deep sequencing, while others (n=22) were enriched for ∼1.35 million SNPs using Twist Ancient DNA reagent kit (Twist Bioscience) before deep sequencing (2×150 bp) (*34*) (Table S1). Deep sequencing was performed using Illumina NovaSeq 6000, NovaSeq X Plus and HiSeq X (PE150) platforms by Novogene (UK) Company Limited (Cambridge, United Kingdom) and Macrogen Europe BV (Amsterdam, The Netherlands) companies. Details on dsDNA libraries and sequencing information are presented on Supplementary Table S1.

### Bioinformatic Sequence Data Processing

The *PAPline* (*72*) package (https://github.com/ArchGenIn/papline/tree/main/papline) was used with default settings in order to map the paired-end reads using the GRCH37.p13 reference sequence; pre-/post-filtering of the data; and calling mitochondrial sequences. The results are in Table S1. Genetic sex was determined using shallow shotgun sequences. Samples with <200 reads on the X chromosome were deemed uncertain for genetic sex. For the rest, we assigned male for the individuals with >10% fraction of Y chromosome/X chromosome reads, whereas a fraction of <3% was assigned female and fractions 3-10% were considered uncertain. Y-chromosomal haplogroups were determined using *Yleaf* (v2.2) (*73*) software. We used the public view Family Tree DNA Discover™ database to evaluate the Y-chromosomal haplogroups in ancient groups (*74*). mtDNA haplogroups were assigned with *HaploGrep* (v2) (*75*). Manual curation and automatic call results for Y-chromosomal and mtDNA haplogroups are presented in Supplementary Table S1.

Male X contamination ratio was measured with *ANGSD* (v0.939-10-g21ed01c, htslib 1.14-9-ge769401) (*76*) and *hapCon* (*hapROH* v0.60) (*77*). For the former method, the starting file was an output from *samtools* mpileup (version 1.10, htslib 1.10.2-3ubuntu0.1), with base quality 30 and mapping quality 25. We used the recommended default parameters in the official documentation. The used chrX_1240k.bed and chrX_1240k.hdf5 files were also supplied by the developers in the software package. The function doCounts of *ANGSD* was run on the X-chromosome between position 5500000 to 154000000, with mapping quality 25 and base quality 30. The toolkit’s contamination executable was then used to process the resulting file, with these parameters: -b 5500000 -c 154000000 -d 3 -e 100 -p 1. The required file for the -h parameter (HapMapChrX.gz) was the one acquired with the software. All individuals were tested for mtDNA contamination with *ContamMix* software (v1.0-10) (*78*) with previously discussed steps (*4*). Individuals with *ANGSD* contamination value of lower bound (of the %95 confidence interval) >5% (*79*) or mtDNA contamination rate >5% (*4*) were excluded from further genetic analyses. Results from contamination analyses are presented in Supplementary Table S1.

For genetic relatedness estimation we used *READ* (v2) (*45*) and *KIN* (v3.1.3) (*46*) softwares (Table S9). We only considered individual pairs as true positives for genetic relatedness if the pair was present in the results coming from both programs (on 1st-3rd degree level), and the log likelihood ratio from KIN was ≥ 1, in order to prevent false positives. The list of kinship analysis results can be found in Supplementary Table S9.

### Isotope Analyses and Radiocarbon Dating

We chose 14 samples to be dated by Isotoptech Zrt. in Debrecen, Hungary using accelerator mass spectrometry (AMS) (Table S1). Genetic outliers and individuals with obscure archaeological information or dating were chosen for absolute dating in this project. During this work, the collagen extraction and purification process was performed using a modified Longin method, extended with a plus ultrafiltration step using VIVASPIN 15R, 30,000 MWCO HY ultrafilters (*80*). The conventional 14C ages were measured by a MICADAS type accelerator mass spectrometer (AMS) and, after data evaluation, they were converted to calendar ages using the program OxCal 4.4.4 and the IntCal20 calibration curve (*81–84*). The collagen matter of these samples were additionally analysed for stable carbon and nitrogen isotope ratio (expressed in δ^13^C and δ^15^N notations) (Table S1). Five individuals in the newly analysed dataset, three from the Altai_FS_Odintsovo_1 analysis group, yielded δ ^15^N values higher than 12‰, in addition the δ^13^C values of these three samples were relatively more negative (<-20‰). These individuals also revealed earlier dates from the previous evaluations, which suggested a case of freshwater reservoir effect (FRE) (Table S1; Text S8). Freshwater reservoir effect has previously been shown to cause ^14^C dates to shift to older age ranges (*85–87*), therefore we followed the archaeological dating for those individuals. Results of isotope analyses are presented in Supplementary Table S1, and the calculations done regarding freshwater reservoir effect can be found in Supplementary Text S8.

### Population Genetic Analyses

The principal component analysis (PCA) was done with *smartpca* software (v16000) in *EIGENSOFT* (v8.0.0) (*39*, *40*), with “lsqproject: YES” and “shrinkmode: YES” parameters. For the PCA calculations, 2121 modern-day Eurasians present in the AADR database (v54.1) Affymetrix Human Origins (HO) array (*36*) were used to cover the Eurasian genetic variation better, and to avoid miscalculations resulting from missing data in ancient genomes (Table S2). Outgroup *-f*_3_ analyses were also performed using the modern groups in the HO database (Table S2). Supervised *ADMIXTURE*, *f*_4_/*D-*statistics and *qpAdm* analyses were performed using reference data from the AADR database (v54.1) 1240k SNP dataset (*36*) (Table S2). Individuals of the newly analysed dataset with less than 50,000 SNPs on the 1240k SNP panel were discarded from all analyses. For further information on the analyses, see below.

### Sample Grouping, Labeling and Outlier Detection

We incorporated previously published ancient Altaians with related archaeological contexts into our dataset, as explained in detail in Supplementary Text S2 and listed in Supplementary Table S2, which contains the full list of the published ancient and modern individuals used in this study. We initially labelled the newly analysed samples individually based on the geographical regions, as well as the material cultures and periods of the individuals, to which they were assigned through archaeological methods (Fig. 1; Table S1; Text S1). We detected the outliers of the chronological-geographical groups based on the chi-square distributions and Mahalanobis distances on the samples on the principal component of the PCA (PC1-PC2 and PC1-PC3 spaces) (*88*). We labeled the outliers based on the p-values of the samples in the tests (<0.05) in either of the PCA spaces, where the results are presented in Supplementary Table S3. For the non-outlier individuals, we further divided the groups with the same labels into subgroups of different genetic profiles found within these regional groups based on combined results of PCA and supervised *ADMIXTURE* analyses (e.g. Altai_FS_Srostki_1, Altai_FS_Srostki_2) (Fig. 1, 2, S2-S4). With this, we aimed to increase the resolution of the subsequent genetic analyses, and to explain the population events more accurately while adhering to the definition of genetically reassigned groups.

In order to run *ADMIXTURE* (v1.3.0) analysis (*41*) we first pruned our dataset with *plink* (v1.9) software (*89*). We used the “-geno” 0.999 flag to include only the sites that were covered at least once in most of the ADMIXTURE dataset. We then pruned for linkage disequilibrium (LD) with “-indep-pairwise” 200 25 0.4 parameters. 957,956 SNPs remained after pruning. We performed K=6 supervised ADMIXTURE with “-supervised” parameter. We used six Neolithic – Early Bronze Age groups as sources to model deep ancestries with previously displayed genetic impact on later groups; Turkey_N (n=33), Iran_GanjDareh_N (Iran_N, n=10), Russia_Samara_EBA_Yamnaya (Steppe_EBA, n=10), Eneolithic Baikalians (Baikal_EN, n=17), Late Neolithic Chinese (China_LN, n=11), and a combined group of the closely related West Siberian hunter-gatherers and Botai culture individuals (WSHG_Botai, n=4 (*7–9*, *11*)) (Fig. 2, S4; Table S2). The list of the ancient individuals used for these source groups can be found in Supplementary Table S2, and the supervised *ADMIXTURE* bar charts of the whole set is shown in Figure S4.

### Ancestry Modelling and Admixture Dating

For group-based outgroup-*f*_3_-statistics, *f*_4_*/D*-statistics and *qpAdm*, we retained only the individual with the highest number of 1240k SNP hits between genetically related ancient Altaian individuals (up to 3rd degree) (Table S9). Outgroup-*f*_3_-statistics and *f*_4_*/D*-statistics were computed with *ADMIXTOOLS* software (v7.0.2) (*42*) commands qp3pop (v651) and qpDstat (v980) respectively. During the outgroup-*f*_3_ analysis we followed the model *f*_3_(Test, Target; Outgroup) and used “inbreed: NO” and “printsd: YES” qp3pop parameters. We calculated *f*_4_*/D*-statistics in the form of *f*_4_*/D*(Outgroup, Target; Test1, Test2) and set “inbreed: NO”, “printsd: YES”, “f4mode: YES” (for *f*_4_-statistics) and “f4mode: NO” (for *D*-statistics) as qpDstat parameters. We used |Z-score| ≥ 3 as the significance threshold. Results from the outgroup-*f*_3_-statistics and *f*_4_*/D*-statistics are provided in Supplementary Tables S4 and S5.

The *qpAdm* analyses were performed in *ADMIXTOOLS* (v2) (*43*) R package, with parameter “allsnps: YES” (represented as allsnps=T in the package). For these analyses, we used only the samples with more than 100,000 SNPs on the 1240k SNP panel as targets (Table S1, S6). For the proximal ancestry modeling in *qpAdm*, we used a right-group set of populations with differentially related ancestries to the left-group set (source and target) populations (*90*), where the right-group set was: Mbuti.DG, Serbia_IronGates_Mesolithic, Turkey_N, Iran_GanjDareh_N, Russia_CentralYakutia_LN.SG, Russia_DevilsCave_N.SG, Russia_Samara_EBA_Yamnaya, WSHG_Botai, prepared to create high resolution models on an Inner Asian scale. The right-group set populations of the deep ancestry models on the other hand were as follows: Mbuti.DG, Israel_Natufian, Turkey_N, Iran_GanjDareh_N, Italy_North_Villabruna_HG, ONG.SG (Onge), Ami.DG, Mixe.DG. In the proximal models, we ran *qpAdm* analyses using previously and newly published groups from the Iron Age and later Central/Inner Asian groups as sources, whereas in the deep ancestral models we used pre-Iron Age groups as sources (Table S2, S6). It was recently shown that *qpAdm* is prone to a high rate of false positive models (*91*), so we sought regional models and sources that generalise biological changes and avoid hidden false positives. We retained only the lowest-rank plausible models (*90*) with component standard error ≤ 0.2 and no negative values. Significance was set at p = 0.05 (Table S6). The resulting feasible models are listed in Supplementary Table S6.

We inferred the time of admixture events by using *DATES* (v753) (*9*) software, reporting results in generations and converting them to years, assuming 29 years per generation (*2*, *3*, *42*). The same source groups as in the regional proximal qpAdm models were used to date the admixture of each analysis group. We presented the inferred dates as generations and approximate years relative to the mean time of the analysis groups in Supplementary Table S7.

### Runs of Homozygosity (ROH) and Effective Population Sizes (*Ne*)

For detecting runs of homozygosity (ROH) blocks and signals of consanguinity, and estimating effective population sizes (*N_e_*), we used the *hapROH* (*44*) software. ROH was calculated with default parameters and pseudo-haploid genotypes with a threshold of at least 400,000 SNP hits in the 1240k SNP dataset. *N_e_* calculations were done using the curvature of the likelihood method implemented in the *Ne* vignette of the software. hapROH results are displayed in Figure S7, and the effective population sizes of the analysis groups are provided in Supplementary Table S8.

### Imputation

We considered individuals with at least 300,000 SNP hits on the 1240k SNP dataset for imputation, which was performed using the *GLIMPSE* (v1.1.1) software suite (*92*). The process mainly followed the official GRCh37 tutorial of the developer, using the precompiled static binaries (v1.1.1). As reference, we used the 1000 Genomes Project (30X from NYGC) panel on hg19/GRcH37, provided by Ali Akbari (Harvard Medical School). We computed genotype likelihoods with a standard method that includes bcftools, mpileup, bcftools call (v1.10.2) (*93*), with both the base- and mapping quality (flags -Q and -q) set to 20. Differently from the official workflow, we determined chromosomal level imputation regions on a per sample basis, with the executable GLIMPSE_chunk_static, applying default parameters (--window-size 2000000 --buffer-size 200000).

### Identity-by-Descent Autosomal Haplotype Segment Sharing Analysis

The criteria for including the samples into identity-by-descent (IBD) analysis were the following: higher than 0.99 genotype probability (GP) for the 1240k SNP variants and fraction≥0.8 on Chromosome 3 after imputation. Successfully imputed samples were analysed together with a set of published samples imputed with the same process, on a dataset of total 2426 ancient Eurasian individuals. We used *ancIBD* (v0.5) (*47*) software for IBD calculations with the default parameters. We built the IBD networks with *Gephi* (v0.9.2) (*94*) software. We kept only the IA and later periods’ samples of the published datasets, with a focus on Central/Inner Asia and the steppe context. To avoid false-positive IBD hits and increase the accuracy of the results, we required at least one segment of 12 cM connections (edges) per individual (node) pair respectively (Table S11). After the filtering criteria, 846 samples remained on the largest connected network of the 12 cM dataset, which we used for downstream analyses (Table S10). Relevant information on these individuals and the IBD links are provided in Supplementary Tables S10 and S11.

For objective analysis of the IBD dataset prior to evaluating the modules based on the analysis groups, we investigated computationally-inferred clustering and the IBD metrics within the 12 cM IBD dataset. We segmented the distribution of edge weights in the 12 cM IBD dataset by dividing the IBD connections into 28 bins, each spanning 125 cM (Table S11). We built the graphs using the *MultiGravity ForceAtlas 2* (*95*) layout algorithm with the following settings: Approximation = 1.2, Approximate Repulsion = True, Gravity = 1.0, Dissuade Hubs = True, Prevent Overlap = True, and Edge Weight Influence = 1. IBD-sharing communities (clusters) of the network were defined using the *Leiden* clustering algorithm (*96*), providing an unbiased approach on the clustering of the nodes. We ran the algorithm in ten steps on the 12 cM dataset, increasing the “resolution” parameter from 0.01 to 0.1 in increments of 0.01. For interpretation, we selected the 0.01 resolution level, which produced the fewest clusters while including the largest number of samples (Table S11). We defined the subclusters of this dataset on the 0.03 resolution level. We calculated the metrics of this dataset on the 0.01 resolution level using the network overview functions in *Gephi*, after filtering out the IBD links higher than 1200 cM to prevent bias from close relatedness. Clique metric calculations were done based on the cliques detected with *CFinder* (v1.21) (*97*) software, on a dataset not filtered for this 1200 cM threshold, to avoid losing valuable data for the cliques.

After observing and explaining the informative newly-analysed dataset as above, we proceeded with modules of new and published analysis groups (Table S10). Since not every published individual can pass the imputation for the IBD analysis, the IBD dataset differed from the dataset for customary population genetic analyses. We aimed to pool contextually –regionally and chronologically– similar non-Altaian individuals together where the published information was limited, to maximize understandability and analysis quality, which resulted in differences between the two datasets (Table S2, S10). We then evaluated the modules with a threshold for at least one segment of 20 cM edges per node pair, by which we aimed to catch recent relatedness and strong links between the groups (*47*). For the main text, we compared the within- and between-module links of the modules with Fisher’s exact test (Table S12). Details of the IBD analyses can be found in Supplementary Text S6.

## Abbreviations

ANA: Ancient Northeast Asian
ANE: Ancient North Eurasian
BA: Bronze Age
BMAC: Bactria-Margiana Archaeological Complex
EBA: Early Bronze Age
EN: Eneolithic
FS: Forest-Steppe
HG: Hunter-gatherer
IA: Iron Age
IBD: Identity-by-descent
Mt: Mountainous Altai
N: Neolithic
*N_e_*: Effective population size
NEAHG: North Eurasian hunter-gatherer
PCA: Principal component analysis
ROH: Runs of homozygosity
WSHG: West Siberian hunter-gatherers

