## Supplementary Text and Supplementary Figures S1.1-S1.35 for "Ancient human genomes from the Altai region reveal population continuity and shifts in the 4th-12th centuries"

**The file includes:**

**Supplementary Text**

**S1:** Archaeological information of the cultures and samples
*Bulan-Koby culture*
*Odintsovo and Upper Ob cultures*
*Turkic culture*
*Srostki culture*
**S2:** Previously published Altaian groups involved in the study
**S3:** Outlier detection in the newly analysed dataset
**S4:** Outgroup-$f_3$-statistics and $f_4$-statistics
**S5:** Genetic kinship analyses
**S6:** Identity-by-descent analyses
*Leiden clustering and network metrics*
*Analysing the Altai through modules*
*Strong connections in the 20 cM threshold IBD dataset*
**S7:** Y-chromosomal haplogroup data
**S8:** Freshwater reservoir effect (FRE) analysis of the bone samples

**Figs.** S1.1-1.35

**Supplementary Tables**

**Supplementary Text S1: Archaeological information of the cultures and samples**

**a. *Bulan-Koby culture***

In the last two centuries of the 1st millennium BCE and the first half of the 1st millennium CE (around 200 BCE – 500 CE), a large association of tribes inhabited the Altai region and the adjacent territories, leaving behind the monuments of the Bulan-Koby culture (*19*) (Fig. S1.1). This archaeological culture has been well studied. Approximately 800 burial mounds have been excavated, the majority of which were undisturbed and unlooted. Additionally, settlements, hillforts and petroglyphs have all been discovered. Excavations of these sites have yielded a substantial body of material evidence, characterising the formation and development of the ancient nomadic society. The name 'Bulan-Koby' culture was coined in 1990 deriving from the name of a large burial complex Bulan-Koby-IV in the Central Altai, studied by Y.T. Mamadakov. However, the investigation of sites belonging to this cultural horizon began much earlier, with the excavations led by V.V. Radlov in 1865 at the Katanda-I and Berel sites.

For this article, sites of the late stage of the Bulan-Koby culture were used (Verh-Uymon –or Verkh-Uimon– stage). They are mainly dated to the second half of the 4th – first half of the 5th centuries CE and were contemporaneous with the period of the Rouran Khaganate's existence in Inner Asia, well attested in Chinese written sources. The Yaloman-II, Choburak-I and Ust`-Biyke-III cemeteries belong to the Verh-Uymon stage. The burial rites of the late Bulan-Koby stage do not show any fundamental changes compared to the previous two stages. The political organization of the Altai nomads in the second half of the 4th-5th centuries CE appears to have been a tribal confederation. Archaeological materials of the Verh-Uymon stage demonstrate the incompleteness of the process of ethnogenesis of the Bulan-Koby, which was expressed in the coexistence of different burial traditions representing groups of differing social status. The excavated assemblages demonstrate a militarized, horse-centric culture. Sophisticated forms of iron armour, weapons for ramming/crushing, and long-bladed and other weapons were widespread. The material culture largely continued the development of Late Xianbei traditions. In 460 CE, the Rourans relocated the Ashina tribe from East Turkestan to the Altai, which managed to consolidate the Bulan-Koby tribes around them. As a result, the community known historically by the endonym 'Türk' was formed. An important feature in the Bulan-Koby culture is the spread of a material complex (rigid saddles with wooden frames, tubular quivers, stirrups, horn buckles, etc.), which later, with some modifications, became characteristic of the Turkic culture of the second half of the 5th-11th centuries CE.

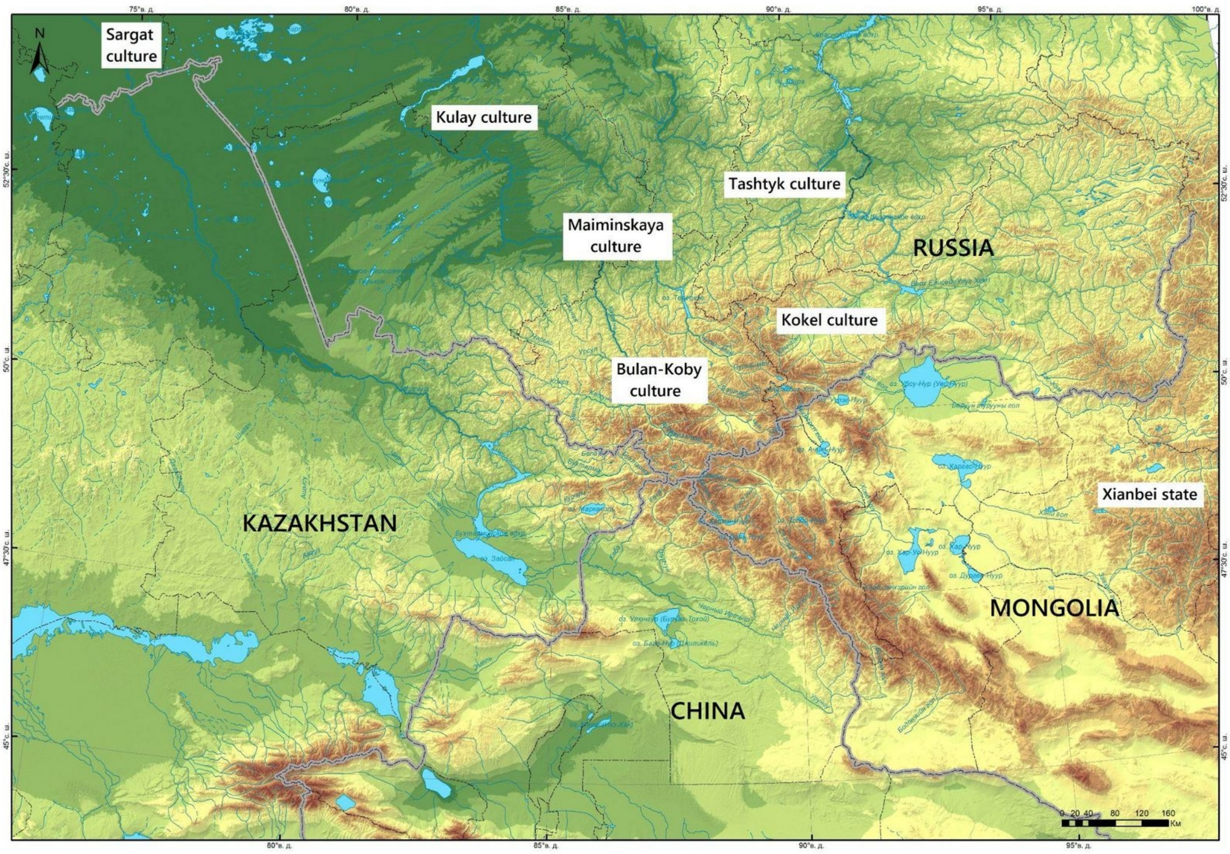


**Fig. S1.1.** The main region where the Bulan-Koby culture monuments are concentrated (adapted from (*19*), fig. 2.195).

**Choburak-I**

The Choburak-I burial and memorial complex is located 4.6 km south-southeast of the village of Elanda, Chemalsky District, Altai Republic (Russia), on the right bank of the Katun River, 1 km east of the confluence of the Choburak Creek (51.10424 N, 86.06785 E). The archaeological site included 12 kurgans that formed a local necropolis (Fig. S1.2). All burials were found to be undisturbed by looting, and had numerous material finds. Samples from Kurgans No. 29, 31, 31a, 32 and 38, which the materials are fully published (*98*), were used for this study (Fig. S1.2). Based on the analysis of numerous archaeological materials and AMS-dating of 26 samples, this necropolis of the Bulan-Koby culture was dated to the second half of the 4th century CE (*98*).

In **Kurgan No. 29**, the skeleton of a child aged 9-11 years was recorded, laid supine with its head oriented to the northwest. Iron and bone arrowheads, part of an iron knife and an iron buckle were found in association with the remains.

**Kurgan No. 31** contained an undisturbed skeleton of a male of approximately 40 years. The deceased, with a set of accompanying implements, was lying supine, stretched out on his back, with his head to the northwest. The bones bore incised wounds with no signs of healing. In the south-eastern half of the pit there was a burial of a riding horse, the skeleton of which overlapped to the middle of the human femur bones. There were items of horse equipment with it.

In **Kurgan No. 31a**, the skeleton of an elderly male, of over 55 years old, was found lying supine with his head to the northwest. The deceased was accompanied by a variety of burial implements consisting of weapons, equipment, and tools. In the south-eastern half of the pit there was an accompanying burial of a riding horse with the typical riding equipment. The animal overlapped the legs of the male.

In **Kurgan No. 32**, a male of 25-30 years old was buried. He was lying supine with his head to the northwest. Different categories of weapons, equipment, and tools were found in the grave. In the southeastern half of the pit, there was an accompanying burial of a horse with a set of horse equipment.

**Kurgan No. 38** contained the burial of a male of 25-30 years of age, placed supine and oriented with his head to the northwest. A variety of burial inventory was found with the male, which included items of armament, equipment elements, and tools. As in the previous kurgans, at the 'feet' of the deceased there was an accompanying burial of a riding horse with equipment.

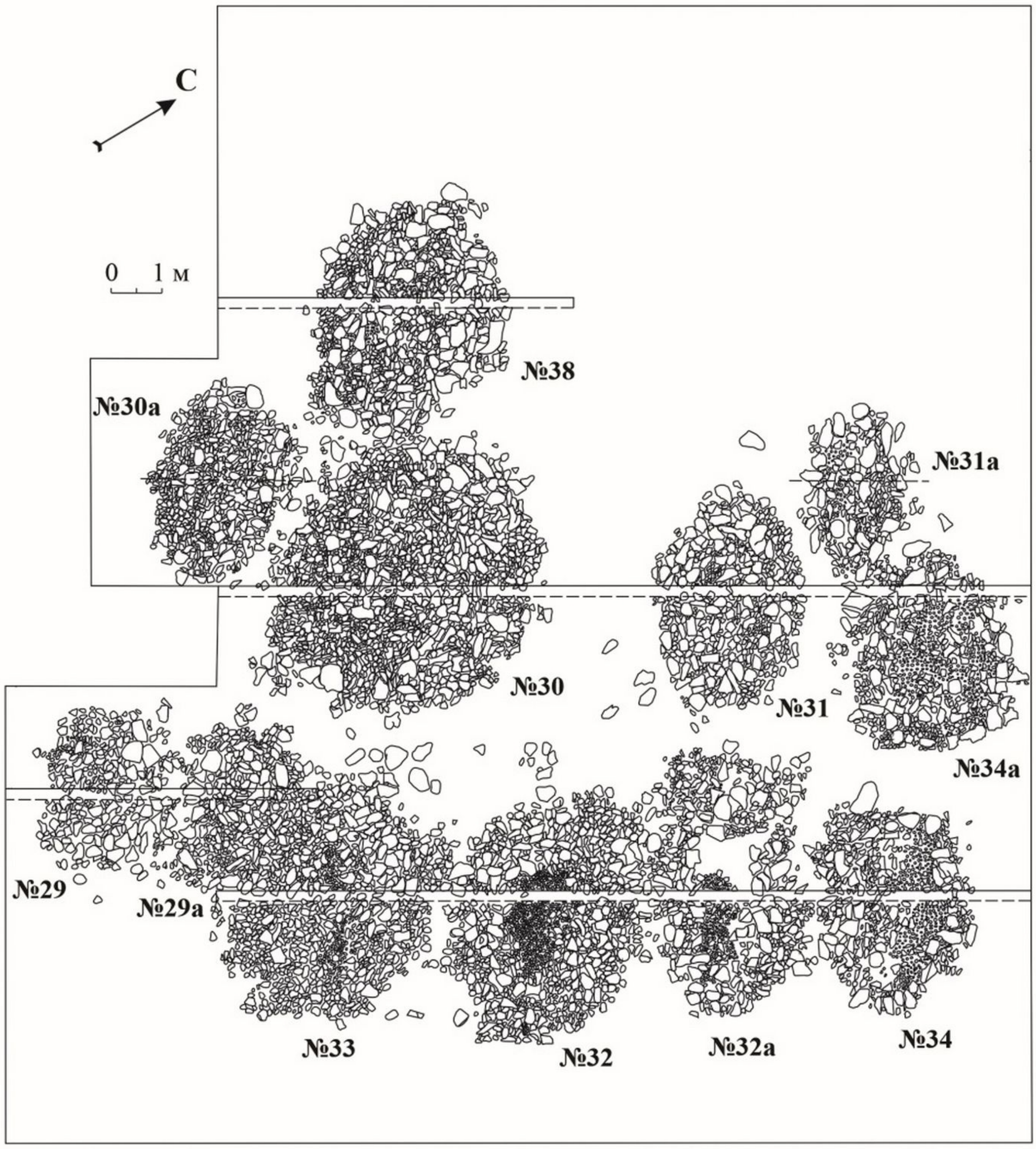


**Fig. S1.2.** Plan of the Choburak-I necropolis of the Bulan-Koby culture. The arrow labeled "C" (*Север*, "North") indicates the direction of the north.

**Yaloman-II**

The Yaloman-II archaeological complex (*99–101*) is located in the Ongudai district of the Altai Republic (Russia), on the left bank of the Bolshoy Yaloman River, 0.4 km northwest of its confluence with the Katun River (50.313144 N, 86.332254 E). In 2003, Altai State University excavated four kurgans of the late (Verh-Uymon) stage of the Bulan-Koby culture. They were stone constructions of rounded and oval shape, approximately 1 m high, with a ringwall of flat slabs around the perimeter. The kurgans, which were attached to each other, formed a cluster which outwardly resembled a bee honeycomb. The inner space of the mounds consisted of the remaining soil, which was left after backfilling the graves. The top of this soil was covered with stones. Beneath these structures were one or two relatively deep (1.5 to 2.7 m) pits of elongated-oval shape, serving as the graves. At the bottom of the graves were narrow stone boxes of rectangular or trapezoidal shape, decorated and covered with thin slabs from above. The deceased people lay stretched out on their backs in supine positions, and were mostly oriented with their heads towards the northeastern sector of the horizon. A whole horse carcass or only parts of a horse (head with hide, limbs, and tail) were placed on the slab and behind the wall of the box. Excavations uncovered numerous ancient artifacts, including those made from preserved organic materials. The main finds were items of offensive and defensive equipment (swords, armour, bows, arrows, combat knives).

**Kurgan No. 33, Grave-1** revealed a double inhumation of children laid in a 'valet' position ('валетом', with their heads oriented in opposite directions) (Fig. S1.3). The article uses the results of genetic analysis of a sample from the skeleton of one of them. The peculiarity of the burial is the presence of preserved wooden bases of a rigid saddle, which was used without stirrups. Taking into account the high degree of validity of the chronology of the late group of the Yaloman-II necropolis, this site can be considered one of the basic sites for identifying indicators of the pre-Turkic period. The analysis of archaeological materials and several radiocarbon dates obtained from samples from the mentioned kurgans allows us to determine the chronology of the studied group within the late 4th - first half of the 5th centuries CE.

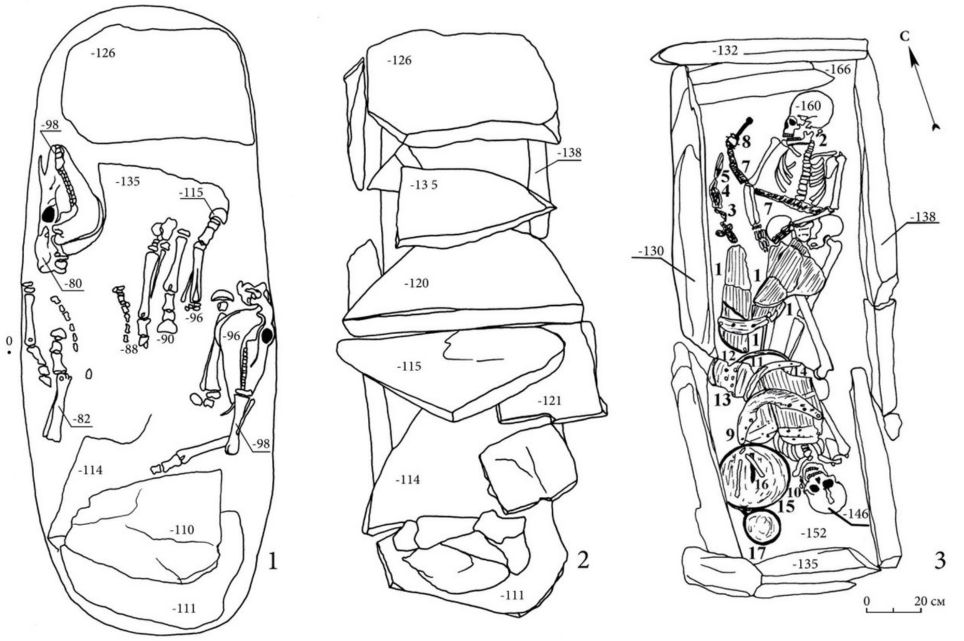


**Fig. S1.3.** Yaloman-II. Mound No. 33, grave-1: 1 - discovered parts from two horse skeletons; 2 - cover of a stone box; 3 - burial of children with saddles. The arrow labeled "C" (*Север*, "North") indicates the direction of the north.

**Ust`-Biyke-III**

The kurgan burial site Ust`-Biyke-III is located in the Chemalsky District, Altai Republic (Altai), approximately 0.55 m northwest of the mouth of the large Biyke brook, a right tributary of the Katun (51.10243 N, 86.9387 E). In 1997, the archaeological team of Altai State University investigated **Kurgan No. 4** (*102*), which had a small stone mound (Fig. S1.4). The grave pit revealed the burial of a supine male with his head to the southwest. The accompanying inventory was a typical representation of the grave goods in the final (Verh-Uymon) stage of the Bulan-Koby culture. Based on the analysis of armoury items, Kurgan No. 4 was dated to the second half of the 4th - first half of the 5th centuries CE.

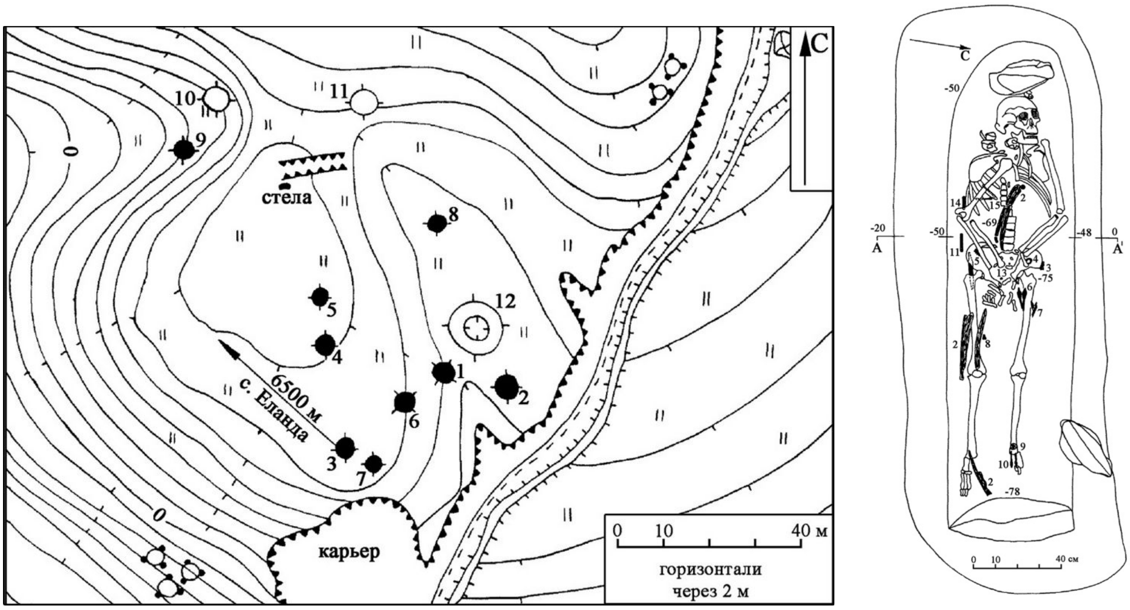


**Fig. S1.4.** Plan of the burial site Ust`-Biyke-III on the left and the burial in Kurgan No. 4 on the right (1, 4 - iron buckles; 2 - bone bow plates; 3 - horn composite buckle; 5, 13 - fragments of ironware; 6, 7, 10 - iron and bone arrowheads; 8, 9 - bronze plates; 11 - stone pick and bone tube; 12 - iron knife). The arrow labeled “C” (*Север*, “North”) indicates the direction of the north.

**Tytkesten’-VI**

The Tytkesken’-VI site (*103*), situated on the left bank of the Katun River, approximately two kilometers south–southwest of the village of Yelanda in the Altai Republic, represents a key multi-period archaeological complex of the region (*103*). Excavations by the Altai State University revealed both a settlement and a burial-commemorative area composed of stone kurgans and ritual structures.

In 1989, six burial mounds and one ritual stone construction were excavated, followed by isolated finds from surface structures in 1990 (*104*). The burials were single inhumations, most commonly of males aged 25–60 years (adultus-senilis), placed on their backs (supine) with their heads oriented to the west. The burials were covered by flattened stone mounds with ring-shaped stone settings (*crepida*), and the pits often contained traces of wooden frames or coffin-like boxes (*105*). Accompanying grave goods included iron belt buckles, fittings, bone and iron arrowheads, composite bow fragments, knives, beads, and spindle whorls—all typical of the Bulan-Koby culture (*103*). In some cases, fire traces were discovered near the kurgans, possibly connected with ritual purification by fire, a practice also observed in contemporaneous Kokel culture burials in Tuva (*106*, *107*).

Comparative analysis of the material culture shows close parallels with the Hun–Xianbei–Rouran horizon across Northern and Central Asia. Radiocarbon dating of charcoal from a hearth within the settlement (1970 ± 105 BP) and typological correlations place the complex in the late 2nd to early 3rd century CE, aligning it with the period of Xianbei political dominance in Central Asia (*108*). Thus, Tytkesken'-VI serves as an important reference point for understanding the spread and transformation of steppe nomadic cultures in the Altai during the early stages of the Migration Period.

**Kurgan No. 5:** The boundaries of the above-grave structure of this kurgan were difficult to determine, due to the proximity to three additional mounds. After the removal of the stonework, which consisted of rounded stones, a contour of a sub-rectangular grave pit with rounded corners was revealed (length: 1.47 m, width: 0.55 m, oriented west–east). At the bottom of the grave, 0.71 m below the ancient surface level, lay the burial of a child (presumably a female aged 6–7 years), placed in an extended supine position, with her head oriented to the west. The accompanying inventory included an unornamented clay spindle whorl, located near the left side of the head, and two beads lying in the area of the right shoulder and neck. The preservation of the skeletal remains was poor. The same condition was noted during the examination of other burials in this complex.

### b. *Odintsovo and Upper Ob cultures*

In the northern flat lands and foothills of the Forest-Steppe Altai a change of population took place in the 1st century BCE. Tribes of hunters and fishermen migrated from the West Siberian taiga (*20*). This process was caused by the humidification of the climate, which began at the turn of the 4th-3rd centuries BCE, leading to the swamping of forests in the taiga zone of Western Siberia and a reduction in the space suitable for habitation. The resulting excess population was forced to move south following the forest advancing on the steppe, beginning from the Narym Ob region. During the period of the 3rd-2nd centuries BCE, taiga tribes populated the territory of the Tomsk and Novosibirsk Ob regions, and from the turn of the 2nd-1st centuries BCE, their mass penetration into the forest-steppe regions of Altai began. The previous locals who lived there were partially exterminated or assimilated into the new population.

In the mid-4th century CE, various groups of nomads began to penetrate into the forest-steppe zones of the Altai, by then already inhabited by the Samoyedic tribes. In particular, from the Yueban principality–formed by the northern Xiongnu in Semirechye and engaged in conflict with the Rouran Khaganate in the 4th-5th centuries CE–a portion of the Kenkol culture's population living under this principality migrated to the Alei River basin, a left tributary of the Ob'. Additionally, from the Altai Mountains, the bearers of the Bulan-Koby culture –also involved in the military and political events of the period– moved northwards. As a result of contacts between these nomadic groups and the local Samoyed population, a new community was formed in the southernmost part of Western Siberia, represented by the sites attributed to the Odintsovo archaeological culture (*21*, *24*) (Fig. S1.5). Despite the multi-ethnic composition of this formation, the Samoyedic population remained dominant, and assimilated the migrants. In 1930. M.D. Kopytov and S.M.Sergeev excavated a ground grave near the settlement of Odintsovka. It was this site that gave the name to the culture, which was formally identified by T.N. Troitskaya in 1981. Along the northern border of the marked area of distribution of the Odintsovo culture sites (Fig. S1.5) there was a constant interaction with the bearers of the Upper Ob culture (Верхнеобская культура), the materials of which are also used in this article.

At the present, the notion of 'Odintsovo culture' is reserved only for the sites in the forest-steppe Altai. They date from the second half of the 4th to the first half of the 8th century CE and consist of settlements, hillforts and flat-ground burial sites, which are known on the right and left banks of the upper Ob River. Odintsovo settlements are located in forested areas on the banks of rivers and lakes, at the edges of steep and high terraces. A considerable portion of them are surrounded by a rampart and a ditch, though in some cases two concentric lines of such fortifications are observed. The main occupation of the Odintsovo culture bearers was semi-nomadic pastoralism. The deceased were buried according to the rites of inhumation and cremation. With the arrival of the new populations, burials of people with horses and with dogs began to appear. The cemeteries were established either on river floodplain elevations (елбаны) or on long promontories. There are several hidden burials made on high riverbanks and at great depth, belonging to the most noble and wealthy representatives of society.

The Turkic Khaganates had a great cultural influence on the local population in the 6th – 7th centuries CE. Sets of items such as armament, horse equipment, and jewellery made in the Turkic tradition appeared. However, these "Turkic" grave goods were found in burials using local funerary rites. Apart from Turkic items, these graves also contained inventory made in the Samoyedic tradition: arrowheads, knives and their scabbards, belt sets, pottery, and costume jewellery. It is noteworthy that many Turkic objects were not used for their original purpose. For example, various horse harness ornaments were found as part of personal belt ensembles in human burials. These observations testify only to the influence of the Turkic material culture on the population of the West Siberian forest-steppe, but not the arrival of Turkic groups at this period.

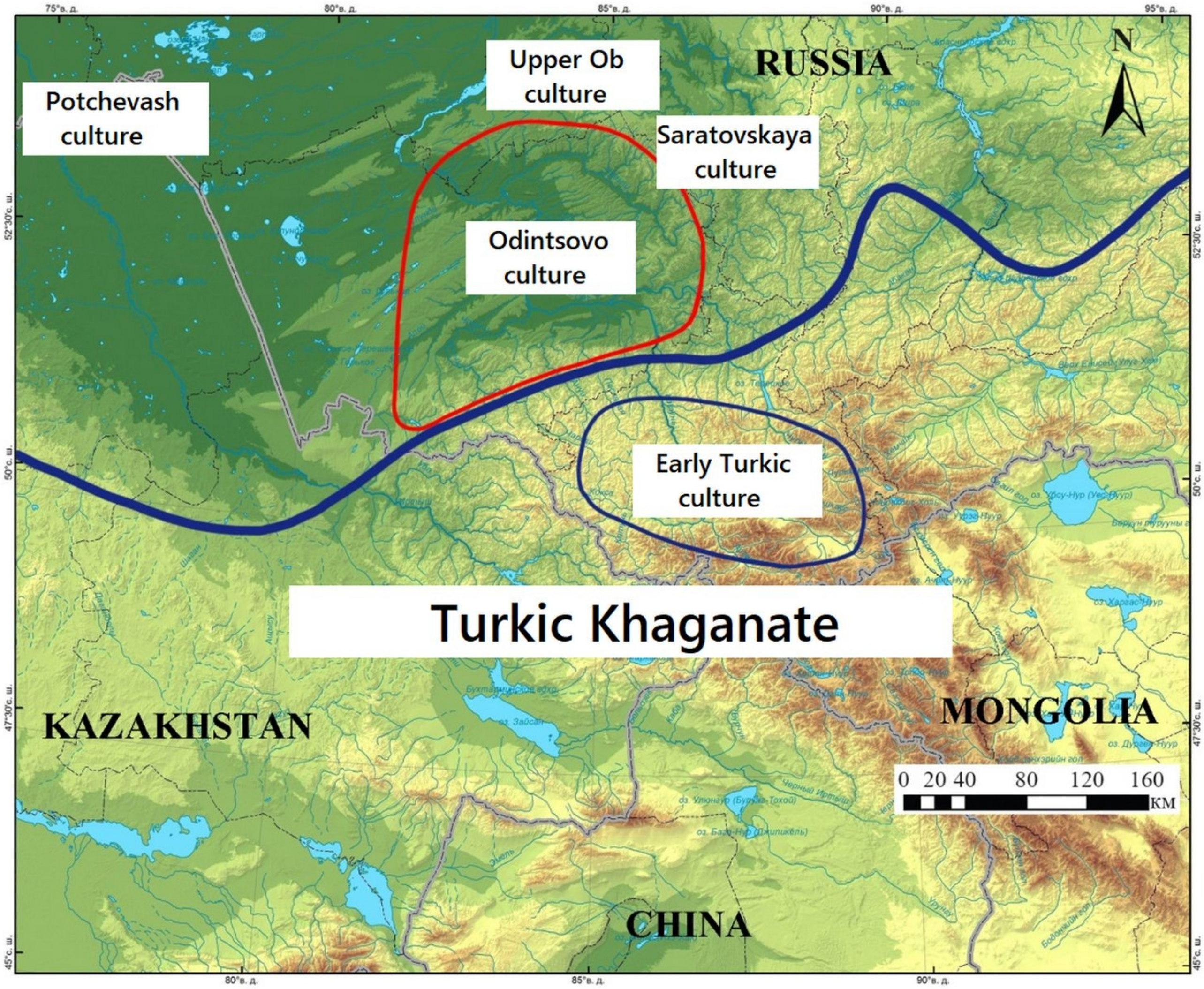


**Fig. S1.5.** The area of distribution of archaeological sites of the Odintsovo culture in the 6th century CE (adapted from (*21*), Fig. 3.11). The wide dark-blue line indicates the approximate area of the Turkic Khaganate in the region.

**Gorny-10**

The Gorny-10 flat-ground burial site is located on a promontory on the right bank of the Isha River, approximately 1.3 km west-northwest of the mouth of the Karaguzh River, and 0.6 km northwest of the Gorny settlement of Krasnogorsky District, Altai Krai (Russia) (52.166850 N, 86.081550 E). From 2000-2003, 75 burials were excavated in the necropolis area. The excavations were carried out by M.T. Abdulganeev and N.F. Stepanova. The materials of this site, currently one of the key complexes in the south of Western Siberia from the early Early Medieval period, have been only partially published (*109–115*). The dating of the necropolis is from the second half of the 6th - first half of the 8th centuries CE. At the same time, most of the material culture, judging by the available data, appears attributable to the late 6th - 7th centuries CE.

**Grave No. 7:** A single inhumation of a male aged 35-40 in a flat-ground grave. The deceased was laid extended and supine at the bottom of a shallow grave and oriented with his head to the northwest. A fairly representative accompanying inventory of grave goods was found, including items of armament, horse equipment, tools, and costume ornaments.

**Grave No. 8:** Burial of a male of 23-25 years in a flat-ground grave, accompanied by a horse and a dog. The deceased was laid at the bottom of a shallow grave in supine position and oriented with his head to the northwest. A very representative accompanying inventory was found, including weapons, horse equipment, tools, and costume ornaments.

**Grave No. 16:** A single inhumation of a male aged 20-30 in a flat-ground grave. The deceased was laid at the bottom of a shallow grave laid supine, and oriented with his head to the northwest. The small inventory included an iron buckle, a knife, and an arrowhead.

**Grave No. 20:** A single inhumation of a female of 20-25 years in a flat-ground grave. The deceased was laid at the bottom of a shallow grave laid supine and oriented with her head to the northwest. The small inventory included a small series of jewellery, an iron needle, and a buckle.

**Grave No. 24:** A single inhumation of a male aged 25-30 years in a flat-ground grave. The deceased was laid at the bottom of a shallow grave laid supine and oriented with his head to the northwest. A fairly representative accompanying inventory was found, including weapons, horse equipment, tools, and costume ornaments.

**Grave No. 27:** A single burial of a male aged 25-30 years in a flat-ground grave. The deceased was laid at the bottom of a shallow grave in supine position and oriented with his head to the north-northwest. A fairly representative accompanying inventory was found, including items of armament, horse equipment, tools and costume ornaments.

**Grave No. 29:** Partially destroyed burial (the central part of the site was damaged). A single burial of a mature male aged 50-55 years, laid at the bottom of a shallow grave and oriented with his head to the northwest. Some items related to an armoury set have been preserved.

**Grave No. 33:** A single burial of an elderly male over 55 years in age, in a flat-ground grave. The deceased was laid at the bottom of a shallow grave stretched out on his back and oriented with his head to the northwest. A fairly representative accompanying inventory was found, including weapons, tools, and costume ornaments.

**Grave No. 39:** Partially destroyed burial of a child of approximately 5-6 years. The deceased was laid at the bottom of a shallow grave placed stretched out, supine, and oriented with the head to the northwest. The few surviving implements included an iron knife, a paste bead, and a bronze ornament.

**Grave No. 65:** An extremely poorly preserved adult burial. Judging by the recorded situation, the deceased was laid at the bottom of a shallow grave extended and supine, and oriented with their head to the northwest. Some bronze objects and beads were preserved.

**Grave No. 66:** A single burial of a female aged 25-35 years in a flat-ground grave. The deceased was laid supine at the bottom of a shallow grave extended and supine, and oriented with her head to the northwest. A fairly representative inventory included a set of jewellery.

**Grave No. 67:** A single burial of a child of 11-12 years, in a flat-ground grave. The deceased was laid at the bottom of a shallow grave, placed supine and oriented with the head to the northwest. The small inventory included individual items of armament, tools, and elements of costume.

**Grave No. 72:** A single burial of a mature female of 50 or more years old in a flat-ground grave. The deceased was laid supine at the bottom of a shallow grave, stretched out on her back and oriented with her head to the northwest. The accompanying inventory included costume ornaments, horse equipment, and tools.

**Chumysh-Perekat**

The Chumysh-Perekat flat-ground burial site (Fig. S1.6) is located in Zalesovsky District of Altai Krai (Russia) (53.492843 N, 84.364370 E). It is located in the foothills of the Salair Ridge, on the right bank of the Chumysh River, on a high flat cape-like terrace. The dating of the necropolis is from the 6th - first half of the 8th centuries CE.

**Grave No.15:** The skeleton was discovered directly below the sod top layer. The inhumed individual (male?) was laid supine, with the head to the southwest. His legs were slightly bent at the knees, and arms placed along the body. The preservation of the skeleton was good, with a stature of 185 cm. The head was tilted to the right side. Six bone arrows were found to the right of the cranium, and a ceramic vessel turned upside down was found to the left. Under the mandible was a plate of non-ferrous metal. In the area of the pelvis, on the right, were fragments of a birch bark quiver, the shape of which was difficult to establish. A horn buckle was found under it. Bronze plaques and the remains of a leather strap were found near the left subcostal region. A fragment of pottery was on the thoracic part of the skeleton, and roe deer antlers were located above the skull of the deceased.

**Grave No. 21:** The grave pit was rectangular, with rounded corners (width 115 cm, length 200 cm, depth 60 cm from the ground level), oriented on the long axis along the south-north line with a slight offset. The buried male was laid supine, head to the north, at the western wall of the pit on a birch bark mat, which also covered the body from above. An iron buckle and an iron arrowhead, or dart, with remnants of the shaft, were found near the pelvic bones. At the left clavicle there was a cluster of objects consisting of corroded iron, a non-ferrous earring and remains of organics. On the left humerus there were 2 bone objects (one with notches), possibly bone plates. Under the mandible in the area of the cervical vertebrae, beads consisting of 18 objects (bronze, stone) and remnants of leather straps were found. A coloured metal earring was found to the right of the skull. In the eastern part of the pit, a bone product –possibly a bow plate– was found.

**Grave No. 26:** The grave pit was rectangular in shape, with rounded corners (dimensions: length 165 cm, width 110 cm, depth 53 cm from the ground level), oriented on the long axis along the line southeast-northwest. An earring was found in the fill. The inhumed individual, a child, was laid in the southeastern part of the pit close to the wall. The bones of the skeleton were partially displaced. The burial position was supine, with the head to the northwest. The skull was slightly turned to the right. The bones of the left arm were missing. The bones of the legs were displaced, and the feet were absent. The phalanges of the fingers were also missing or displaced. In the area of the cervical vertebrae, cylindrical non-ferrous metal pendants (8 short and 1 long) were found. Three additional pendants of a similar style were located in the northwest corner of the pit in front of the skull, and two more were found in the area of the pelvis. A poorly preserved iron object, possibly knife or dagger, was found under the pelvis and the left femur. Remains of organic decay were found under the pelvis. On the right side in the thoracic region, an iron object of poor preservation was found.

**Grave No. 29:** The grave pit was rectangular, with rounded corners. Length 130 cm, width 50 cm, depth 20-23 cm, oriented along the southeast-northwest axis. It contained a double inhumation.

*Skeleton 1*: The buried individual was laid in the southeastern half of the pit in such a way that the pelvic bones and leg bones were located on the side of the pit at the subsoil level. The skull, superior limb, and thoracic bones were located at the bottom of the pit. The buried person lay slumped on his right side, in right lateral decubitus. The left arm was bent at the elbow, with the hand placed on the elbow of the right arm. The right arm was bent at the elbow. The forearm bones (radius and ulna) were located under the left femur. The bones of the feet were partially preserved. A round-bottomed vessel, with a pronounced neck and a beaded (pearl-row) ornamented rim, was placed in front of the facial part of the skull, on the side, with the mouth downwards. Fragments of organics were found under the mandible. A fragment of an iron knife was found near the right hand. The northwestern half of the pit was covered with a partially burnt birch bark mat, on which 8 non-ferrous metal belt plaques, a buckle, and a bead were found. The second fragmented vessel was in the northeastern part of the pit, to the right of the mat.

*Skeleton 2*: After removing the birch bark behind the head of skeleton 1, another grave was discovered. The grave pit was an irregular-subrectangular shape with rounded corners. Dimensions: length - 128 cm, width - 50 cm, depth - 20 cm. At the bottom was a partially preserved skeleton of a child. Judging by the position of the remaining in situ remains, the buried individual was laid in an extended supine position on his back, with the head to the northwest. The remains of a wooden plank were found to the left of the cranium. The grave goods associated with this context included an iron knife and a plaque of non-ferrous metal.

**Grave No. 31:** The badly preserved burial was identified by the skull, which was recorded at a depth of 30 cm from the modern day surface. The bones of the skeleton formed 3 clusters, stretched along the line south-north, at a distance of 145 cm. The northern cluster consisted of bones of the cranium, clavicles and scapula. In the central part, mainly bones of the thoracic section were found. A ceramic vessel was also located here. The southern cluster contained a few bones and items: a copper based alloy ring and finger ring, a pendant, and a bead.

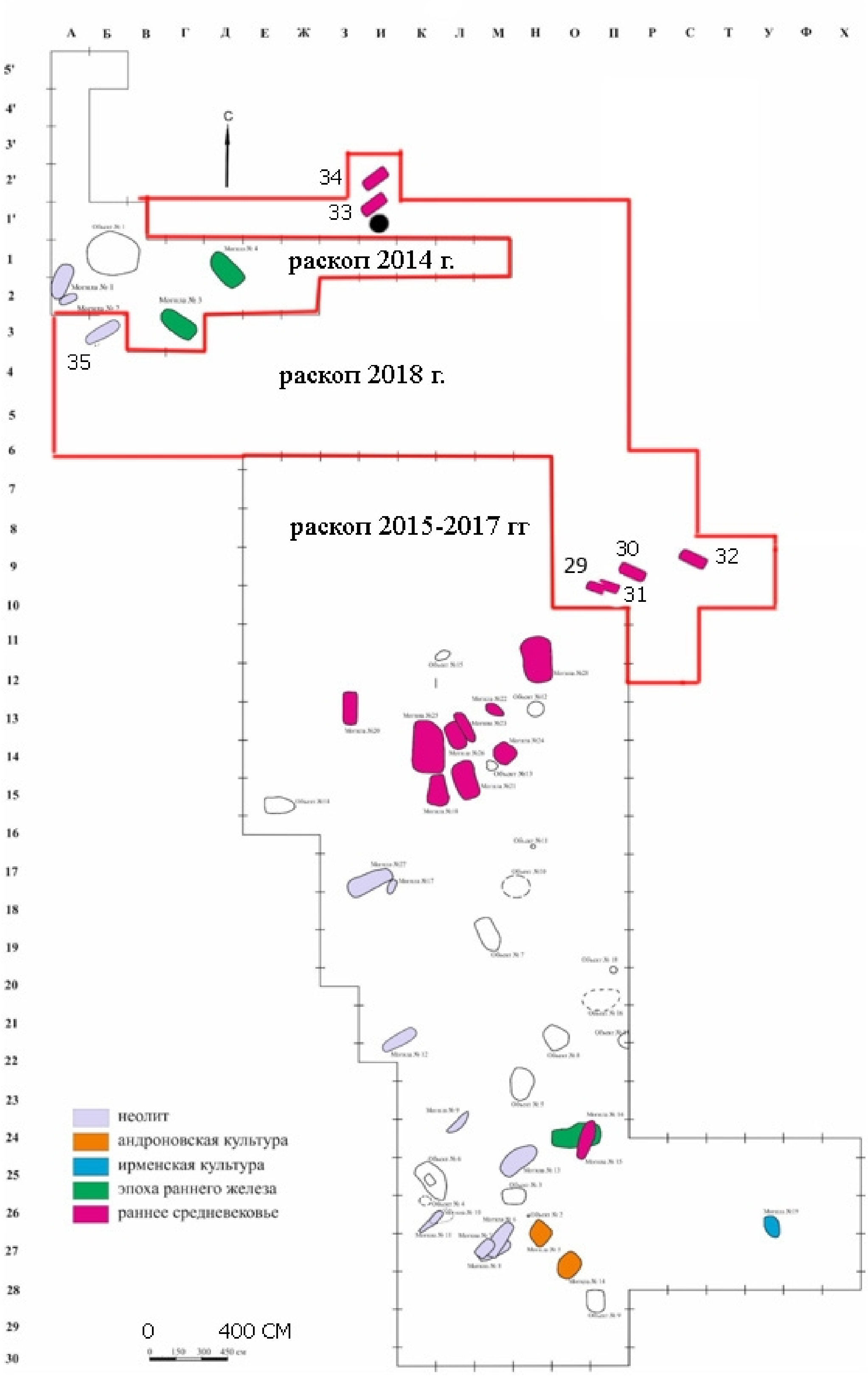


**Fig. S1.6:** Excavation plan of the Chumysh-Perekat burial site (in Russian language). The colours indicate the archaeological periods of the burials, where pink colour corresponds to the Medieval era, corresponding with the Odintsovo culture in this burial site. The arrow labeled "C" (*Север*, "North") indicates the direction of the north. The colours in the legend from top to bottom indicate the burials from the Neolithic period, Andronovo culture, Irmen culture, Early Iron Age and Early Medieval.

**Strashnyi Yar-1**

The archaeological site of Strashnyi Yar-1 is located in the modern territory of the city of Barnaul (Russia), approximately 3.5 km west of the settlement of Nauchny Gorodok, on a promontory of the left-bank terrace of the Ob River, 300 m from its edge along the eastern side of the eponymous ravine (53.25536 N, 83.274213 E). In 1999, an excavation of the site revealed two graves. One of them (**Grave-1**), belonging to the Odintsovo culture (Fig. S1.7), overlapped an earlier one (Grave-2), which dates to the Early Iron Age. A birch bark 'envelope' was first found in the fill of the grave pit, with traces of fire on its surface. Inside the envelope was the skeleton of an adult individual of undeterminable sex, which had been laid supine and fully extended with the head to the southwest. The accompanying inventory included the following set of finds: bronze pendants, remains of wooden items (a narrow box with a lid and a small vessel with a ring-shaped handle), an iron buckle, ten belt plaques, a birch bark quiver with three bone arrowheads and a bone buckle (Fig. S1.7). The grave also contained the skeletal remains of a dog and the skull of a small carnivorous animal. The inventory from this burial is dated to the 2nd half of the 6th - 1st half of the 7th century. Elements of the burial ritual find close analogies in later burials of the Odintsovo culture. The excavation materials have been published (*116–118*).

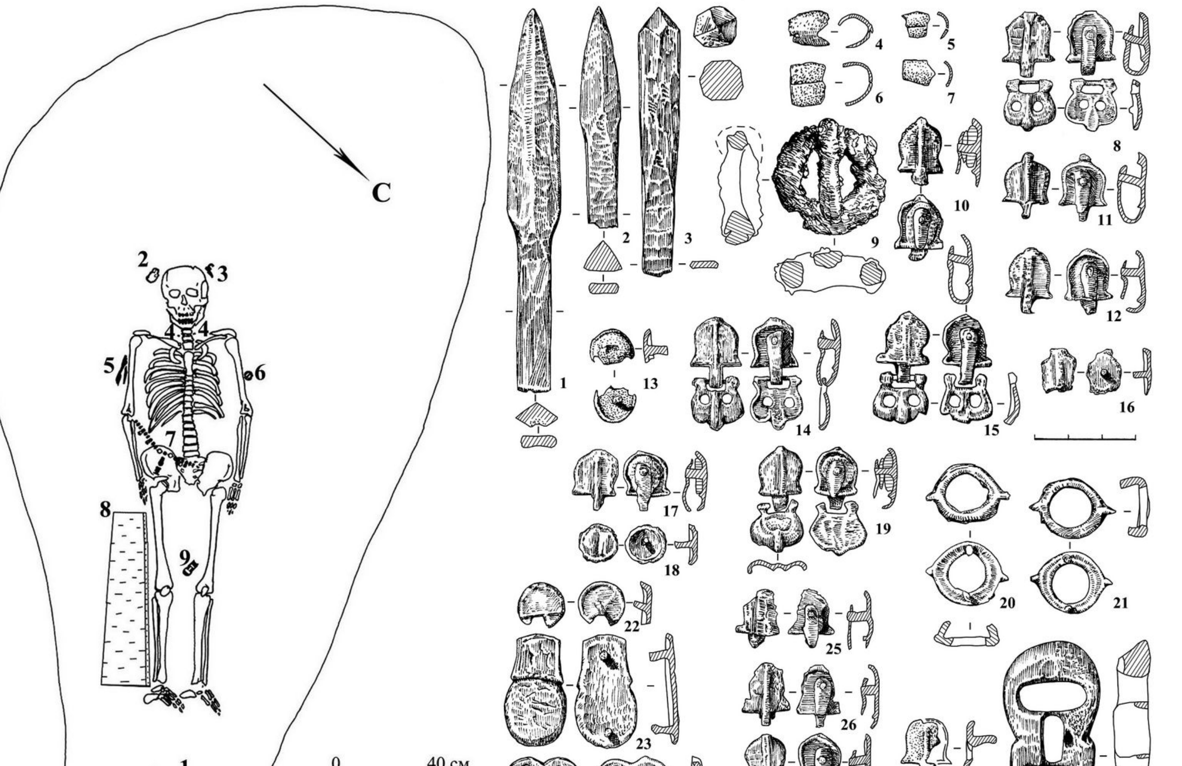


**Fig. S1.7.** Strashnyi Yar-1. Grave plan on the left: 1 - skeleton of a dog, 2 - skull of a small predator; 3 - wooden vessel; 4 - bronze pendants; 5 - wooden box; 6 - iron buckle; 7 - bronze plaques; 8 - birch bark quiver; 9 - bone buckle. The arrow labeled "C" (*Север*, "North") indicates the direction of the north. Finds on the right: 1-3 - arrowheads; 4-7 - pendants; 8, 10-29 - belt mounds; 9, 30 - buckles; 1-3, 30 - bone; 4-8, 10-29 - bronze; 9 - iron.

**Chekanovsky Log-IX**

The Chekanovsky Log-IX flat-ground burial site (*119*) is located in the Tretyakovsky District of Altai Krai (Russia), on the right bank of the Gilevskoye Reservoir, approximately 1 km northeast of the village of Korbolikha (51.52908 N, 81.59262 E). In 1998, the staff of the archaeological expedition of Barnaul State Pedagogical University, while examining the shore, discovered a part of a destroyed burial: the remains of two human skeletons laid in a supine extended position, with their heads to the north-northeast (Fig. S1.8). *Skeleton-1* belonged to a male aged 20-23, and *Skeleton-2* to a young male aged 17-19. According to the osteological assessment, cranial morphology and robust postcranial features indicate affinities with groups from the catacomb-sidewall-niche-grave traditions of the Tian Shan and Pamir-Alai regions. In the grave there were bone plates for two bows, knives, iron and bone arrowheads, sword fragments, buckles, a bone tube, a quiver loop and a part of iron bridles. Analysis of the inventory revealed the presence of two main components (Bulan-Koby and Kenkol), the presence of which is typical for burials of the early stage of the Odintsovo culture. The investigated burial at the Chekanovsky Log-IX site was dated to the second half of the 4th - 5th century CE.

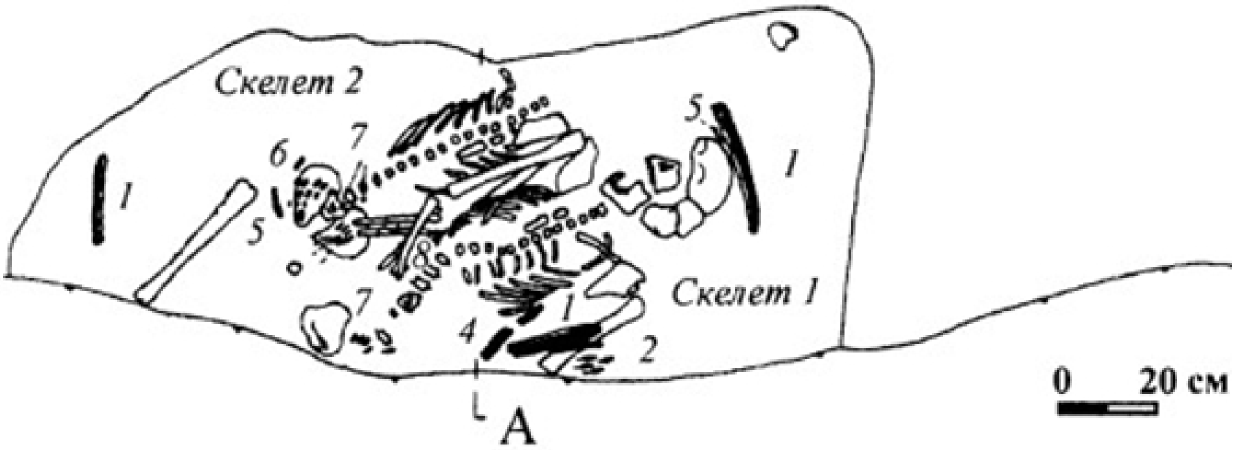


**Fig. S1.8.** The poorly-preserved burial of Chekanovsky Log-IX. "*Скелет*" stands for skeleton.

**Stepnoi Chumysh-1**

The Stepnoi Chumysh-1 flat-ground cemetery (*120*, *121*) is located in the Tselinnoye district of Altai Krai (Russia), on the left bank of the Chumysh River, and approximately 2 km below the village of Stepnoi Chumysh (53.9212 N, 85.574444 E). In 1978, the staff of a joint archaeological expedition of the Institute of Archaeology of the USSR Academy of Sciences and Barnaul State Pedagogical Institute excavated three burials of the 4th-5th centuries CE, similar in burial rites and material culture to the previously studied graves of the period. The obtained materials have not been published. In the Anthropology Cabinet of Tomsk State University under the number 2935 there is a damaged complete skull of a female individual of 40-45 years, with an artificial cranial deformation, which comes from **Grave 2**, investigated in the 5th excavation season of the necropolis of Stepnoi Chumysh-1. The available facts allow us to attribute the burial to the Odintsovo culture.

**Yurt-Akbalyk-8**

The Yurk-Akbalyk-8 kurgan burial site, located on the left bank of the Uen River flowing in the Ob floodplain, near the village of Yurt-Akbalyk, Kovan District, close to the border with Tomsk Oblast (Russia) (55.649376 N, 83.569158 E). It consisted of 52 kurgans with traces of looting and overgrown with cedar forest. In 1963, 1964 and 1975, T.N. Troitskaya excavated 36 kurgans, of which eight were completely empty. The material was partially published (*122*). This burial site is associated with the Medieval Upper Ob culture.

**Kurgan No. 27:** Measuring 9.2 m in diameter and 80 cm high, the kurgan contained coals, fragments of vessels, and burn spots. In the grave (2.15×2×0.12 m) five disturbed (reburied) skeletons were lying with their skulls to the northeast. The inhumed individuals consisted of three children and two adults. One of the buried adults had legs bent at the knees. A vessel, a bone arrowhead, and an iron article were found with the skeletons.

**Kurgan No. 28:** The kurgan measures 9.2 m in diameter, and 80 cm high. Within the mound, there were the skeletal remains of a dog, as well as coals and a fragment of a vessel. In a grave measuring 2.15×2×0.12 m, five reburied skeletons were found, three of which belonged to children and two to adults. The inhumed individuals were lying with their heads to the northeast, on their backs, and one of them had his legs bent. The grave had been looted previously. Several damaged bone arrowheads were found.

#### c. *Turkic culture*

Most researchers still refer to this culture as “ancient/old Turkic”. However, it is more correct to call it “Turkic”, without the addition of “ancient”, as the people who created it existed as a single community only in the early Medieval period. To date, the earliest archaeological sites of the Turkic culture dating to the second half of the 5th – first half of the 6th century CE have been identified and investigated only in the Altai, which reflects the processes of formation of the Old Turkic world at the early (Kyzyl-Tash) stage. It is important to note that such monuments demonstrate features of continuity with the local Bulan-Koby population’s traditions of the previous centuries, both in material culture and in funerary and memorial rites. The rich natural resources of the region were an important factor determining the rather rapid development of the Turkic culture in Altai. The most important were convenient pastures for grazing domestic animals, as well as abundant deposits of iron ore necessary for the manufacture of stirrups, weapons, and armour. As a result, Altai became the base for the formation of the Turkic community, whose entry into the political arena contributed to the creation of large nomadic empires of various scales.

To date, kurgan burial mounds, memorial structures, petroglyphs and runic inscriptions of the Turks have been studied practically in all parts of the region (*26*, *123*) (Fig. S1.9). The availability of significant archaeological materials has allowed us to outline the following stages of the development of the Turkic culture of Altai, and correlate them with historical events. The Kudyrge stage reflects not only clearly visible changes in material culture, but also the events of the military and political history of Inner Asia associated with the formation and existence of the First Turkic Khaganate (552–603 CE). The next stage, the Katanda (second half of the 7th - first half of the 8th centuries CE), reflects the revival of the Turkic culture and its dominance in Inner Asia, which is associated with the period of the Second Eastern Turkic Khaganate (682–744 CE). The Tuekta stage (second half of the 8th - first half of the 9th centuries CE) is chronologically connected with the time of the Uighur Khaganate. The next period reflects the relative dependence on the Kyrgyz Khaganate and is called the Kurai stage. It is dated from the second half of the 9th to the first half of the 10th centuries CE. At that time, the integration of the Turkic culture with the culture of the dominant ethnos is noted. The final period of the existence of the Altai Turkic culture (the second half of the 10th - 11th centuries CE) within the Kyrgyz Khaganate is associated with the gradual weakening and disintegration of the power into separate principalities. This situation is evidenced in the archaeological material found in the burials of the Baltargan stage, which is characterised by the fading of the Turkic traditions with regards to the appearance and creation of the material culture.

The Turks developed stable traditions of burial rites, which demonstrate a special system of world outlook of the early Medieval nomads of Altai. A deceased person was usually laid in a shallow pit with his head to the east (towards the sunrise). One or several horses were placed next to the individual, depending on the social status of the buried person. The grave was filled with things that were used during life –weapons, jewellery, and household and other items. A stone mound was built over it. Altai nomads in the early Medieval only rarely created large cemeteries. More often they buried deceased people near older burial mounds, emphasising their relationship to distant ancestors who lived in the area. Among the monuments, the so-called Turkic enclosures (тюркские оградки) stand out, often accompanied by sculptural monuments. The study of the Turkic culture is an important scientific direction for ethno-cultural reconstructions in the Medieval period.

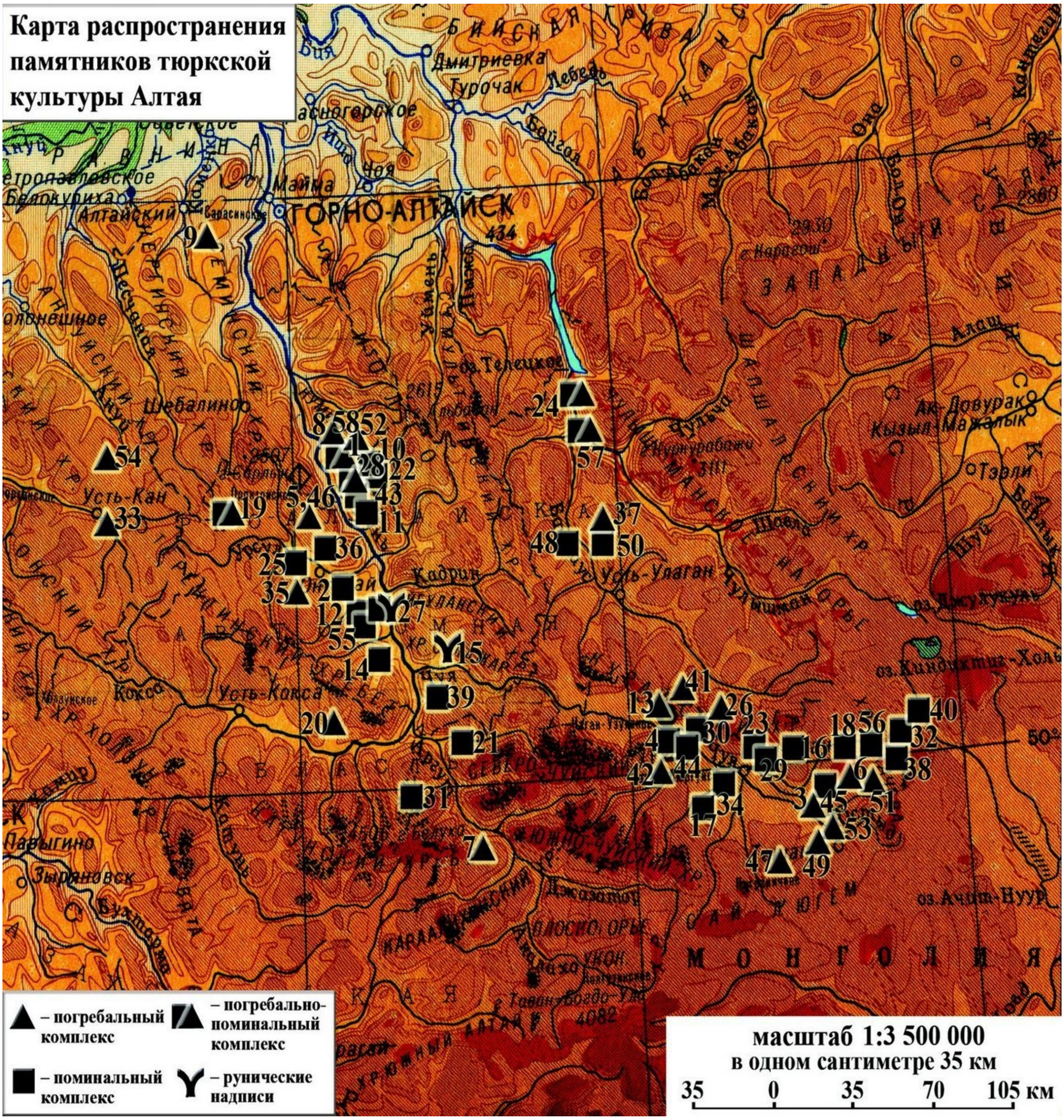


**Fig. S1.9.** Map-scheme of distribution of archaeological sites of Turkic culture in Altai (from: (*123*), figure on page 13). Triangle: burial complex, Square: memorial complex, Triangle-and-square: burial and memorial complex, Y-shaped sign: runic inscriptions.

**Kurota-I**

The site of Kurota-I is located in the Ongudai district of the Altai Republic (Russia), on the left bank of the Kurota River, approximately 2 km from its confluence with the Ursul River (50.814984 N, 85.973646 E). In 1937, one kurgan of the Turkic culture was excavated at the Kurota-I burial ground. In 1977, an expedition of the Institute of Archaeology of the USSR Academy of Sciences conducted research there, opening **Kurgan No. 2**. Under a small stone kurgan mound there was a disturbed grave, in which the remains of a burial of a young male (16-18 years old) accompanied by a foal were found. The deceased male was lying in the northern part of the pit supine, oriented with his head to the east. The animal burial was separated by a partition made of upright stones. No accompanying inventory was present. The results of the excavation are briefly published (*124*). In the absence of material culture objects, a broad dating of 6th to 10th centuries CE is indicated.

**Taldura-I**

The site of Taldura-I is located in the Kosh-Agach District of the Altai Republic (Russia), approximately 5-6 km west of the village of Bel`tir, on the right bank of the Taldura River (49.963964 N, 88.140258 E). It was investigated by the archaeological detachment of the expedition of the Institute of Archaeology of the USSR Academy of Sciences in 1977.

Under the embankment of the Early Iron Age **Kurgan No. 2** an inset (secondary) burial of the Turkic period was found, distinguished by the unusual arrangement of the horse interred above the human individual. The deceased was laid in an extended supine position, with his head to the east (with a slight deviation to the north). A birch bark quiver with iron arrowheads was lying along the right leg. A bone buckle was found near the quiver, as well as fragments of an iron knife. The horse was oriented in the opposite direction. It had iron horse bits with cheekpieces and two stirrups. The excavation materials of the inset burial have been published (*125*). It is dated to the 2nd half of the 7th - 1st half of the 8th centuries CE and attributed to the Katanda stage of the Turkic culture (*126*).

**Kurgan No. 4** of the Taldura-I site is a burial site of the Pazyryk culture (*126*, *127*). It is probable that the inventory is contained in the anthropology cabinet of Tomsk State University, where osteological materials from the mentioned barrow are kept. The available skull was erroneously defined as belonging to the Medieval period. A similar situation applies to Kurgan No. 3.

**Ust`-Biyke-III**

The Ust`-Biyke-III kurgan cemetery is located in the Chemalsky district of the Altai Republic (Altai), approximately 0.4 m above the confluence of the Biyke River (51.10243 N, 86.9387E). In 1997, the archaeological team of Altai State University investigated **Kurgan No. 6** (*128*). Beneath a small stone covering was a relatively large grave pit with a partition made of large stones. In the northwestern part of the grave there was a recess in which a deceased male of 55-60 years old with his legs flexed at the knees, supine, with his head to the northeast, had been laid (Fig. S1.10). Three iron arrowheads, bone plates from a bow, two iron buckles, a stone whetstone, a vertebra of an unreported animal, and a fragment of an iron knife were found. In the southeastern part of the grave, there was a step structure void of material remains. It appears that this part of the pit was intended for a horse, but no animal was interred. Kurgan No. 6 is dated to the 2nd half of the 5th - 1st half of the 6th centuries CE and attributed to the early (Kyzyl-Tash) stage of the Turkic culture.

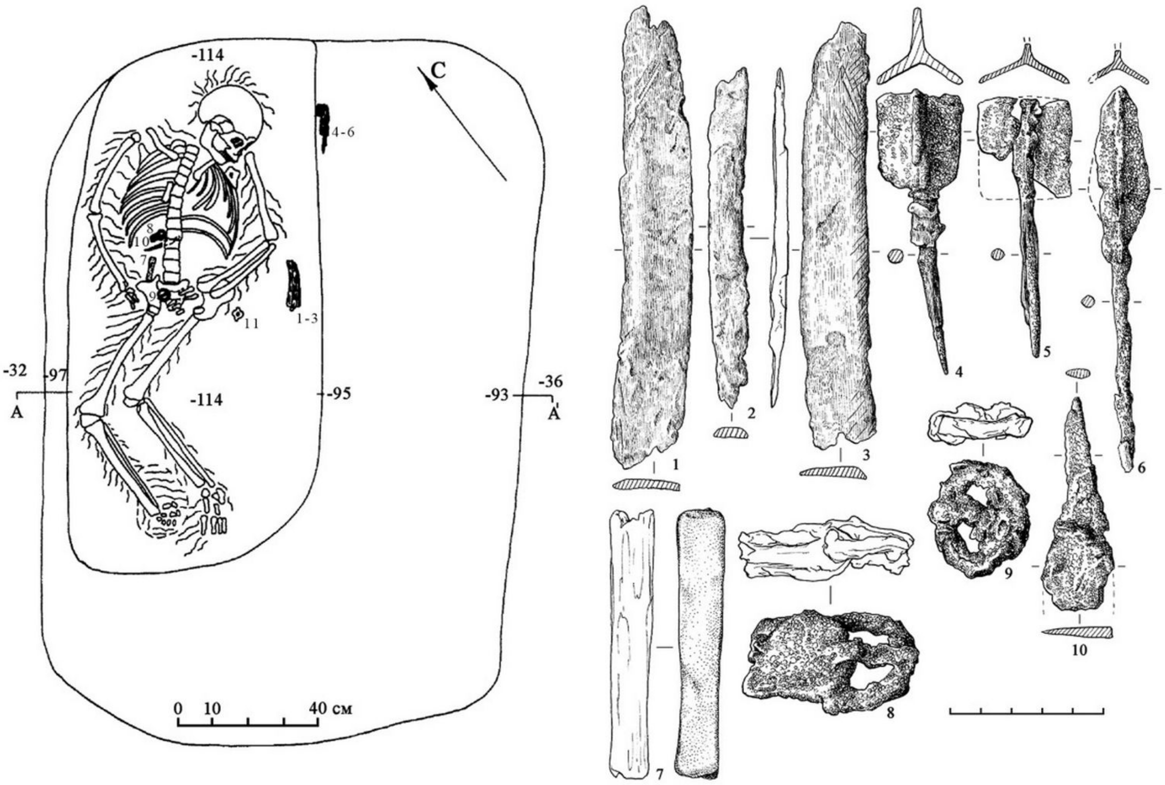


**Fig. S1.10.** Ust'-Biyke-III, Kurgan No. 6. The arrow labeled "C" (*Север*, "North") indicates the direction of the north. Findings: 1-3 - bone plates; 4-6 - iron arrowheads; 7 - stone axe; 8, 9 - iron buckles; 10 - iron knife (from (*128*)).

**Biryuzovaya Katun`-1**

The Turkic kurgan of Biryuzovaya Katun`-1 (*129*) was located in the Altai Krai (Russia), near the Bolshaya Tavdinskaya Cave, on the left bank of the Katun` River (51.492885 N, 85.461490 E). Work at the site was carried out in the summer of 2005 by Altai State University. Beneath a small kurgan mound composed of round river pebbles, there was a grave pit reaching up to 0.7 m deep. The burial of a male aged 50-65 accompanied by a horse was found inside (Fig. S1.11). The male was lying stretched out on his back and had his head oriented northeast-east, while the horse was oriented in the opposite direction. The male edentulous maxilla and all alveoli were remodeled due to antimortem tooth loss. The burial inventory included the following finds: four fragmented arrowheads, a large adze, a small iron tool, a knife, a stone grinding set (saddle quern with its hand-stone) (Fig. S1.11), and belt decorations of an archer's belt and main belt were found. In addition, there was a cluster of millet seeds, which were apparently placed in a leather pouch. The horse skeleton was examined and found to contain a horse bit with cheekpieces, two iron stirrups, and a horn cinch buckle. Based on the analysis of the material culture, the mound of Biryuzovaya Katun`-1 is dated to the second half of the 7th - first half of the 8th century, to the time of the Second Eastern Turkic Khaganate (682-744 CE).

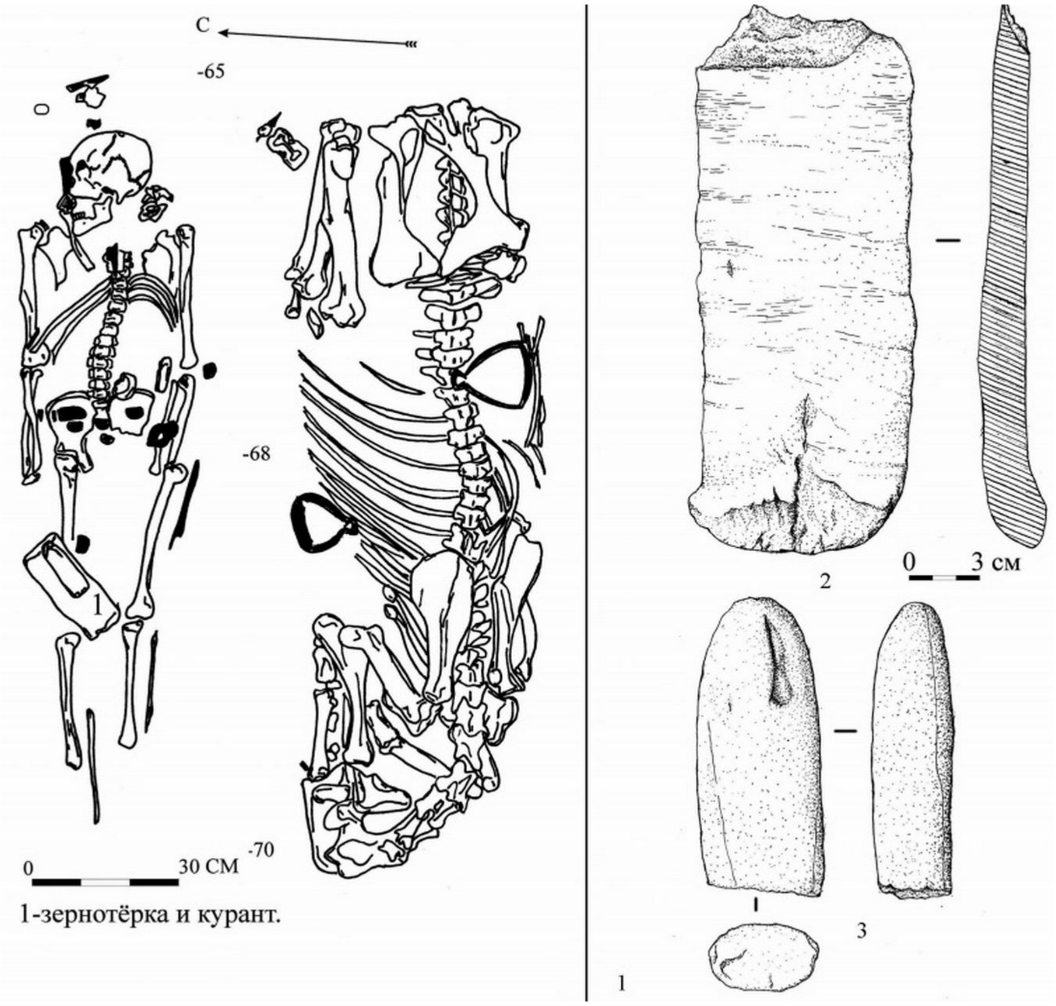


**Fig. S1.11.** Biryuzovaya Katun`-1. The arrow labeled "C" (*Север*, "North") indicates the direction of the north. On the left - burial of a male with a horse; on the right - 1, 2, 3 - stone grinding set (from (*129*))

**Biryuzovaya Katun`-3**

The single Turkic kurgan of Biryuzovaya Katun`-3 (*129*, *130*) was located in the district of Altai Krai (Russia), approximately 4.5 km from the mouth of the Tavdushka River up the left bank of the Katun`, and 0.15 km southeast of the road running along the coastal zone of the tourist complex 'Biryuzovaya Katun`' (51.484805 N, 85.453379 E). Archaeological work at the site was carried out in the summer of 2006 by Altai State University. Beneath a small kurgan mound with a crepida of large stones, there was a grave pit, in which there was a single burial of an adult male, laid stretched out on his back, supine, with his head to the southeast (Fig. S1.12). The following accompanying inventory was found: 14 iron arrowheads, elements of an archer's belt, the garniture of a basic belt set, a bronze earring, and various iron items (a ring, an adze, a woodworking tool, a knife). Judging by the traces of burial rites and the composition of the grave good assemblage, the kurgan in question belongs to the Turkic period. The time of its construction can be determined quite confidently by a fairly typical set of items, especially the belt assemblage. Close analogies allow us to date the investigated object within the 2nd half of the 7th – 1st half of the 8th centuries, which corresponds to the Katanda stage of the Altai Turkic culture. This burial marks the northern border of the known Turkic inhabitance in the Altai during the Second Eastern Turkic Khaganate (682-744 CE).

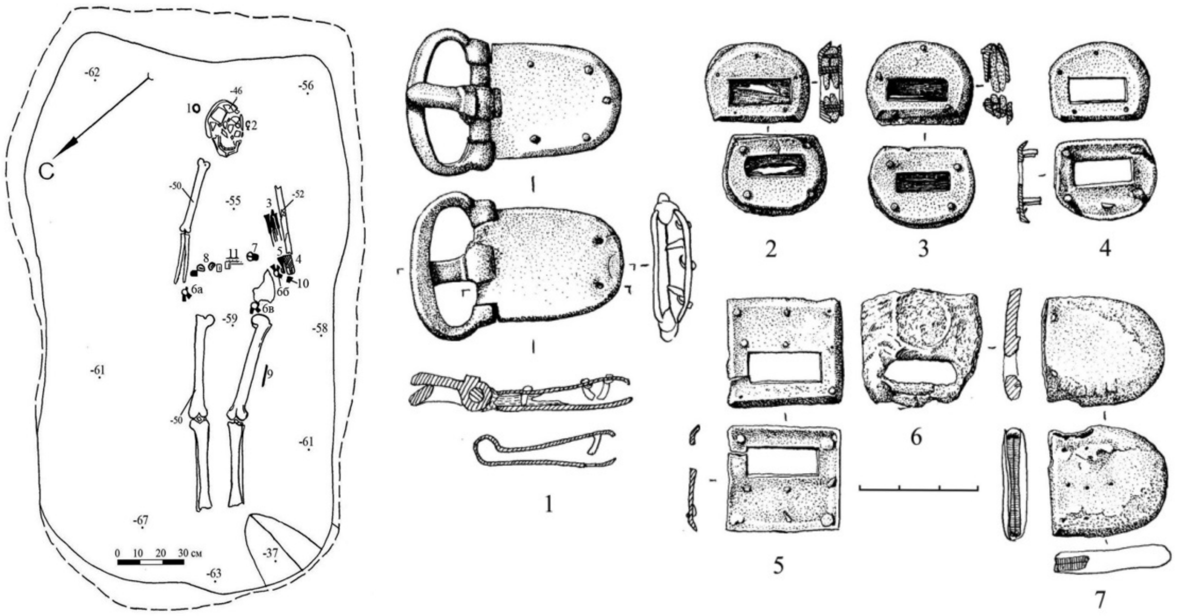

**Fig. S1.12.** Biryuzovaya Katun`-3. The arrow labeled "C" (*Север*, "North") indicates the direction of the north. Human burial and details of a belt set of the Turkic period (from: (*129*), Figs. 22, 24).

**Biryuzovaya Katun`-9**

The Turkic burial kurgan of Biryuzovaya Katun`-9 (*129*, *131*) was located in Altai Krai (Russia), directly at the foot of the Bolshaya Tavdinskaya Cave, on the left bank of the Katun` River (51.463982 N, 85.435571 E). Work at the site was carried out in the summer of 2009 by Altai State University. Beneath a small kurgan mound with a crepida built of large stones, there was a grave pit, at the bottom of which, there was a skeleton of a male aged 50-55 years, and next to it a complete horse skeleton. The deceased male was lying in left lateral decubitus, crouched (with his legs bent and tucked), and his head to the southeast. He was found with bone plates for a bow, five iron arrowheads, a knife, remains of clothing, an archer's belt with a buckle and three strap distributors, and a basic leather belt with a buckle and eight overlay plaques (Fig. S1.13). Additionally, the vertebral column of a sheep was found. The horse equipment is represented by a bit with cheekpieces, one iron stirrup, a cinch buckle and three saddle spring buckles (Fig. S1.13). The excavated objects can be attributed to the time of the Second Eastern Turkic Khaganate and dated to the second half of the 7th century.

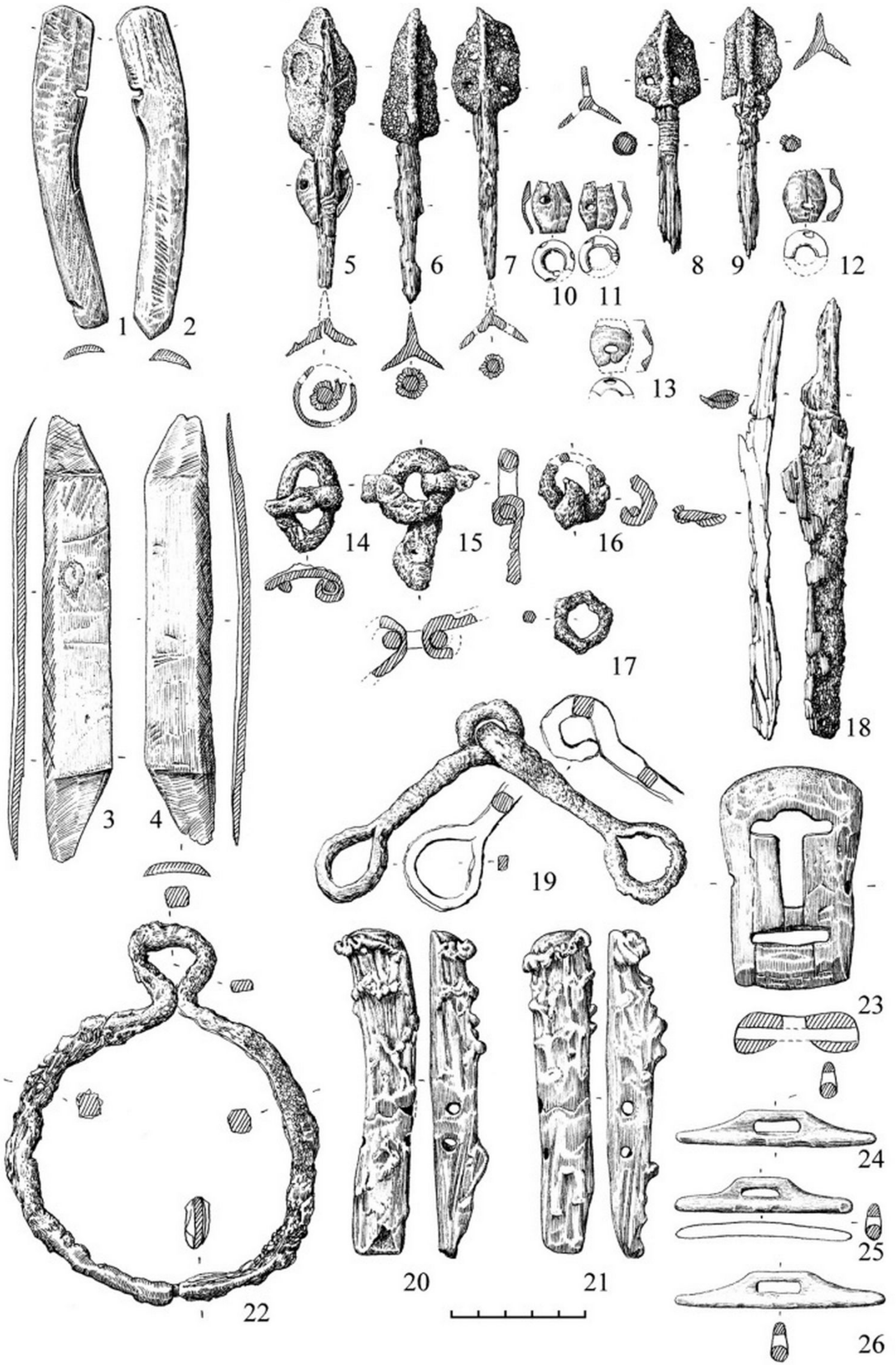


**Fig. S1.13.** Biryuzovaya Katun`-9. Inventory from the mound. 1-4, 10-13, 20-21, 23-26 - horn; 5 - iron, wood, horn; 6-9, 18 - iron, wood; 14-17, 19, 22 – iron (from: (*129*), Fig. 40)

**Shibe-II**

The Shibe-II kurgan cemetery (*132*) is located in the Ongudai District of the Altai Republic (Russia), on the left bank of the Ursul River, approximately 100-150 m north of the nearest outskirts of the village of Shiba (Shibe) and 250 m west of the Great Shibe kurgan, which was excavated in 1927 (50.511906 N, 85.448380 E). In 1986, Altai State University conducted research at the Shibe-II site (*133*). As a result, the burial kurgans of the Turkic period were studied. The obtained materials have not been published in full. Based on the available sources (report, field documentation and collection), it is possible to give a brief characterisation of the result of the research of kurgans Nos. 4, 11 and 12, from which the samples taken for genetic analysis originate. These kurgans are dated to the 2nd half of the 7th - 1st half of the 8th centuries CE and belong to the Katanda stage of the Turkic culture (*126*).

Under a small stone embankment of **Kurgan No. 4** there was a grave pit, at the bottom of which a burial of an individual with a horse was found (Fig. S1.14). The human skeleton lay 10 cm above the horse's skeleton, behind a partition of four upright stones. The deceased female, aged 20-25 years old, was lying on her back, with her legs slightly bent at the knees, and her head oriented to the southeast. At her left foot lay the bones of a sheep. The individual had no accompanying grave goods. The horse was positioned along the southwest wall and oriented in the opposite direction. It was bridled and saddled. This is evidenced by iron finds (a horse bit with cheekpieces, a rim of the front bow of the saddle, two stirrups, a cinch buckle), as well as bronze bridle ornaments (plaques).

**Kurgan No. 11** also had a mound of stones, under which there was a burial of a male with a horse. The deceased male, aged 25-30 years, was lying on a 25 cm high ledge in an extended position, supine, with his head to the east. The horse was oriented in the opposite direction. In the grave, there were parts of bone plates on a bow; a birch bark quiver with 10 iron arrowheads and fragments of shafts; a bronze buckle; an oxide from a silver piece; a sheep skeleton (under birch bark); fragments of iron bridles; bridle plaques (2 pieces); fragments of two stirrups; a bone buckle (beneath the quiver); bone fetters; and an iron knife.

In **Kurgan No. 12**, a deceased accompanied by a horse was buried, the skeleton of which was on a 15 cm high ledge and behind a partition made of stones. Numerous charcoals were found in the grave fill and at the bottom of the grave. The deceased female, aged 40-45 years, was lying in an extended position, supine, with her head facing east-northeast. The person had no accompanying inventory. The animal was oriented in the opposite direction. Among the bones of its skeleton the following finds were noted: remains of a bridle, the straps of which were decorated with plaques; iron horse bit; fragments of a stirrup; a bone buckle; and fragments of an iron object of unclear purpose.

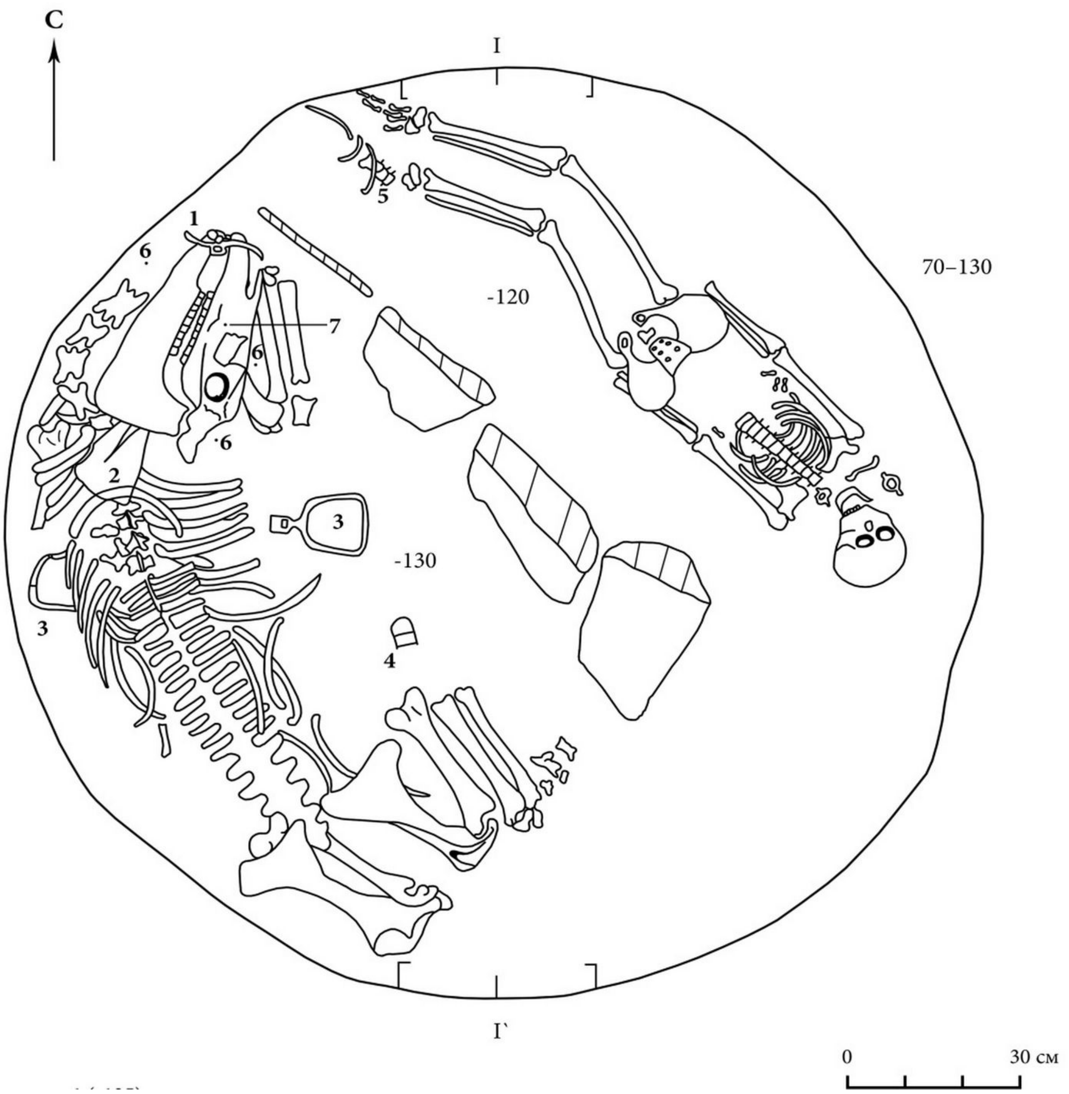


**Fig. S1.14.** Shibe-II. Kurgan No.4. Burial of a female with a horse: 1 - iron bit with cheek pieces, 2 - iron lining of the front saddle bow; 3 - iron stirrups; 4 - iron girth buckle; 5 - sheep bones; 6 - bronze plaques (from: Archive of IA RAS. R-1. No. 11267). The arrow labeled "C" (*Север*, "North") indicates the direction of the north.

**Katanda-I/3**

The archaeological complex Katanda-I (*134*, *135*) is located near the village of the same name, in the Ust-Koksinsky district of the Altai Republic (Russia), on the right bank of the Katanda River, a left tributary of the Katun (50.122074 N, 86.83284 E). Under the leadership of V.V. Radlov in 1865, burial mounds of the Turkic time were excavated, the materials of which are only partially published. In 1984, Altai State University investigated 10 burials of the Turkic culture (Kurgans No. 1-7, 11, 16, 21), as well as a 'ritual' kurgan (No. 22). When publishing the results of the excavations, the authors labelled this burial ground as Katanda-3. Therefore, the designation Katanda-I/3 is used in this article. Genetic analysis of three samples from Kurgans No. 3, 6 and 16 was carried out; they were composed of stone mounds, under which burials of people accompanied by a horse were found in the graves (Fig. S1.15). The time of construction of the presented burial mounds is determined by the 2nd half of the 7th - 1st half of the 8th centuries and corresponds with the period of the Second Eastern Turkic Khaganate (682-744 CE).

In **Kurgan No. 3**, the grave was partially looted. Judging by the preserved bones, the male was lying in an extended supine position, with his head to the southeast, while the horse was oriented in the opposite direction. The animal was perched on a small ledge and was separated by a partition made of two large stones. Two earrings were found. An iron buckle was found among the bones, as well as seven non-ferrous metal belt plaques. A horse was buried with an iron buckle and seven belt plaques made of non-ferrous metal.

In **Kurgan No. 6**, the grave was also disturbed by looting activities. It is likely that the male was lying stretched out on his back, in an extended supine position, with his head to the north-northeast, while the horse was oriented in the opposite direction. A ‘necklace’ of fish vertebrae and maral teeth were found next to the cranium. Two buckles, horse’s cheekpieces and two clasps made of horn, as well as an iron nail, were found from horse equipment.

In **Kurgan No. 16**, the grave was undisturbed. A male was in an extended supine position with his head oriented east-southeast. One arm was bent and the hand was located in the area of the pelvic bones. A ceramic disc (part of a spindle), an earring and a pendant from a second earring were found in the area of the cranium. An iron knife was also associated with the individual, near the right hand. The horse was lying on a small ledge. It was accompanied by a bit with sidebars, two strap distributors, a bridle buckle, a horse’s forehead ornament, two stirrups and a cinch buckle. An iron dagger lay separately in the grave.

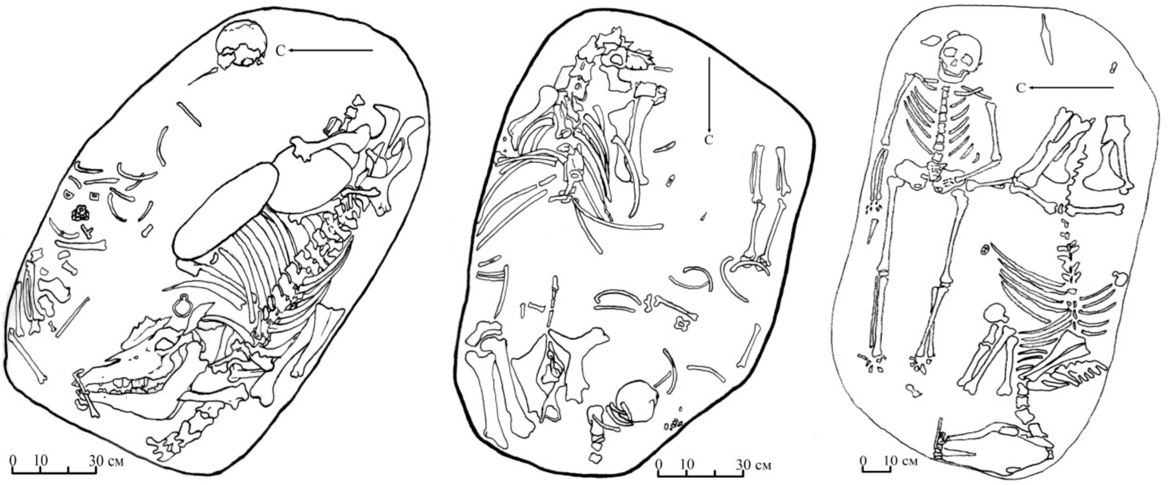


**Fig. S1.15.** Katanda-I/3. Investigated graves of Kurgans No. 3, 6 and 16 (according to: (*135*), Figs. 4, 5, 7). The arrow labeled “C” (*Север*, “North”) indicates the direction of the north.

**Kyrlyk-II**

The Kyrlyk-II kurgan cemetery is located in the Ust-Kansky District of the Altai Republic (Russia), on the right bank of the Kyrlyk River, a left tributary of the Yabagan River flowing into the Charysh River, approximately 4 km south-southeast of the village of Kyrlyk, in a ploughed field (50.47984 N, 84.585035 E). In 1983, Altai State University excavated **Kurgan No. 3**, one of the 27 originally visible mounds. It was a small mound with a ring of larger stones (*crepida*) around its perimeter. The grave contained a burial of an individual with a horse. The skeleton of the individual lay in a recess in the bottom of the pit, along the northern wall, in an extended supine position, with their head to the east. Sheep bones were found next to the right leg. There were no accompanying grave goods near the inhumed individual. A bridled, and most likely saddled, horse was on a step, with its head oriented to the west-northwest. Iron horse bits with cheek pieces, two stirrups, and a buckle, as well as leg bones of a ram were found. The excavation materials have been published (*136*). Kurgan No. 3 is dated to the 2nd half of the 7th - 1st half of the 8th centuries CE and belongs to the Katanda stage of the Turkic culture (*126*).

**Tumechin-II**

The archaeological complex Tumechin-II is located in the valley of the Ursul River, on the bank of the Tumechin Creek, in the Ongudai District of the Altai Republic, Russia (50.46445 N, 85.35028 E). The excavations of Kurgan No. 2, belonging to the Early Medieval period, were carried out by Yu.T. Mamadakov and V.N. Vladimirova in July 1991.

**Kurgan No. 2:** The grave, divided by a stone partition at the bottom, contained the burial of a male with his head oriented west, accompanied by a horse oriented in the same direction. The accompanying inventory included elements of the horse's equipment. The dating of the site is from the second half of the 6th - first half of the 7th centuries CE.

**Tozhon**

The archaeological complex Tozhon is located on the second terrace above the floodplain of the Kayerlyk River, approximately 1.5 km east of the village of Elo, Ongudaysky District, Altai Republic, Russia (50.770474 N, 85.573964 E). The excavation of Kurgan No. 1, belonging to the Early Medieval period, was carried out by Yu.T. Mamadakov and V.N. Vladimirova in July 1991.

**Kurgan No. 1:** Beneath the kurgan was a partially looted grave, which contained the bones of a male and a horse. The deceased individual had his head oriented to the east; the orientation of the horse was not established. The surviving accompanying inventory included iron stirrups, two arrowheads, a horn on the bow, and part of a buckle. The site is dated to the second half of the 7th-8th centuries CE.

**Tytkesken`-VI**

The well-known archaeological complex Tytkesken`-VI (*137*, *138*) is located in the Chemalsky District of the Altai Republic (Russia), on the left bank of the Katun River, near the confluence of the Tytkesken` River, approximately 1.5 km south of the village of Elanda (51.12635 N, 86.45494 E). Excavations at the site were carried out by Altai State University in 1988-1989. Three Turkic-period kurgans (No. 1, 5 and 10) were investigated. They were located to the east of the Pazyryk-culture burial structures, on the edge of the terrace.

In **Kurgan No. 5** at the bottom of the grave pit, was the skeleton of an adult male, who was extended and supine, with his head to the northeast-east (Fig. S1.16). There were two stones near the cranium, as well as two iron arrowheads of poor preservation. Another stone stood at the feet of the buried male. In the other half of the grave was a horse skeleton. Remains of an iron horse bit with horn cheekpieces and one stirrup were found on it. The horse was oriented in the opposite direction to the male. In addition, a clasp for a harness and remains of birch bark were found. The investigated kurgan was dated to the second half of the 5th - first half of the 6th centuries and is attributed to the earliest (Kyzyl-Tash) stage of the Turkic culture (*126*).

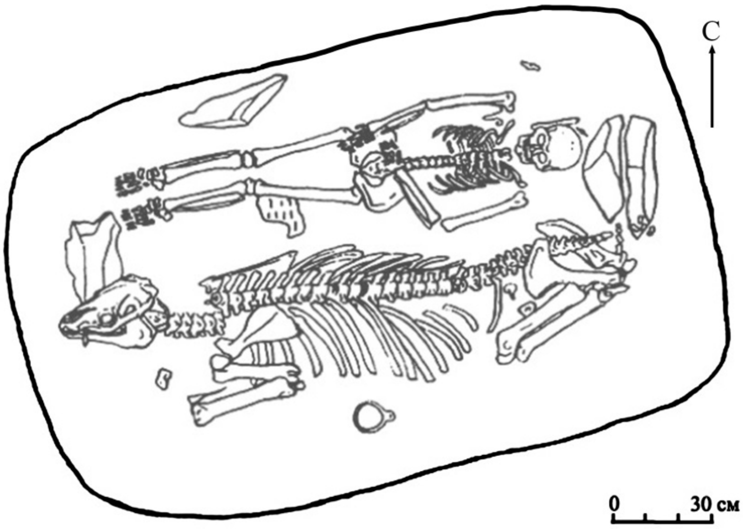


**Fig. S1.16.** Tytkesken`-VI. Mound No. 5. Burial of a male with a horse (from: (*137*), Fig. 4: 1). The arrow labeled “C” (*Север*, “North”) indicates the direction of the north.

### d. *Srostki culture*

In the middle of the 8th century CE, the political situation in the south of Western Siberia changed, as a significant part of the Turks from Altai advanced northwards and subdued the local Samoyedic population. The Turks occupied the Ob plateau, with the adjacent lands of the Kulunda steppe and the right-bank floodplain in the upper reaches of the Ob. As a result of this interaction, a new community exhibiting significant social stratification emerged, in which the Turks occupied a dominant position, and the Samoyeds a relatively subordinate one. In archaeology this phenomenon is considered to be the start of the Srostki culture (*30–32*), which received its name after the designation of the base necropolis Srostki-I. The sites of the Srostki culture (kurgan and flat-ground burial sites, settlements and hillforts) are evidenced across a large territory in the south of Western Siberia by the 2nd half of the 8th - 12th centuries CE (Fig. S1.17). The Turkic-Samoyed population lived in the forest, steppe, and foothill zones, in close proximity to sources of water. Settlements are found near river and lake floodplains, with most of them located on hills (елбан/elban) or cape-shaped coastal terraces.

The tribal economy was complex, but was mainly composed of semi-nomadic pastoralism. Agriculture also constituted an important branch of the economy. Despite the development of productive economies, hunting, fishing, and gathering continued to represent important elements of subsistence. A wide range of handicraft production and extensive trade relations are also attested. Among the settlements, a distinct subset is represented by those with advanced fortifications, including refuge-fort settlements and large central fortified centers. Excavations at more than 125 funerary monuments have resulted in an extensive source base, allowing researchers to distinguish four developmental stages within the Srostki culture, each corresponding to known historical events of the Early Medieval period. These stages are as follows:

1. The Inya stage (second half of the 8th - first half of the 9th centuries CE) marks the formation of the Srostki culture and is named after the type-site Inya-1. Its beginning is connected with the collapse of the Second Eastern Turkic Khaganate in 744 CE and the subsequent migration of some Turkic tribes into southern parts of Western Siberia.

2. The Gryaznovo stage (second half of the 9th - first half of the 10th centuries CE) reflects the completion of the process of consolidation of the Srostki tribes and the formation of distinctive features within their culture. Its beginning is contemporary with the military expansion of the Kyrgyz Khaganate from 840 to 847 CE, and the formation of the Kimak power around 850 CE.

3. The Shadrintsevo stage (second half of the 10th - first half of the 11th centuries CE) represents the period of prosperity in the Srostki culture. Coinciding with the campaigns of the Liao Empire in Mongolia of 924-994 CE, which was founded by the Mongol-speaking Khitans, a number of Turkic-speaking tribes were displaced to the west. The Kyrgyz and Kimak Khaganates were additionally weakened.

4. The Zmeyevo stage (2nd half of the 11th - 12th centuries CE) corresponds to the gradual decline of the Srostki groups. The beginning of this process coincided with new migrations of nomadic tribes from Inner Asia, which contributed to the collapse of the Kimak Khaganate and the strengthening of the Kipchaks, who occupied part of the western area of the Srostki culture.

In the beginning of the 13th century (1207 CE), the Mongols conquered the territory of the forest-steppe Altai, marking a major political and cultural transformation in the region. As a result, a new community was formed, called the Karmatsky culture, which largely consisted of the descendants of the Srostki population.

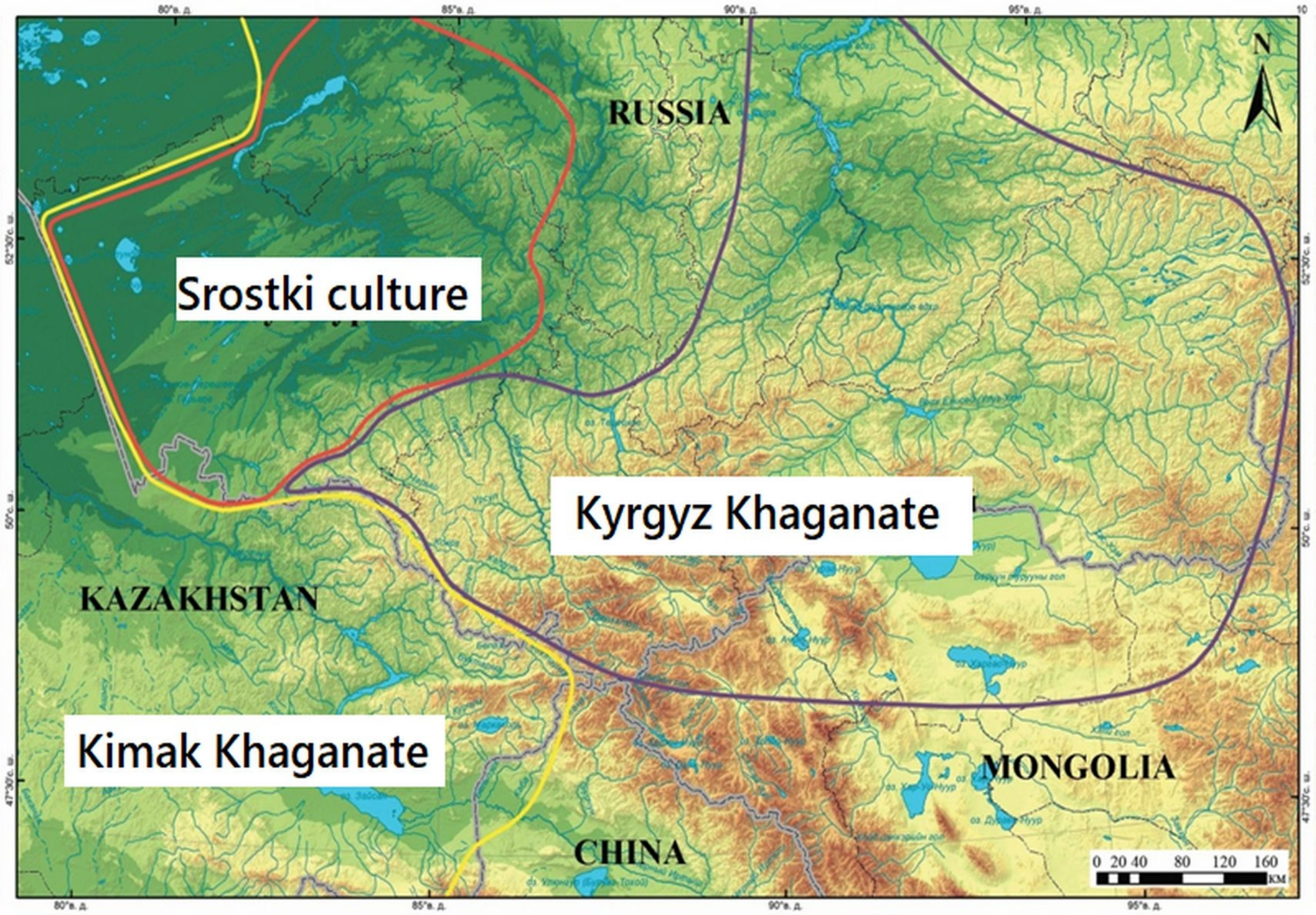


**Fig. S1.17.** The area of distribution of archaeological sites of the Srostki culture in the 10th century CE (adapted from: (*30*), Fig. 3.32).

**Inya-1**

The Inya-1 kurgan cemetery (*32*, *139–143*) is located in Shelabolikhinsky District, Altai Krai (Russia), 0.8 km southwest of the village of Inya, on the terrace of the left bank of the Inya River, a right tributary of the Ob (53.294184 N, 82.383724 E). The site consists of 37 earthen kurgans ranging from 5–12 m in diameter, and 0.1–0.7 m high (Fig. S1.18). A total of 29 kurgans were excavated, containing 52 burials with a wide range of mortuary rites including inhumations with a horse, dog, sheep, or cow; single-human inhumations; cremations; and cremations accompanied by a horse. Among the findings, iron arrowheads, swords, lamellar armour plates, daggers, knives, adzes, horse's forehead ornaments, birch-bark quivers, antler bow plates, buckles, fasteners, bronze buckles, strap-ends, strap-ends and distributors of straps from bridles and belts, earrings, rings, pendants, coins, and ceramic vessels were present (Fig. S1.18). Additional finds, not shown in the figure include horse bits with cheekpieces, stirrups, bracelets, piercings, and beads. The investigated kurgans in this study, numbered **20**, **23**, and **26**, date from the second half of the 8th to the first half of the 9th century CE and reflect the earliest stage of the Srostki culture, the Inya.

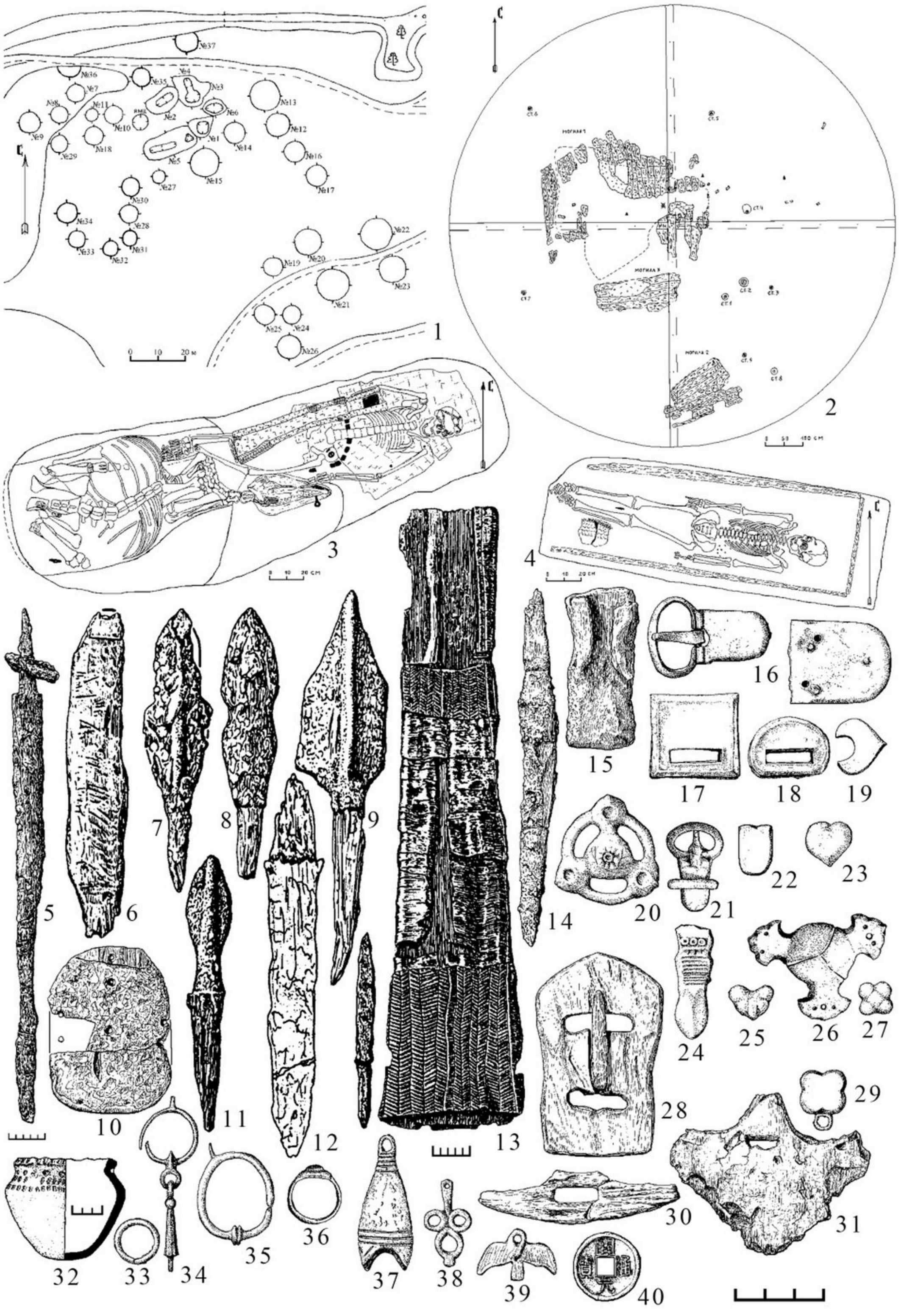

**Fig. S1.18:** Kurgan burial mound Inya-1: 1 - site plan, 2 - plan of the barrow, 3-4 - plans of graves, 5-40 – inventory (from: (*32*), Fig. 26). The arrow labeled “C” (*Север*, “North”) indicates the direction of the north.

### Ekaterinovka-3

The kurgan cemetery Ekaterinovka-3 (*32*, *144*) is located in the Kulunda district of Altai Krai (Russia), 0.8 km east of the village of Ekaterinovka, within the eastern part of the Kulunda Plain (52.34454 N, 79.374962 E). The 1988 excavation conducted by Altai State University uncovered the site, which consisted of five earthen kurgans ranging from 16–28 m in diameter, and 0.5–0.94 m high. Five burials were found according to the rites of single and double inhumation, collective inhumation with a horse, and a cenotaph with a horse. Among the objects found there were present bronze buckles, plaques from bridles, belts and bags, an earring, pendants, a mirror, and a gilded plaque in the form of a lion or dog figure (Fig. S1.19). Fragments of iron horse bits with cheekpieces, a cauldron, and fragments of leather straps from bridles were also found. The necropolis is dated to the second half of the 9th – first half of the 10th centuries CE.

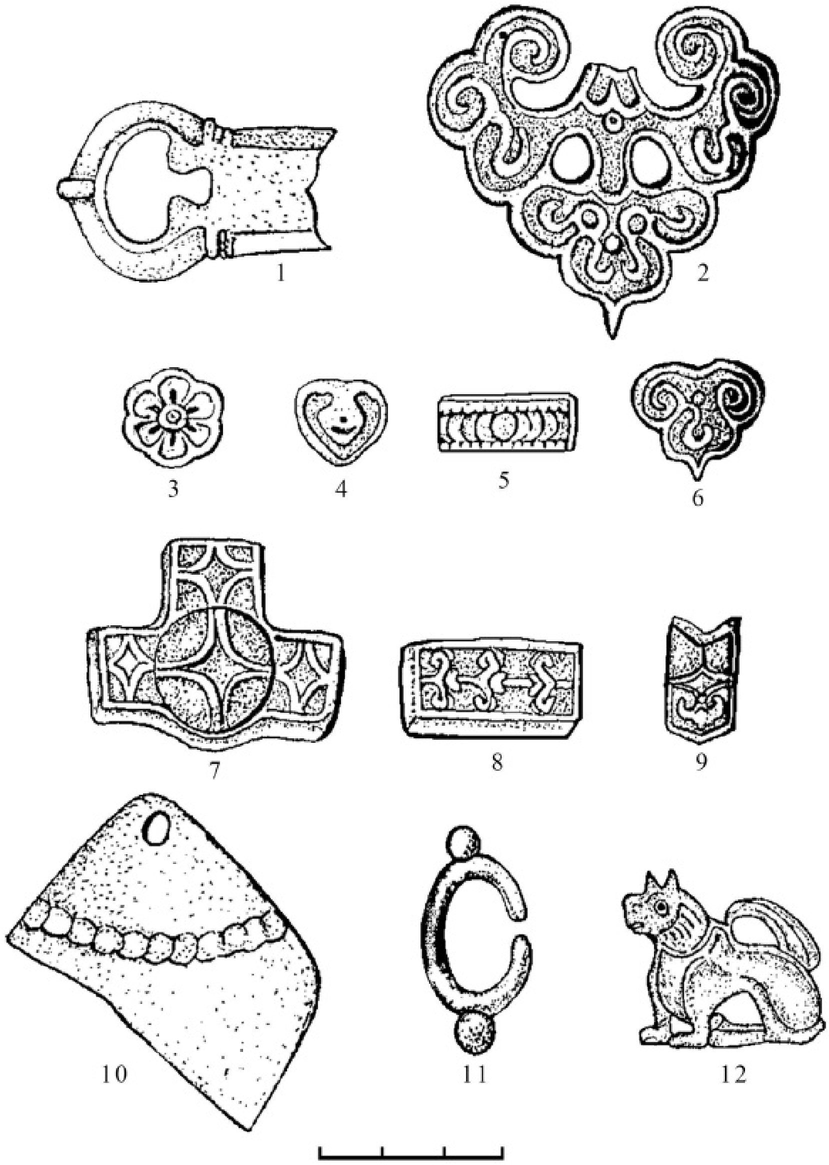


**Fig. S1.19:** Metal finds from the burial mound Ekaterinovka-3 (from: Gorbunov, Tishkin, 2025. Fig. 25: 8–19).

**Shchepchiha-I**

The Shchepchiha-I kurgan cemetery (*32*, *145*) was located 2.5–3.5 km from the village of Lokotok, Zmeinogorsky District, Altai Territory (Russia), on the crest of an interfluve ridge and a hill between the Blizhnyaya Shchelchikha and Dal'nyaya Shchelchikha Rivers (51.14333 N, 81.473254 E). In 1991, the archaeological expedition of the Altai State University excavated all identified kurgans at the site. Three of them (Nos. 3, 4 and 6) yielded extensive archaeological material relating to the Early Middle Ages (Fig. S1.20). These finds include items of armament, horse equipment, jewellery, tools, and household items. They can be correlated with the material culture of the Shadrintsevo stage (second half of the 10th – first half of the 11th centuries) of the Srostki culture.

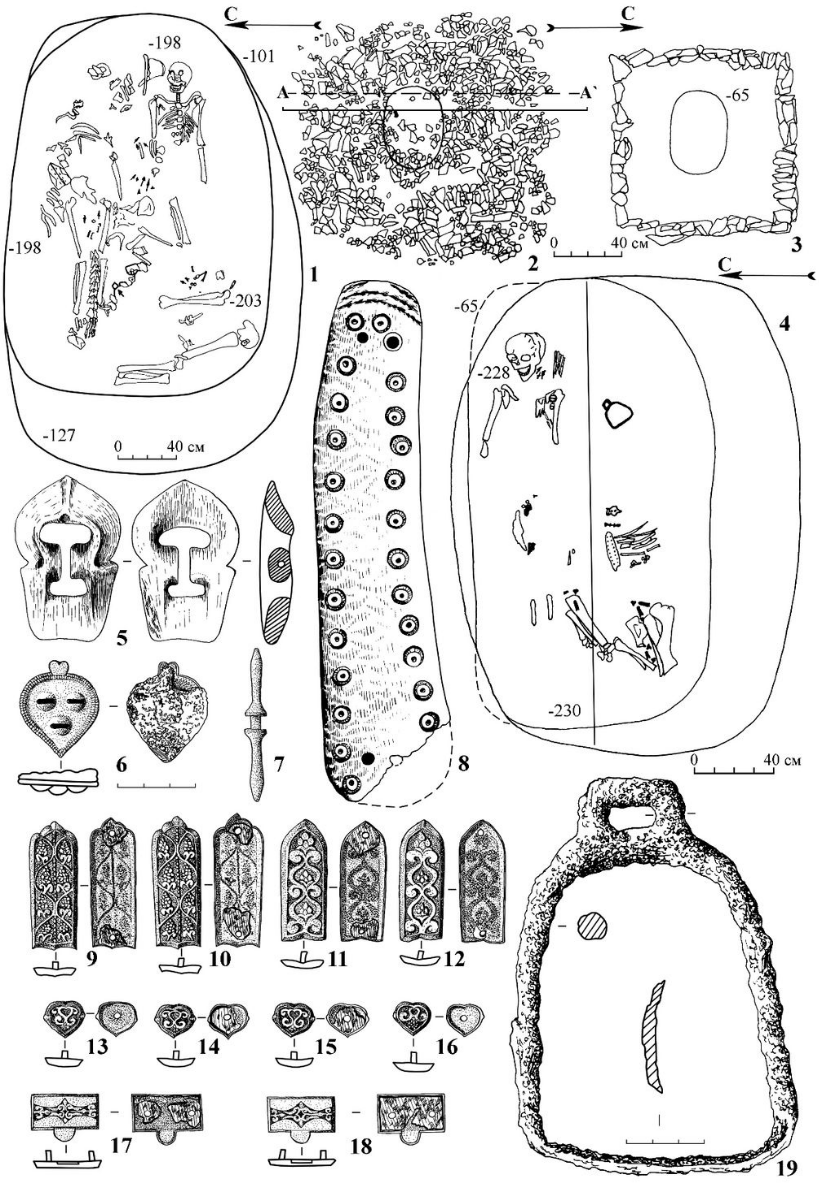


**Fig. S1.20:** Kurgan cemetery Shchepchiha-I: 1-4 – plans of graves and barrow, 5-19 – finds (from: Gorbunov, Tishkin, 2025. Fig. 60). The arrows labeled “C” (*Север*, “North”) indicate the direction of the north.

**Borovikovo-IV**

The Borovikovo-IV kurgan cemetery (*31*, *32*, *146*) is located in Pavlovsky District, Altai Krai (Russia), 0.5 km northeast of the village of Borovikovo, at the base of a promontory-like protrusion of a fluvial terrace of the left bank of the Ob River, within the Ob Plateau (53.254667 N, 82.52029 E). It was investigated in 2004 by the archaeological expedition of the Altai State University. The site consisted of two earthen kurgans, each 10–11.5 m in diameter, and up to 0.4–0.5 m high (Fig. S1.21). **Kurgan No. 1**, which is dated to the second half of the 11th-12th centuries CE, contained two wooden posts and two graves, referred to as Grave-1 and Grave-2 (Fig. S1.21).

**Grave-1** consisted of a wooden superstructure and an earth-cut grave pit, and appeared to have been looted. The pit fill contained mixed bones from the skeleton of a mature male, over 55 years old, and animal hindquarters. Iron objects (a knife, a saddler's awl, a bridle, a girth buckle), as well as items made of bone and horn (a bow plate, an arrowhead, and a belt buckle with an iron tongue) were found in connection (Fig. S1.21). Beneath the collapsed wooden cover, the partial skeleton of a 14–18 year old, young female with her head oriented to the northeast was preserved. Three glass beads were found in connection with this context (Fig. S1.21). Remains of a wooden frame were recorded on the sides of the grave pit.

**Grave-2** was located 3.6 m north-northeast of Grave-1, within Mound 1, and consisted of a multi-individual pit grave. At the bottom of the pit were the skeletal remains of three subadults lying supine in a row with heads oriented to the northeast (Fig. S1.21). No grave goods were found within the burial.

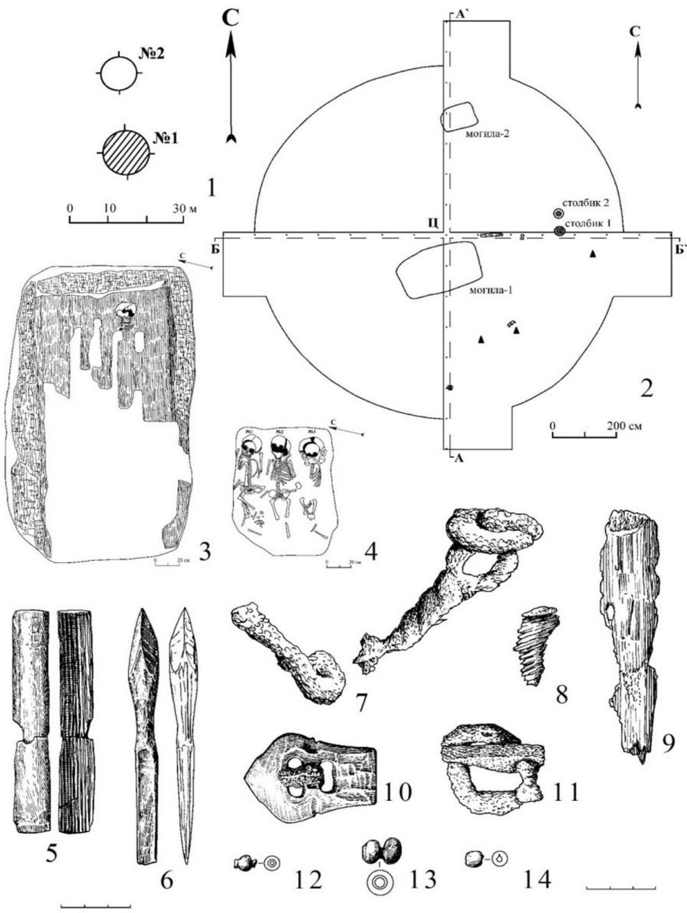

**Fig. S1.21:** Borovikovo-IV kurgan cemetery: 1 - plan of the site, 2 - plan of the barrow, 3-4 - plans of graves, 5-14 – finds (from: Gorbunov, Tishkin, 2025. Fig. 8). The arrows labeled “C” (*Север*, “North”) indicate the direction of the north.

**Chernyi Kamen'-I**

The Chernyi Kamen'-I kurgan cemetery (*32*, *147*), consisting of 12 kurgans, is located 3 km south-southeast of the village of VI Congress (VI Конгресс), Rubtsovsky District, Altai Krai (Russia), on the left bank of the Ustyanka River, on the northeastern slope of a small hill (51.165462 N, 81.320636 E). Six kurgans were excavated by an archaeological expedition of the Altai State University in 1990. All of the kurgans had sustained heavy damage by looters (Fig. S1.22), nevertheless the osteological remains of both humans and horses were recovered. The material finds include a fragment of an iron figure-eight-shaped stirrup (Fig. S1.22), as well as fragments of pottery. The site is likely attributable to the Gryaznovo stage of the Srostki culture, dating within the second half of the 9th – first half of the 10th centuries CE.

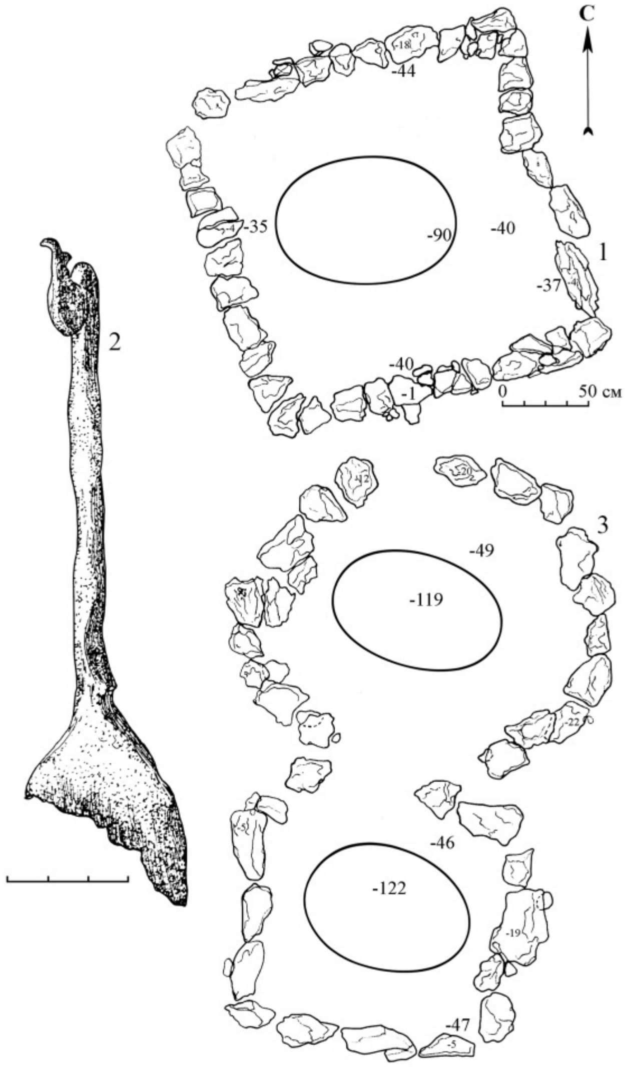


**Fig. S1.22:** Kurgan burial mound Chernyi Kamen'-I: 1, 3 - plans of stone structures and location of graves; 2 - fragment of an iron stirrup (from: (*147*)). The arrow labeled “C” (*Север*, “North”) indicates the direction of the north.

**Borovikovo-V**

The Borovikovo-V flat-ground burial (*32*, *148*) is located in the Pavlovsky district of Altai Krai (Russia), 1 km east of the village of Borovikovo, on the edge of a promontory of a fluvial terrace on the left bank of the Ob River, within the Ob Plateau (53.255022 N, 82.543194 E). In 2004, during the work of the archaeological expedition of the Altai State University, a partially collapsed burial was discovered during the inspection of the bank edge. Under a cover of timber logs there was a burial of a male laid supine, oriented with his head to the northeast-east (Fig. S1.23). The deceased was accompanied by grave goods including a bone buckle, iron bits, and a knife (Fig. S1.23). The grave is dated to the second half of the 11th – 12th centuries CE.

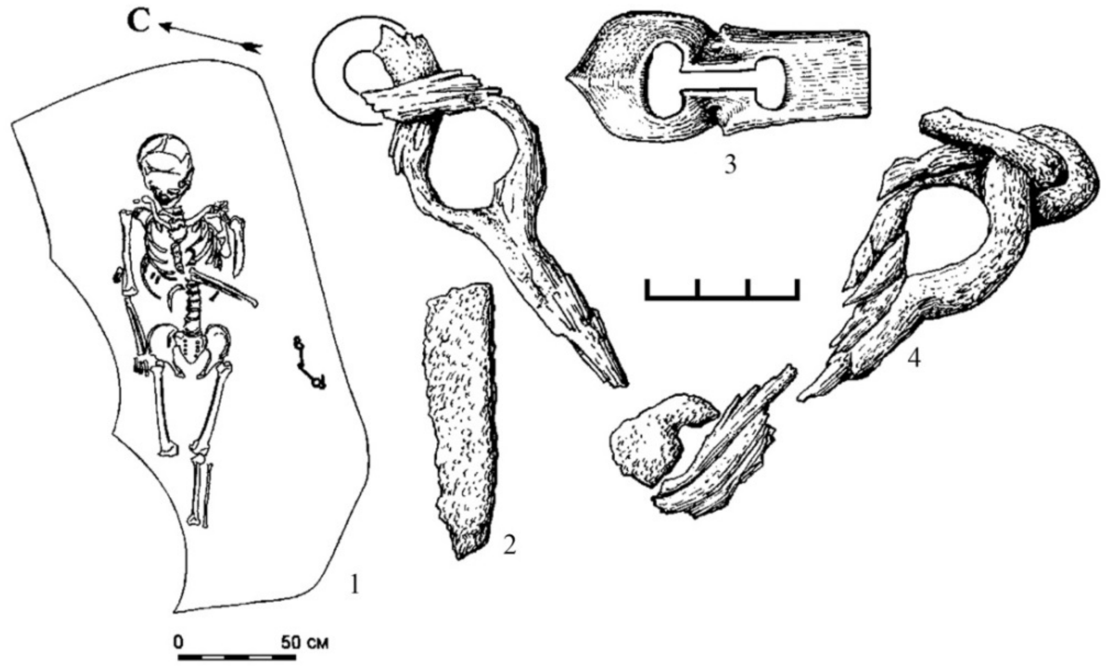


**Fig. S1.23:** Flat-ground burial Borovikovo-V: 1 - grave plan, 2-4 – finds (from: (*32*), Fig. 64: 15–18). The arrow labeled “C” (*Север*, “North”) indicates the direction of the north.

**Krasnaya Gorka-I**

A destroyed burial was discovered at the Krasnaya Gorka-I site (*149*), located 3 km northeast of the village of Novoskluikha, Rubtsovsky District, Altai Territory (Russia) and 0.12 km east of the right (high) floodplain terrace of the Skluikha River (51.325774 N, 81.260389 E). Previous interventions in the area had documented and collected osteological materials, several iron three-sided and three-bladed keel-shaped arrowheads, and a bronze earring (Fig. S1.24). In 2009, the area was investigated by researchers from Altai State University. During the excavations, two bronze belt strap-ends were found, as well as indeterminate fragments of iron objects and osteological material consisting of the remains of an adult male skeleton and horse bones. The burial belongs to the Srostki culture and is tentatively dated to the second half of the 9th – first half of the 10th centuries CE.

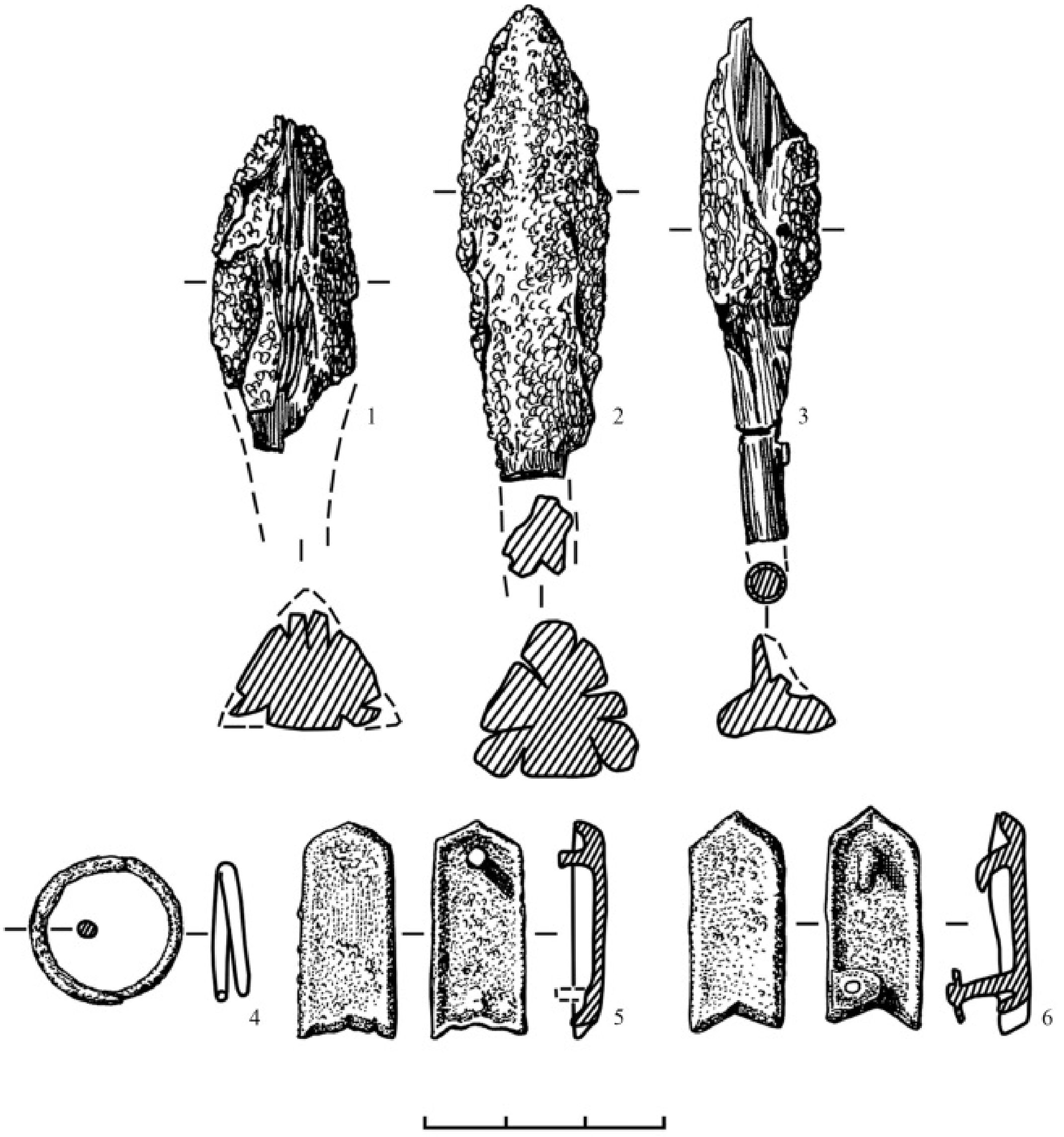


**Fig. S1.24:** Burial at Krasnaya Gorka: 1-6 - finds (from: (*149*)).

**Chineta-II**

The Chineta-II kurgan burial site (*32*, *146*, *150*) is located in the Krasnoshchekovsky District of Altai Krai (Russia), approximately 1 km south-southeast of the village of Chineta (51.185020 N, 83.23903 E), on the second overflow left-bank terrace of the Inya River, a tributary of the Charysh. In the early 2000s, an archaeological expedition of Altai State University carried out excavations at the site. Five kurgans containing both single burials and inhumations with a horse were excavated. Notable finds include iron arrowheads, a knife, horse bits with cheekpieces, stirrups, a bone overlay of a bow and a shaped piece, a bronze garniture from a belt and bridle, earrings, bells, a framed six-petalled pendant, an anthropomorphic pendant in the form of a male's face, and a needle holder in the form of paired fish (Fig. S1.25). The investigated kurgans, including Kurgan 9, are attributed to the Gryaznovo stage of the Srostki culture and dated to the second half of the 9th – first half of the 10th centuries.

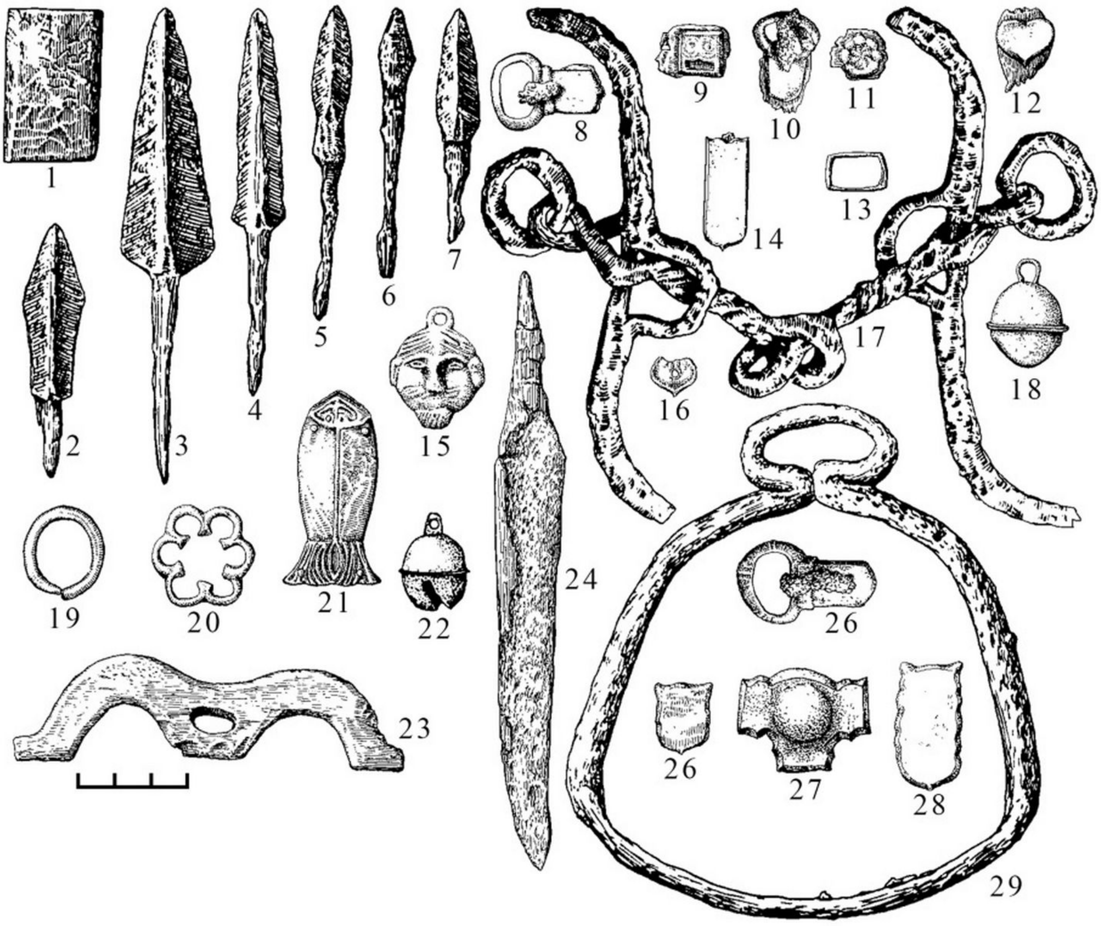

**Fig. S1.25:** Chineta-II kurgan burial site: 1-29 – finds (from: (*32*), Fig. 57: 1–29).

#### Belyi Kamen`

The Belyi Kamen` kurgan burial site (*32*, *151*) is located approximately 1.8 km south of the village of VI Congress (VI Конгресс), Rubtsovsky District, Altai Territory (Russia), about 200 m west of the nearest mountain, Chernyi Kamen`, near a reservoir (51.171931 N, 81.312921 E). All of the kurgans consist of stone-and-earth mounds, sodded and overgrown with low steppe vegetation. The mounds were circular in plan, measuring 4-11 m in diameter and 0.2-0.6 m high. In the centre of each mound, a hollow was recorded, representing traces of looting activities. In 1990, the archaeological expedition from Altai State University investigated sites No. 1, 2 and 3. They found iron arrowheads and spearheads, items of horse equipment, and numerous ornaments of bridles made of non-ferrous metal, as well as tools (Fig. S1.26). The site is attributable to the Gryaznovo stage of the Srostki culture and is dated to the second half of the 9th – first half of the 10th centuries CE.

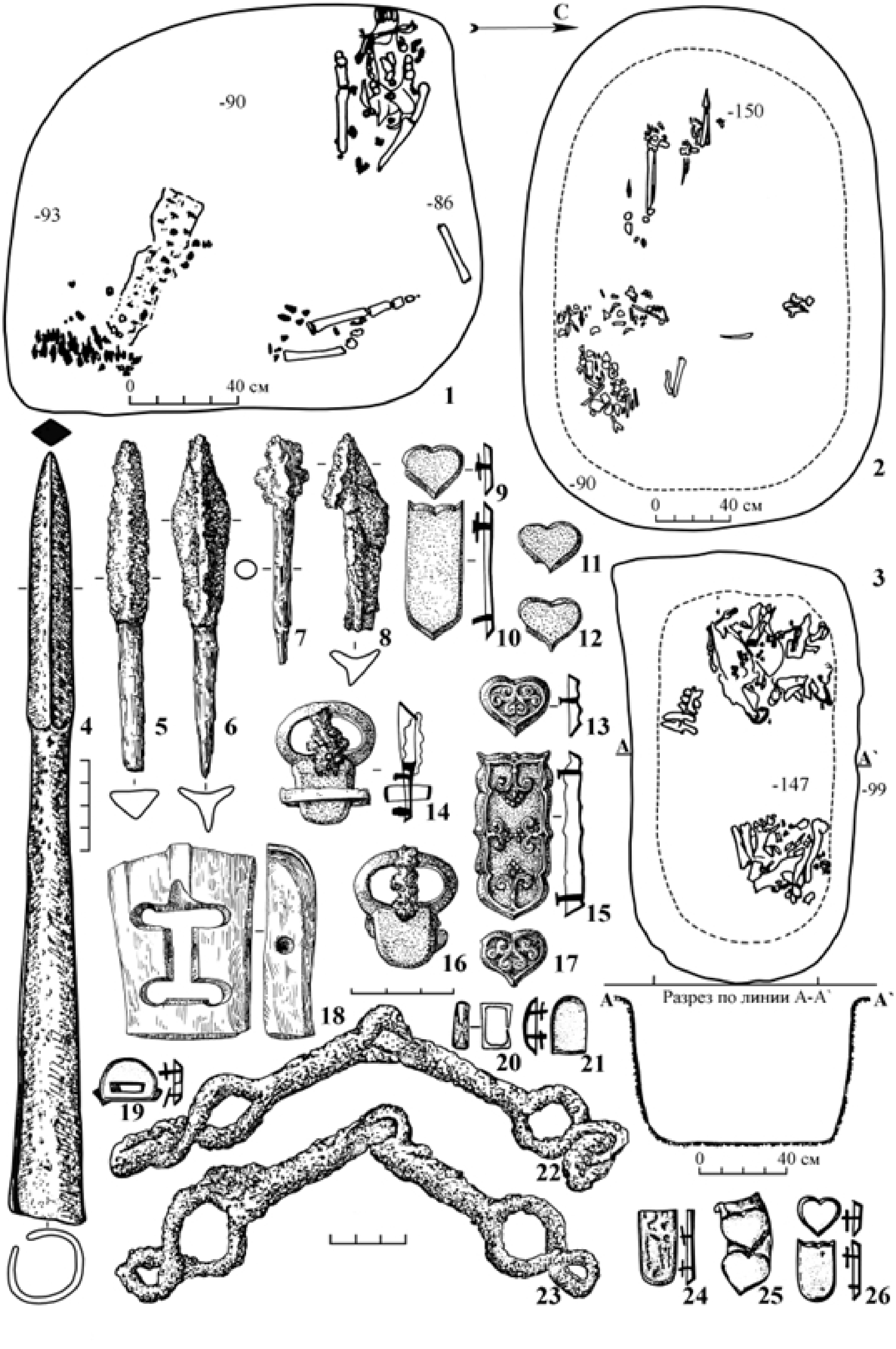


**Fig. S1.26:** Belyi Kamen` kurgan burial site: 1-3 - grave plans, 4-26 – finds (from: Gorbunov, Tishkin, 2025. Fig. 4). The arrow labeled "C" (*Север*, "North") indicates the direction of the north.

### Myshinyi Log-I

The Myshinyi Log-I site (*152*) is located approximately 1 km east of the village of Elbanka, Ust-Pristansky District, Altai Territory (Russia). It is located on the left bank of the Charysh River at the confluence of the Myshiny Log tributary (52.143568 N, 83.373417 E). During excavations in 2011–2012 the expedition of Altai State University investigated two burials. In one of the graves the buried female was lying in lateral decubitus in a contracted position with her head to the east. Osteological materials including a horse skull were found near the south-eastern wall of the pit. Earrings and a ring of non-ferrous metal, iron stirrups, a horse bit with cheekpieces, a plaque, a buckle, and a knife, as well as a clay spindle-whorl and other finds were found in connection with the deceased. The second grave contained a double bi-ritual burial, with one inhumation and one cremation, with numerous accompanying implements including animal bones. The present grave goods, in spite of the object complex characteristic of the Gryaznovo stage (second half of the 9th – first half of the 10th c. CE) of the Srostki culture, differ from the previously known burials by their distinctive funerary rites.

### Gran' solitary kurgan

The single kurgan Gran' (*31*, *32*, *153*, *154*) was located on the border of the Aleisky and Topchikhinsky districts of Altai Krai (Russia), approximately 4 km northeast of the village of Bezgolosovo, on the right bank of the Alei River, in the Ob Plateau (52.331109 N, 83.25287 E). It was excavated in 2000 by Altai State University. The mound was an earthen embankment with a surrounding ditch and a wooden post, measuring 27x30 m in width and 0.7 m high, with a grave in the center. A pit with a fallen wooden post was recorded to the northeast of the grave (Fig. S1.27). The burial construction consisted of a superstructure and an earth-cut grave pit, though the grave was targeted by looters. In the southwestern part of the grave bones from two horses were found, likely piled there by the robbers. Initially, these horses were most likely located on the covering above the burial chamber. Damaged and whole bones from the skeleton of a male aged 20–25 and from another horse were found in the grave pit among the remains of a wooden burial chamber (Fig. S1.27). Judging by the position of the cervical and caudal vertebrae, this horse was laid on the bottom of the grave along its southeastern wall, to the left of the deceased, and was oriented in the same manner as the individual, with its head to the northeast. Grave goods include bronze gilded plaques and tips from bridles and belts, a bronze belt buckle, iron arrowheads and spearheads, a girth buckle, fragments of a sword, a horse bit with sidebars, a leather bridle strap, and whetstones (Fig. S1.27). The mound is considered to belong to an individual from the 'elite' of the Srostki period, and dated to the second half of the 10th – first half of the 11th century CE (*153*).

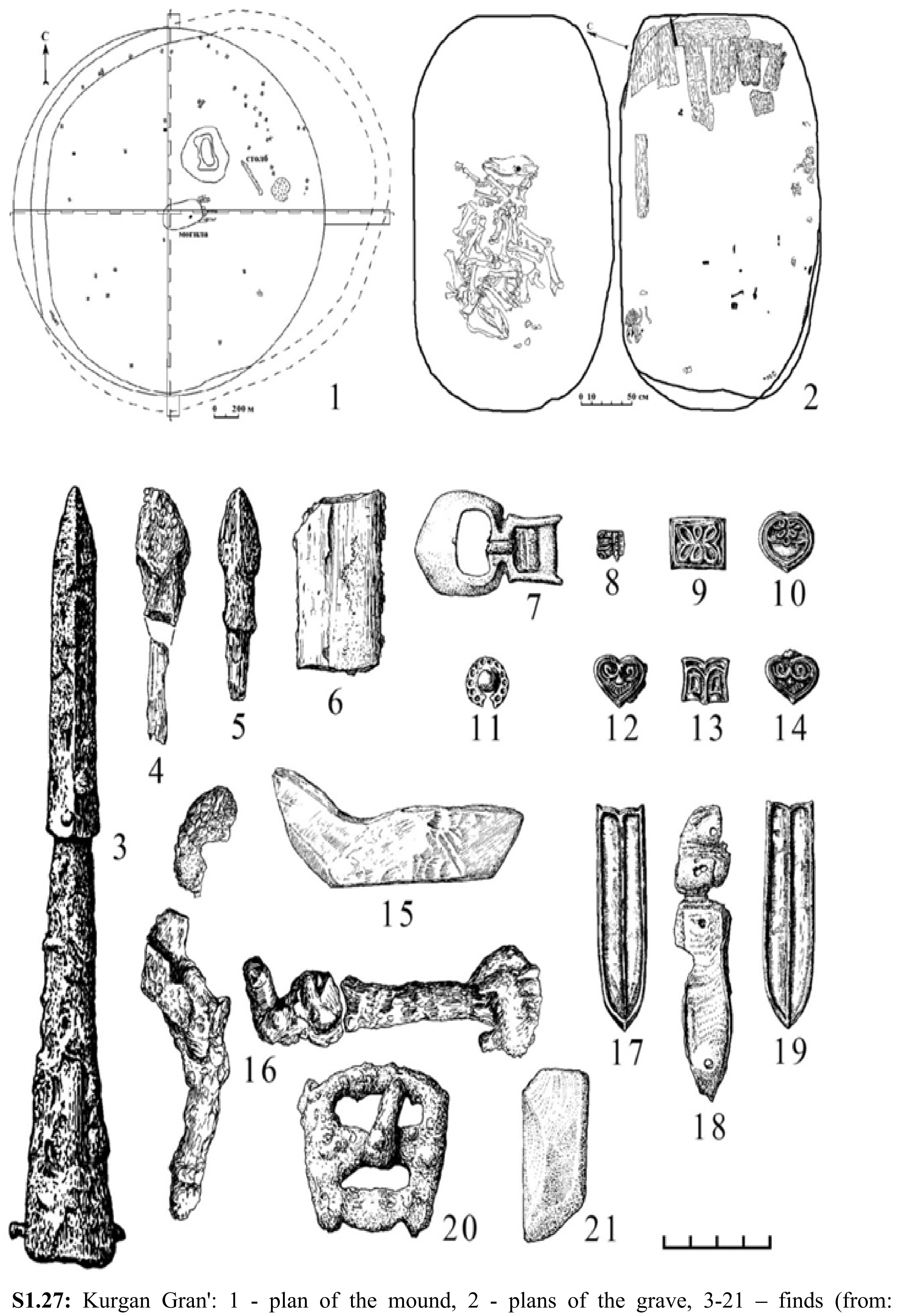


**Fig. S1.27:** Kurgan Gran': 1 - plan of the mound, 2 - plans of the grave, 3-21 – finds (from: Gorbunov, Tishkin, 2025. Fig. 20). The arrow labeled “C” (*Север*, “North”) indicates the direction of the north.

**Prudskoi**

The Prudskoi kurgan burial site (*31*, *32*, *155*) is located in the Kalmansky District, Altai Krai (Russia), approximately 4 km southeast of the village of Prudskoi and 5.5 km northwest of the village of Shadrino, on a promontory of an interfluve terrace of the left bank of the Shadriha River (left tributary of the Ob), on the Ob Plateau (53.082010 N, 83.320827 E). The site consists of six earthen kurgans ranging from 10-16 m in diameter, and 0.2-0.35 m high, which were excavated by Altai State University in 2001 (Fig. S1.28). One mound was surrounded by a ditch, three featured wooden frame enclosures, and three had wooden posts (Fig. S1.28). Within the mounds, 10 single inhumation burials were discovered (Fig. S1.28). Among the material finds in the graves, it is worth mentioning iron arrowheads, swords, knives, raspers, bridles, plaques, horse head ornament, strap distributors and strap-ends, bridle strap-ends, saddler's awls, a bronze buckle, earrings, pendants, bone buckles and bow grip overlay with remains of the wooden arc of the bow and birch-bark binding, and a wooden arrowhead (Fig. S1.28). The site is dated to the second half of the 10th – first half of the 11th century CE.

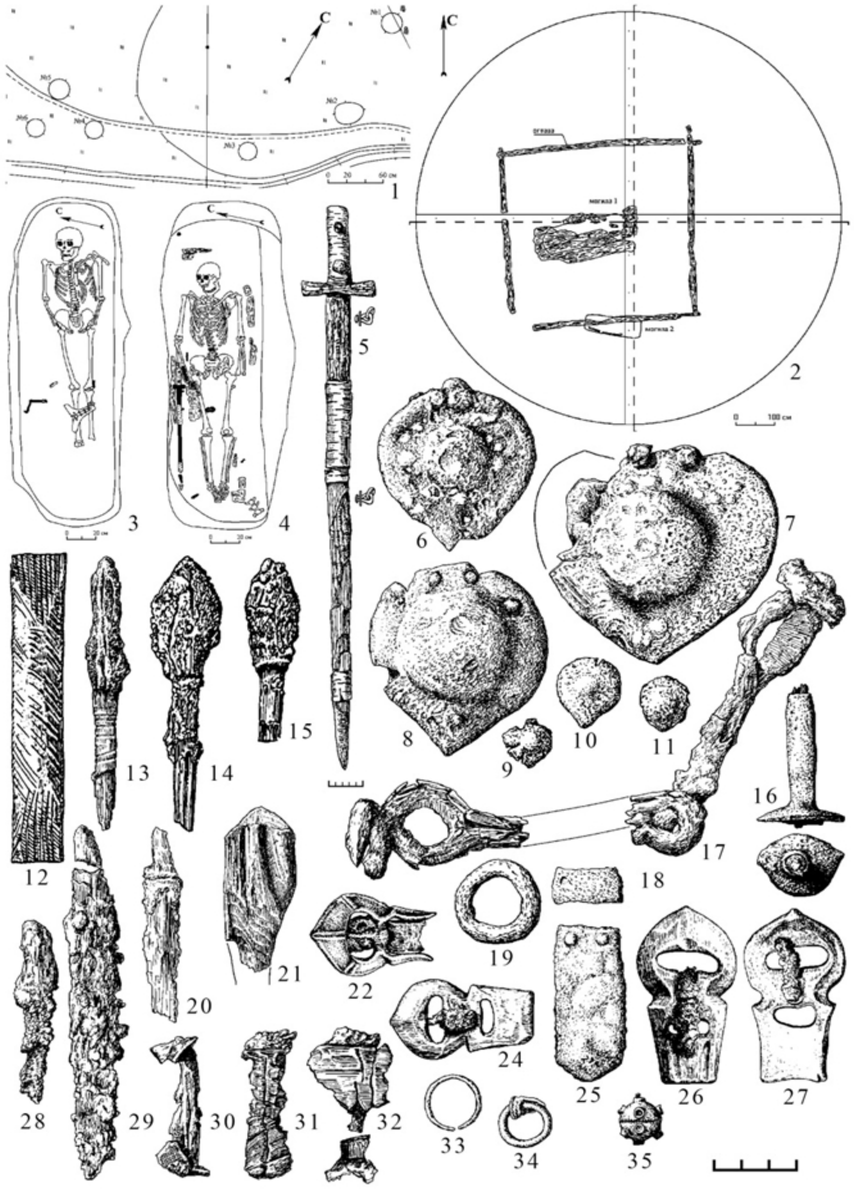

**Fig. S1.28:** Prudskoi burial mound: 1 - plan of the site, 2 - plan of the burial mound, 3-4 - plans of graves, 5-35 - finds (from: Gorbunov, Tishkin, 2025. Fig. 47). The arrows labeled “C” (*Север*, “North”) indicate the direction of the north.

**Srostki-I**

The Srostki-I kurgan burial site (*32*, *156–160*) is located in the Biysk District of Altai Krai (Russia), on the eastern outskirts of the village of Srostki (southern slope of Mount Piket), approximately 25 km southeast-east of the city of Biysk, on the right bank of the Katun River, within the foothill zone of the Biysk-Katun interfluve (52.241991 N, 85.425118 E). This site allowed the identification of the Srostki archaeological culture of the early Medieval period in the south of Western Siberia. The site, which has been the subject of several excavation campaigns, consists of more than 60 earthen mounds ranging from 4-14.5 m in diameter and 0.3-1 m high (Fig. S1.29). In 2012-2014 and 2016 the research was carried out by the archaeological expedition of Altai State University. A total of 56 kurgans were excavated at the necropolis, but only six of them were fully opened (Fig. S1.29). Materials from 66 graves were studied. The burial rites are represented by single and paired inhumation, inhumation with a horse, cremation, and burial of a horse leather (Fig. S1.29). Important material from the site includes iron swords, arrowheads and spearheads, horse bits with sidebars, stirrups, adzes, fire-strikers, knives, files, buckles, clips, ring hooks, antler bow plates, cheekpieces, birch-bark quivers, bronze buckles, plaques and strap-ends from belts and bridles, two-piece fasteners, earrings, pendants, Chinese coins, and glass and stone beads (Fig. S1.29). Of particular note are bronze, partly gilded parts from the scabbards and hilt of swords with vegetal motifs and images of lions, and anthropomorphic pendants and plaques (Fig. S1.29). All finds are dated to the second half of the 9th – first half of the 10th century CE.

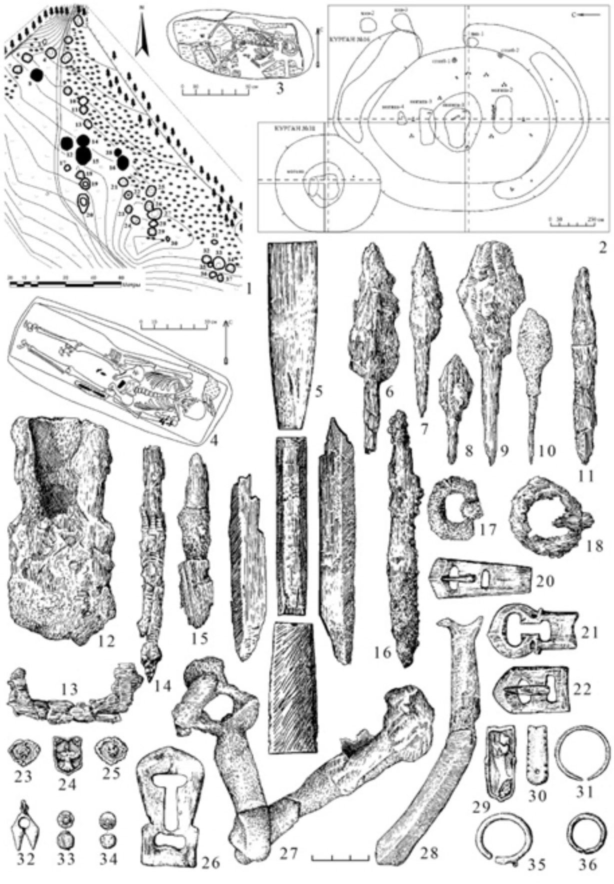

**Fig. S1.29:** Srostki-I kurgan burial complex: 1 - plan of the site in 1930, 2 - plan of the burial mound, 3-4 - plans of graves, 5-33 – finds (from: Gorbunov, Tishkin, 2025. Fig. 50). The arrow labeled "C" (*Север*, "North") indicates the direction of the north.

#### Yarovskoe-III

The Yarovskoe-III kurgan burial site (*31*, *161*) is located in Aleisky District, Altai Territory (Russia), approximately 2.8 km northeast of the village of Bezgolosovo and 0.5 km east of Lake Yarovskoe, on the right bank of the Alei River, on the Ob Plateau (52.331887 N, 83.2530 E). In 1997, Altai State University excavated Kurgan No. 1, which is an earthen mound 20 m in diameter, and 0.35 m high. The kurgan had been previously looted, but nevertheless contained a double inhumation within a wooden burial chamber (Fig. S1.30). Beads made of various materials (rock crystal, carnelian, and chalcedony), a fragment of an iron sword blade, a part of a bronze mirror (Fig. S1.30), and fragments of patterned fabric were found, dating the burial from the second half of the 9th to the first half of the 10th century CE.

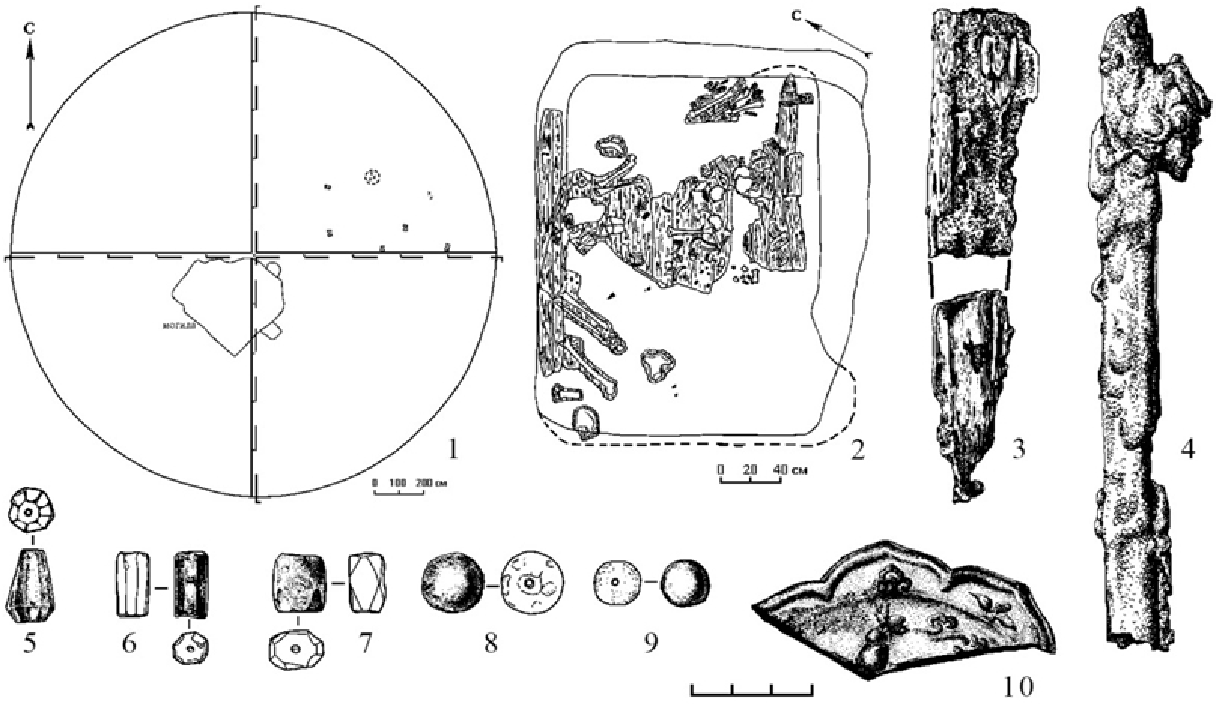


**Fig. S1.30:** Yarovskoe-III kurgan burial site: 1 - barrow plan, 2 - grave plan, 3-10 – finds (from: Gorbunov, Tishkin, 2025. Fig. 62: 1–10). The arrows labeled "C" (*Север*, "North") indicate the direction of the north.

#### Yarovskoe-V

The Yarovskoe-V kurgan cemetery (*31*, *154*) is located in Aleisky District, Altai Territory (Russia), 2 km southwest of the village of Bezgolosovo and 0.9 km south of Lake Yarovskoe, on the right bank of the Alei River, on the Ob Plateau (52.32093 N, 83.12200 E). It was investigated in 2000 by Altai State University. Four earthen kurgans ranging from 8-12 m in diameter and 0.2-0.35 m high were excavated (Fig. S1.31) constructed with wooden posts and containing a total of seven single burials.

**Kurgan No. 1** is the northernmost in the chain. Three graves were found beneath the mound. The skeleton of a female aged 40–45 lying supine with her head to the northeast was recorded at the bottom of Grave 1 (the central one). Traces of a birch bark bedding were present, along with the remains of a coarse cloth, which covered the deceased. An iron knife, the bones of a sheep, and the remains of a birch bark bowl with grain husks were also found in connection to this grave. The skeleton of a subadult laid supine with its head to the northeast was uncovered at the bottom of Grave-2. The skeleton of a young child, approximately three years old, lying supine and oriented with the head to the northeast, was found in Grave-3. The studied mounds are dated to the second half of the 10th – first half of the 11th century CE.

**Kurgan No. 2** was located 8 metres south of Kurgan No. 1. Grave-1, the central grave, contained the remains of an adult female and a child of 6-7 years old were found in it. This mound showed traces of looting activities. In grave–2, the skeleton of a male aged 30-35 was found at the bottom of the pit, lying supine with his head to the northeast. The remains of fire-strikers were found with him.

**Kurgan No. 3** was located 6 metres southeast of Kurgan No.2. The disturbed skeleton of a male aged 50-55 was found at the bottom of Grave 1, laid supine in an extended position, with the head to the northeast. Iron horse bits with ring-shaped cheekpieces, an iron knife, flint, sheep bones, etc. were found.

**Kurgan No. 4** was located 14 metres south of Kurgan No. 3. Grave-1 had been previously disturbed, and contained the remains of the skeleton of a male aged 45–50 years laid supine with his head to the northeast. Two iron tips, a fragment of an iron knife, a stone disc with a perforation, an iron awl, the sacrum and vertebrae of a sheep were found. Along the long walls of the grave pit were the remains of a double frame made of logs and planks.

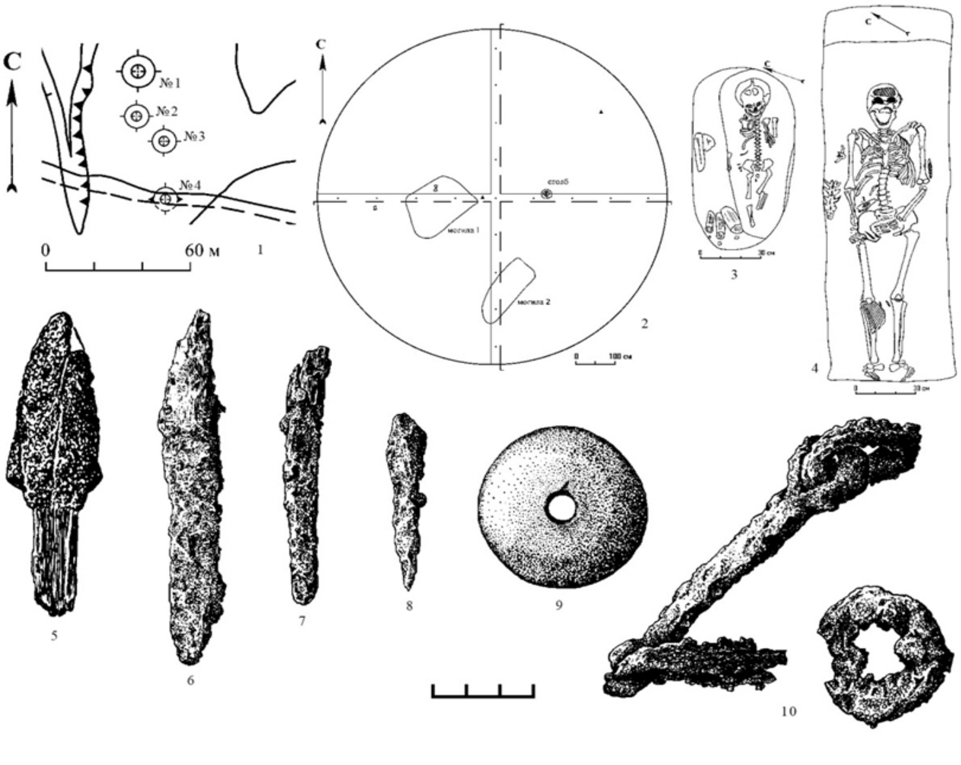

**Fig. S1.31:** Kurgan cemetery Yarovskoe-V: 1 - plan of the site; 2 - plan of the kurgans; 3-4 - plans of graves, 5- 10 - finds (from: Gorbunov, Tishkin, 2025. Fig. 2: 11–20). The arrows labeled "C" (*Север*, "North") indicate the direction of the north.

**Uspenovka-II**

The Uspenovka-II kurgan cemetery (*31*, *162–164*) is located in the Aleisky district of Altai Krai (Russia), approximately 1.75 km northeast of the village of Uspenovka and 3.5 km southwest of the village of Novokolpakovo, on the bedrock terrace of the left bank of the Alei River, Ob Plateau (52.321055 N, 82.545911 E). The site consisted of three earthen mounds, which were 17–23 m in diameter and 0.7–1.7 m high (Fig. S1.32). In 2002, Altai State University excavated a large kurgan, **No. 3,** with a ditch and a posthole, which contained three single inhumations (Fig. S1.32). Various archaeological materials were found within the graves and mound, including: an iron sword blade, a file, an awl, an ornamented horn case, a ceramic vessel (Fig. S1.32). According to archaeological data, Kurgan No. 3 is dated to the second half of the 9th – first half of the 10th centuries CE. The AMS-dating (Ua-49364) of the canine bones is 1100±33. Calibration data (by 1δ (sigma) (68.2%) 895–925 CE, 935–985 CE; by 2δ (sigma) (95.4%) 880–1020 CE) confirmed the indicated range, and also allow its refinement – the end of the 9th – the first quarter of the 10th centuries CE.

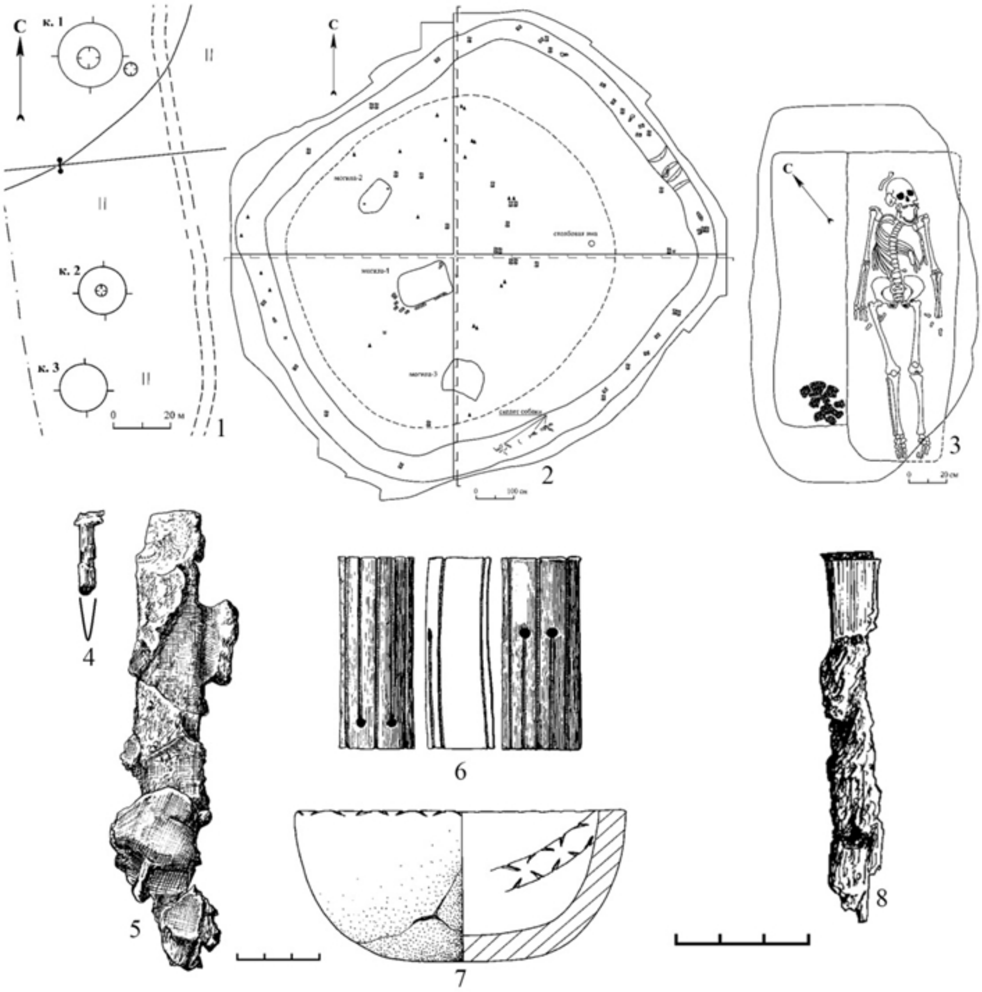


**Fig. S1.32:** Kurgan cemetery Uspenovka-II: 1 - plan of the site, 2 - plan of the barrow, 3 - plan of the grave, 4-8 – finds (from: Gorbunov, Tishkin, 2025. Fig. 54: 1–8). The arrows labeled "C" (*Север*, "North") indicate the direction of the north.

### Popovskaya Dacha

The Popovskaya Dacha single kurgan (*31*, *32*, *153*, *155*) was located in Aleisky District, Altai Territory (Russia), approximately 3.25 km northeast of the village of Bezgolosovo and 0.6 km north-northeast of Lake Yarovskoe, on a promontory on the right bank of the Alei River, on the Ob Plateau (52.334392 N, 83.13002 E). In 2001, it was excavated by the Altai State University. The earthen mound, constructed with a ditch and wooden posts, had dimensions of 25x30 m and a height of 0.8 m. It contained five graves (Fig. S1.33), in which a collective inhumation with two horses, a single burial and a double burial were recorded. The finds include a piece of decorated silk textile, a fragment of a metal mirror, hoop earrings, bronze clasps, belt mounds and strap-ends, as well as iron swords, arrowheads, horse iron bits with sidebars, stirrups, a horse's forehead ornament, horse head ornament tube/plume, strap distributors, and knives (Fig. S1.33). The mound is considered to be 'elite', dated to the second half of the 10th - first half of the 11th century CE and belongs to the Srostki culture (*153*).

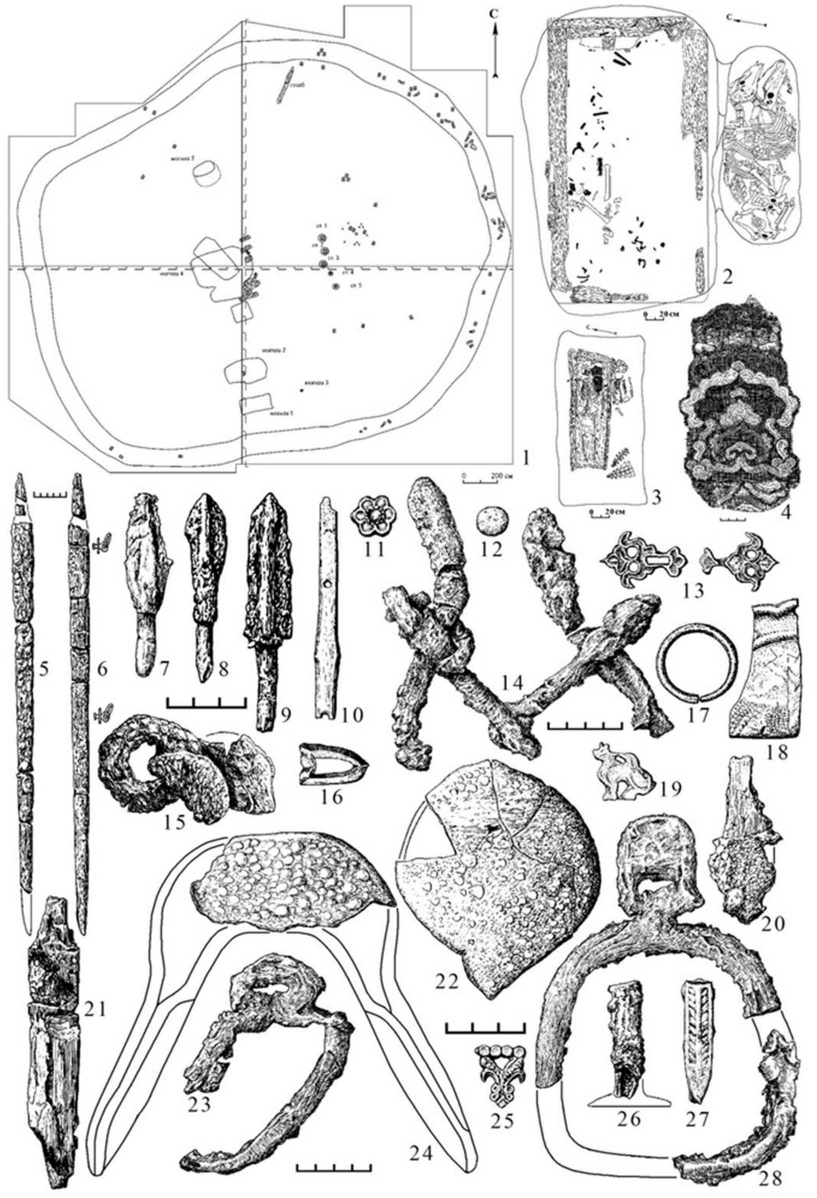


**Fig. S1.33:** Popovskaya Dacha barrow: 1 - plan of the barrow, 2-3 - plans of graves, 4-28 - finds (from: Gorbunov, Tishkin, 2025. Fig. 7). The arrows labeled "C" (*Север*, "North") indicate the direction of the north.

**Blizhnie Elbany-IX (Bol’shie Elbany-IX)**

The Blizhnie Elbany-IX kurgan burial site (*32*, *165*, *166*) is located in the area ‘Blizhnie Elbany’ near the village of Chauzovo, Topchikhinsky District, Altai Krai (Russia), near the Bolshaya Rechka River, on the right bank of the Ob River (52.44815 N, 83.44717 E). On the surface of the sand dune, eight mounds ranging from 7–12 m in diameter, and 0.3–0.4 m high, were initially clearly visible, with traces of earlier excavations. In 1946, 1947 and 1949, the expedition of the State Hermitage and the Institute of the History of Material Culture of the USSR Academy of Sciences under the leadership of M.P. Gryaznov excavated Blizhnie Elbany, including the site in question. Kurgan No. 1, which had a wooden frame and pillars and contained an inhumation burial with a horse hide, was examined. Many items were found, including iron spearheads and arrowheads, a knife, a buckle, a spoon, horn bow fittings and arrowheads, and a silver pendant on a chain (Fig. 1.34). In 1994, a joint expedition of Altai State University and the Museum of History, Literature, Art and Culture of Altai investigated Kurgan No. 2, beneath which there were seven burials. Some of the graves, including number 7, belong to the Zmeyevo stage of the Srostki culture and are dated within the second half of the 11th – 12th centuries CE. Various finds were discovered: a fragment of a metal mirror, an anthropomorphic buckle, paired clasps made of non-ferrous metal, beads, and other items (Fig. 1.34).

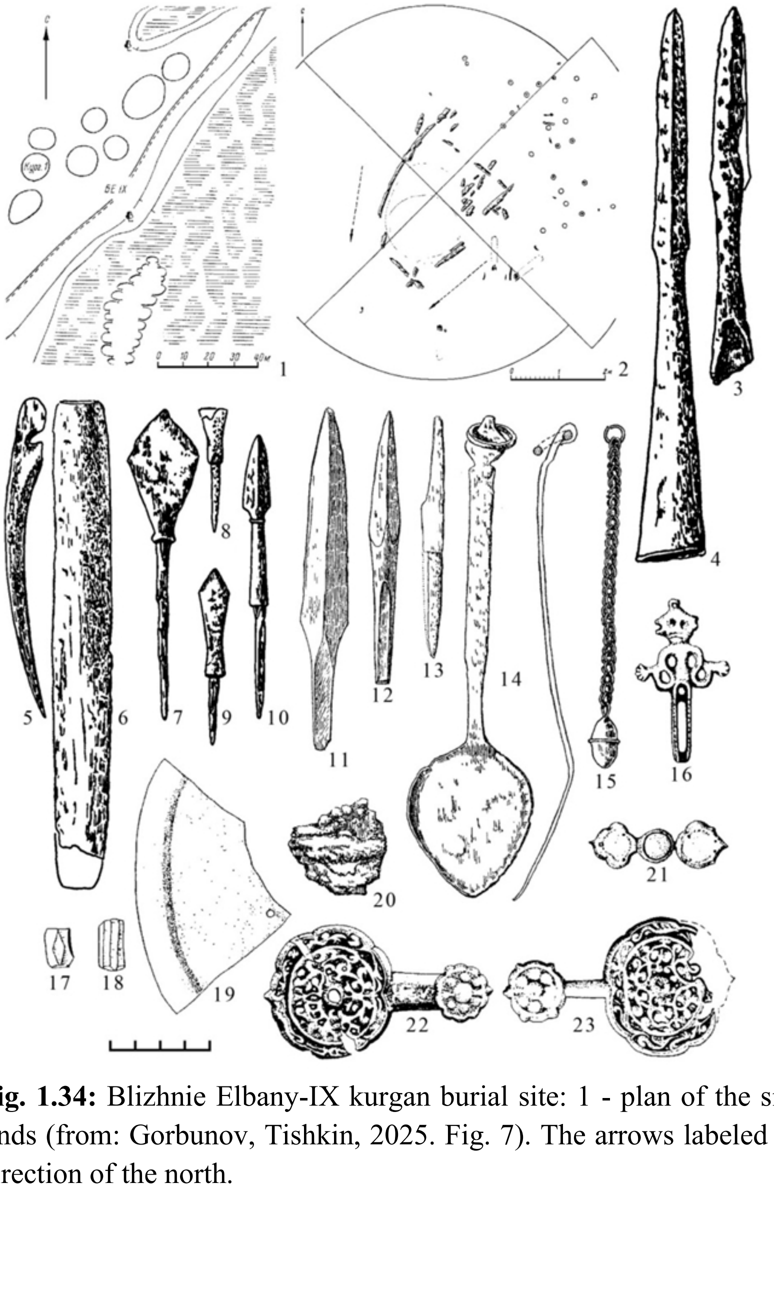

**Fig. 1.34:** Blizhnie Elbany-IX kurgan burial site: 1 - plan of the site, 2 - plan of the kurgan, 3-23 – finds (from: Gorbunov, Tishkin, 2025. Fig. 7). The arrows labeled “C” (*Север*, “North”) indicate the direction of the north.

**Karmatsky**

The Karmatsky archaeological complex (*167*) is located on the right bank of the Ob River, opposite the city of Barnaul, near the settlement of the same name in Pervomaisky District, Altai Krai (Russia). The Mongol-period kurgans are situated on a promontory about 20 meters high, up to 120 meters wide, and more than 100 meters long (53.265310 N, 83.48513 E). The promontory is bordered by an oxbow lake known as Komarinoye Lake, and the territory of the complex is covered with pine forest. In 2002, Altai State University completed the excavation of Kurgan No. 8, which, prior to the excavations, was a rounded earthen mound measuring approximately 5 m in diameter and 0.15 m in height. In the centre of the mound, rounded depression containing a rectangular pit of modern origin was recorded.

Grave-1 was oval in plan, measuring 2.5 m by 0.86 m, and had been disturbed by looting. The fill contained fragmentary remains of an adult skeleton, as well as traces of an internal grave structure. In the western part of the grave, at the bottom, a cluster of ribs, clavicles and vertebrae was recovered. In the eastern part, an in situ fragment of an inner grave structure was preserved, within which the inferior limbs (legs) of a male individual were found. The deceased was lying supine, with his head to the west, in a burial chamber constructed from a wooden frame covered with birch bark. Grave- 2, containing a child, was recorded in the northwestern part of the excavation, outside the barrow mound. Grave-1 from Kurgan No. 8 is dated to the 14th century CE and is attributed to the Karmatsky culture.

A sculptural reconstruction was made based on the skull of a male excavated from Kurgan No. 9 of the same burial site, the nearest to Kurgan No. 8. The reconstruction was made to reflect the physical attributes of one of the individuals of the site, and perhaps represent the typical features of one the population of the Karmatsky culture (Fig. 1.35).

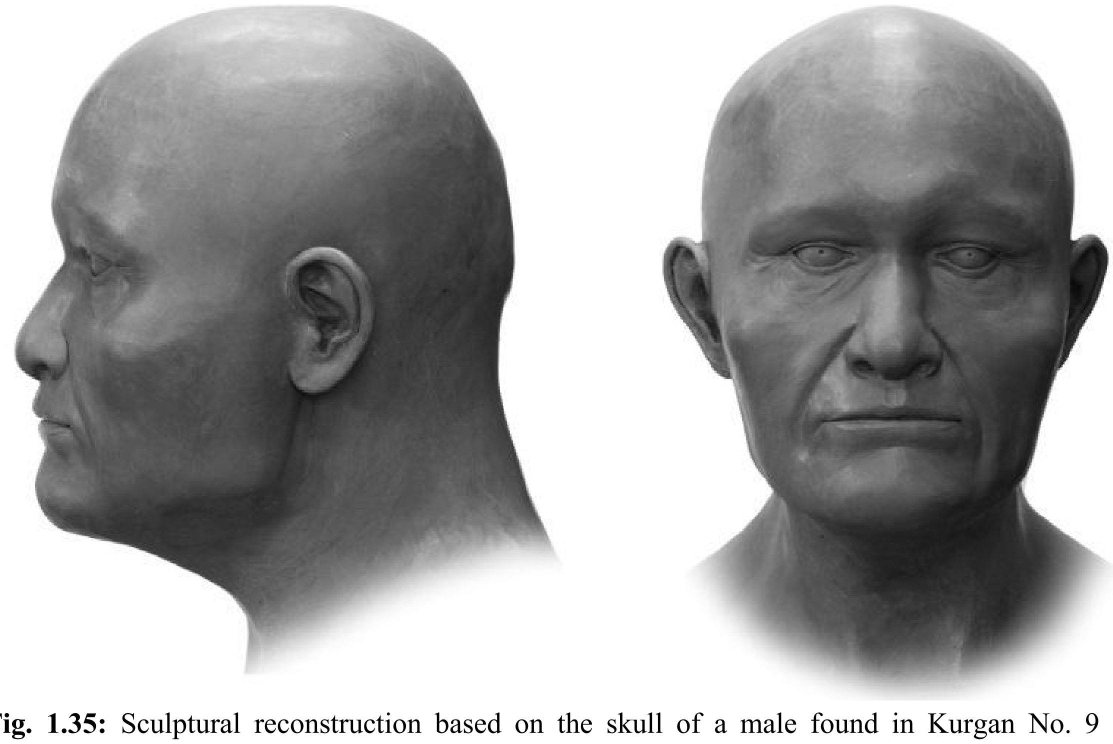

**Fig. 1.35:** Sculptural reconstruction based on the skull of a male found in Kurgan No. 9 of the Karmatsky burial complex (reconstruction by D.V. Pozdnyakov).

**Supplementary Text S2: Previously published Altaian groups involved in the study**

Three previously published Mountainous Altai individuals (RISE600, RISE601, RISE602) were classified in the original article (*49*) and the v54.1 AADR dataset as Iron Age without accurate dating and archaeological affiliation to any material culture. We discovered that they corresponded to the Bulan-Koby period in the region. The burial of RISE602 (Sary-Bel burial site) was an early monument of the Bulan-Koby culture dated to 2nd century BCE – 1st century CE (*168*), and we labeled this individual as Altai_Mt_BulanKoby_Early (Table S2). On the other hand, RISE600 and RISE601 were excavated from the Verh-Uymon burial site, and were actually dated to the second half of the 4th – 5th centuries CE in the archaeological studies (*18*). We present them as Altai_Mt_BulanKoby_published (Table S2). All three individuals exhibit a similar genetic profile to Altai_Mt_BulanKoby, and were kept in the dataset, but were not combined into a single group except for the IBD analysis, as the published individuals' samples were not UDG-treated unlike ours (*49*).

Another individual from the same study (*49*), RISE504 from the Kytmanovo burial site in Forest-Steppe Altai was radiocarbon dated to 709-888 calCE. However, this burial site was investigated in the study for the Andronovo culture context, which the other individuals sampled for the study were assigned to. We could not find the correct archaeological context for RISE504 and labeled him as Altai_FS_Medieval_published (Table S2).

The published individuals from the Eastern Kazakh Altai spanned a long time frame between the Iron Age and Medieval periods, all of whom except two were from (*2*) since those from (*169*) were not yet available during the preparation of the study (Table S2). For the separation of the Pazyryk period individuals from the Berel_50BCE group, where they were assigned to in the original 2021 study, we considered another report by the same team which described the $^{14}C$ dated BRE001 and BRE002 as associated with the Pazyryk culture (*170*). BRE001 is labeled as Berel_300BCE, being dated to 360-175 calBCE. Two other published individuals (I0562 and I0563) (*48*), together with BRE002 (354-182 calBCE) are labeled Berel_300BCE_Eastern to indicate their elevated East Asian ancestries (Fig. 2; Table S2). On the other hand, Berel_300CE group from the Eurasian Migration Period is taken the same as the original study (*2*), and the Medieval period Karakaba_830CE group from the original study (*2*) is labeled as Kazakhstan_Karakaba.

**Supplementary Text S3: Outlier detection in the newly analysed dataset**

We detected the genetic outlier individuals of a given chronological and regional/archaeological group through Mahalanobis distance and the chi-square distribution tests, on the principal components PC1-PC2 and PC1-PC3 (see Methods). The results are presented in Supplementary Table S3, and the outlier individuals are discussed here. In this and the following sections, we evaluate our Y-chromosomal haplogroup data with the public view Family Tree DNA Discover database (*74*) and ISOGG v15.73 Haplogroup Tree (2019-2020).

**ALT004** (Altai_FS_Odintsovo_oW, "oW" standing for "West Eurasian ancestry outlier") is genetically female (Fig. 2; Table S1). She has elevated Southwest Asian ancestry (Iran_N + Turkey_N) compared to other individuals associated with the Odintsovo culture (Fig. 2). This individual's autosomal composition can be modeled with Altai_Mt_BulanKoby (28.0±8.3%) + SlabGrave1 (17.7±3.8%) + Uzbekistan_IA (54.3±5.2%) in qpAdm when we use the regional model (Fig. 3; Table S6). The burial of the individual (Gorny-10, grave 20) is not different in materials or customs from the rest of the burials at the Gorny-10 burial site (Text S1b).

**ALT049** (Altai_FS_Odintsovo_oE, oE standing for "East Asian ancestry outlier") is genetically male with Y-chromosomal haplogroup Q-YP812/YP829/YP791 (Q1a2a1a4a~ in ISOGG v15.73), which was previously discovered in the Early Medieval (Avar period) Carpathian Basin and also in the Berel kurgans in East Kazakhstan (BRE004 and BRE011 of the Berel_300CE analysis group) (*171*) (Fig. 2; Table S1). He has a predominantly East Asian profile on the PCA and ADMIXTURE, and his ancestries can be represented with Xiongnu_Late as a single source or with Altai_Mt_BulanKoby (24.7±2.7%) + SlabGrave1 (75.3±2.7%) in the regional model (Fig. 2, 3; Table S6). Anthropological metrics of the individual and the excavated burial inventory have strong parallels with those typical of some groups in the Tian Shan region (Text S1b). Considering the radiocarbon dating of this individual (266-531 calCE, 95.4% CI) (Table S1), this profile's appearance in the Forest-Steppe Altai could be related to the post-Xiongnu peoples' westwards expansions from East Asia in the 4th century CE, since other contemporaneous individuals from Inner and Central Asia –including the mentioned Tian Shan area– were discovered to have similar, predominantly East Asian ancestries (DA27, 265-539 calCE; DA127, 214-528 calCE; KRY001, 364-423 calCE) (*1*, *2*). However, in identity-by-descent haplotype sharing (IBD) analysis, while there are 14.16 cM links between ALT049 and DA127, the former has stronger connections to some individuals from the Berel kurgans that do not have elevated East Asian ancestries (43.38 cM total and one 26.61 cM long segment sharing with BRE013, and 38.41 cM total sharing with BRE005) as well as some Avar period individuals in the Carpathian Basin (26.44 cM total sharing with ARK-20 and 20.13 cM with SZOD1-76) (Table S10, 11).

**ALT076** (Altai_Mt_Turkic_Khaganate_oE, oE standing for "East Asian ancestry outlier") is a genetically male individual with Y-chromosomal haplogroup Q-BZ427 (Q1b1a3a1b~ in ISOGG v15.73), whose ancestral lineage Q-L332 was found mainly in IA Central Steppe-related groups (*172*) (Fig. 2; Table S1). His autosomal ancestry can not be modelled with the regional model, however other working models are presented in Supplementary Table S6. The burial of this individual (Taldura-I, mound 2) has a remarkable difference from other Turkic period burials of the Mountainous Altai, where the horse was buried above the deceased and not next to him (Text S1c).

**ALT134** (Altai_FS_Srostki_oN, "oN" standing for "northern ancestry outlier") is genetically male with Y-chromosomal haplogroup N-F4155 (N-Tat), which is not present in any other individual of the newly-analysed dataset (Fig. 2; Table S1). Some of the SNPs (such as N-CTS10336, not defined in ISOGG v15.73 deeper than N-F4155) form further sublineages in FTDNA Discover (*173*). In PCA and ADMIXTURE, this individual exhibits a distinctly northern Siberian profile (though not WSHG-Botai related), similar to the individuals of Nizhneobskaya culture (700-900 CE) whose burials are located in the Khanty-Mansi Autonomous Okrug of Russia (*37*) (Fig. 2. S2-S4; Table S2). No plausible qpAdm models could be established for the individual, neither with the Nizhneobskaya (Ob_Nizhneobskaya) group nor the genetically similar MidIrtysh_UstIshim group; however, his only IBD connection is 13.31 cM sharing with individual I19095 of the Nizhneobskaya group (Table S2, S6, S10, S11).

**ALT158 and ALT160** (Altai_FS_Srostki_oW, "oW" indicating "West Eurasian ancestry outlier") are a genetically confirmed father-and-son pair, buried in separate mounds at the same burial site (Fig. 2; Table S1, S9). They have the Y-chromosomal haplogroup J-PF5041 (J2a2a~), and increased Southwest Asian affinities compared to the rest of the Srostki period individuals (Fig. 2). Using the same regional model with other groups (see Main Text and Methods), ALT158 could be modeled with Altai_Mt_BulanKoby (87.9±4.3%) + Uzbekistan_IA (12.1±4.3%), and ALT160 could be modeled with Altai_Mt_BulanKoby (43.6±9.1%) + SlabGrave1 (25.2±4.2%) + Uzbekistan_IA (31.2±5.6%) (Table S6).

**Supplementary Text S4: Outgroup-$f_3$-statistics and $f_4$-statistics**

Through outgroup-$f_3$-statistics and $f_4$-statistics (*42*), we have compared the successive groups within the respective geographical regions in order to define genetic tendencies and events. We performed outgroup-$f_3$ in the form of $f_3$(x, Test; Mbuti.DG) to catch the shared genetic drift between Test (new analysis groups) and x (previously published modern Siberian groups), where Mbuti.DG (a modern sub-Saharan African group) was the outgroup (Fig. S5; Table S4). With $f_4/D$ analyses, we described the allele sharing patterns of the Target and Test groups. We ran two types of $f_4/D$ analyses, first in the form of $f_4/D$(Mbuti.DG, deep ancestral group, Test1, Test2), second in the form of $f_4/D$(Mbuti.DG, Target, deep ancestral group 1, deep ancestral group 2) (Fig. S6; Table S5). In the first form, we compared the newly analysed groups (here Test) for their excess of allele sharing with the previously-published distal ancient groups. In the second form, we compared between the distal groups representing similar deep ancestries (such as two Ancient North Eurasian-related groups), to define which one would have an excess of allele sharing with the newly analysed groups (here Target). The presented Z-scores for this section are of the $f_4$ results, and the whole list of $f_4/D$ results are in Supplementary Tables S5a and S5b.

We observe heterogeneity between the Pazyryk-period Altai groups in the $f_4$ analyses, where they show varying affinities to the Ancient Northeast Asian (ANA)-related groups (Ulaanzuukh, Baikal_EN, Baikal_EBA, Khövsgöl_LBA, CentralYakutia_LN, LenaRiver_MiddleN). Notably, in the Pazyryk-period Altai, Berel_300BCE_Eastern is the group that shares the most alleles with the ANA-related groups in the form of $f_4$(Mbuti.DG, ANA-related, Test1, Test2), where the |Z-scores| were consistently ≥3. Among the ANA-related sources tested in the form of $f_4$(Mbuti.DG, Berel_300BCE_Eastern, ANA-related1, ANA-related2), Ulaanzuukh and CentralYakutia_LN are candidate sources for the increased East Asian ancestries of Berel_300BCE_Eastern. Taken together with the PCA distributions of the Berel_300BCE_Eastern individuals, Ulaanzuukh stands as a possible candidate to represent a major portion of this group's East Asian ancestry, instead of CentralYakutia_LN (Fig. S3). We comment more on the ANA-related groups below.

In the Mountainous Altai, Bulan-Koby and Berel individuals are shifted towards more eastern ancestries when compared to Altai_IronAge on the PCA and ADMIXTURE, however they roughly overlap with Berel_300BCE (Fig. 2, S2). Moreover, $f_4$-statistics in the form of $f_4$(Mbuti.DG, ANA-related, Altai_IronAge, Bulan-Koby/Berel) shows that the allele sharing of the Ancient Northeast Asian (ANA)-related groups are significantly higher (|Z-scores| ranging 3.423 – 21.538) with the Bulan-Koby/Berel-associated groups than with Altai_IronAge, except for Altai_Mt_BulanKoby_Early (Fig. S6.2, Table S5). Remarkably, in the form of $f_4$(Mbuti.DG, ANA-related, Berel_300BCE, Bulan-Koby/Berel), Berel_300BCE has some signals for elevated East Asian ancestries compared to the later Bulan-Koby/Berel groups unlike Altai_IronAge. The relations of Berel_300BCE with these groups are described further with qpAdm results in the Main Text.

Nevertheless, the 2nd-4th c. CE Berel group (Berel_300CE, different from Berel_300BCE) has significantly higher East Asian ancestry than the other Bulan-Koby/ Berel-associated groups of the post-Pazyryk era (|Z-scores| ranging 9.868 – 21.773). $f_4$-statistics in the form of $f_4$(Mbuti.DG, Berel_300CE, ANA-related1, ANA-related2) show that Berel_300CE shares higher alleles with the Ulaanzuukh group compared to every other ANA-related group (|Z-scores| ranging 4.643 – 14.198), except for CentralYakutia_LN (|Z-score|= 1.298). This suggests that the detected East Asian gene flow could be associated with the Eastern Steppe area.

PCA and ADMIXTURE analyses reveal genetic changes in the newly-analysed dataset, where the autosomal ancestries become more East Asian during the 5th-12th c. CE in both Mountainous Altai and Forest-Steppe Altai regions (Fig. 2, S2-S4). We test these findings with outgroup-$f_3$ and $f_4$-statistics, where we discover signals for stronger genetic continuity in the early stages of transition in both geographical regions (Altai_Mt_Turkic_Early and Altai_FS_Srostki_1), that are followed by more outstanding shifts in later periods (Altai_Mt_Turkic_Khaganate and Altai_FS_Srostki_2/3). In $f_4$ analyses, Altai_Mt_Turkic_Early (single individual) has similar allele sharing tendencies to Altai_Mt_BulanKoby in the form of $f_4$(Mbuti.DG, Altai_Mt_Turkic_Early/Altai_Mt_BulanKoby, ancient group1, ancient group2) (Table S5b). However, Altai_Mt_Turkic_Early is an outlier in the PCA distributions on the significance margin (p= 0.0497) when grouped with Altai_Mt_BulanKoby (Table S3b2). Moreover, outgroup-$f_3$-statistics suggest closer relations between Altai_Mt_Turkic_Early and the groups in the Russian Far East, than we detect for Altai_Mt_BulanKoby (Fig. S5). Most notably, Altai_Mt_Turkic_Early has a significant excess of allele sharing with the ANA-related groups Ulaanzuukh, Baikal_EN, Khövsgöl_LBA and China_LN (|Z-scores| ranging 3.009 – 4.353) when compared against Altai_Mt_BulanKoby in the form of $f_4$(Mbuti.DG, ANA-related, Altai_Mt_BulanKoby, Altai_Mt_Turkic_Early), however not with for Baikal_EBA (|Z-score|= 2.853) and the Yakutian groups (|Z-scores|= 2.248 and 1.365). The Altai_Mt_Turkic_Khaganate group of this period is shifted even more towards the East Asian genetic variety on the PCA and ADMIXTURE (Fig. S2, S2-S4). This group also has signals for shared genetic drift with the Russian Far East populations in outgroup-$f_3$-statistics (Fig. S5; Table S4). In $f_4$-statistics, we discover a significant excess of allele sharing between all ANA-related deep ancestral groups and Altai_Mt_Turkic_Khaganate when compared to Altai_Mt_BulanKoby and Altai_Mt_Turkic_Early with in the form of $f_4$(Mbuti.DG, ANA-related, Altai_Mt_BulanKoby/Altai_Mt_Turkic_Early, Altai_Mt_Turkic_Khaganate), where the |Z-scores| range between 3.566 – 20.774. These are in parallel with predominantly East Asian ancestries in Altai_Mt_Turkic_Khaganate, and show a significant shift in the Mountainous Altai towards these ancestries. We observe with $f_4$-statistics in the form of $f_4$(Mbuti.DG, Altai_Mt_Turkic_Khaganate, ANA-related1, ANA-related2) that three ANA-related groups show highest cladality with this group: Ulaanzuukh, Baikal_EN and CentralYakutia_LN (Fig. S6.2; Table S5b). The relations of Neolithic Yakutian and Cis-Baikalian, and Bronze Age Eastern Steppe groups require further clarification through future analyses; at present, we can not distinguish between these groups at a statistically significant level based on the $f_4$ results. However, previous research showed that the Eneolithic Cis-Baikalian ancestry was later replaced by the Bronze Age Cis-Baikalian ancestry (Baikal_EBA in our study) (*7*, *12*) which our results indicate no connections for the Turkic Khaganate period Mountainous Altaians; and contrarily, Ulaanzuukh-related ancestry persisted into the Xiongnu and Medieval eras through the succeeding Slab Grave culture's population (*3*). Similarly with Berel_300BCE_Eastern, the PCA distributions of the Altai_Mt_Turkic_Khaganate individuals indicated Ulaanzuukh as a possible candidate to represent a major portion of this group's East Asian ancestry, instead of the Neolithic Yakutians (Fig. S3).

In ADMIXTURE, Altai_FS_Odintsovo_1 and Novosibirsk_UpperOb groups demonstrate high amounts of allele sharing with the WSHG_Botai group (~30%) that represents Neolithic-Eneolithic Ancient North Eurasian (ANE)-related peoples of Southern Siberia, in relation to the recently-defined North Eurasian hunter-gatherer cline (*12*). We validate the ADMIXTURE results and outgroup-$f_3$ results with $f_4$-statistics. In outgroup-$f_3$-statistics, Altai_FS_Odintsovo_1 and Novosibirsk_UpperOb have the highest drift with modern Kets and Selkups among modern Siberians, whose ancestries and ANE-related connections are mentioned further with references below and in the Main Text (Fig. S5; Table S4). The ANE-related groups, namely WSHG_Botai, Altai_HG, Tarim_EBA_1, Okunevo_BA and Central_Steppe_EMBA (different from Steppe_EBA, which represents the Yamnaya-associated pastoralists) have the highest allele sharing with Novosibirsk_UpperOb and Altai_FS_Odintsovo_1 in the newly-analysed dataset when tested in the form of $f_4$(Mbuti.DG, ANE-related, Test1, Test2) (Fig. S6.1, Table S5a). Moreover, $f_4$(Mbuti.DG, Target, ANE-related1, ANE-related2) indicate a significant (|Z-scores| ranging between 3.768 – 12.618) excess of allele sharing of Novosibirsk_UpperOb and Altai_FS_Odintsovo_1 (and later Altai_FS_Srostki_1, |Z-scores| ranging between 3.179 – 7.175) with the WSHG_Botai and Tarim_EMBA_1 when compared to the other ANE-related groups (Fig. S6.3, Table S5). Furthermore, $f_4$(Mbuti.DG, Novosibirsk_UpperOb/Altai_FS_Odintsovo_1/ Altai_FS_Srostki_1, WSHG_Botai, Neolithic Yakutians) shows that this certain ANE-related ancestry is likely not related to the Ancient Palaeosiberian components present in the Neolithic Yakutians (represented with LenaRiver_MiddleN and CentralYakutia_LN, evaluated in relation to recent research indicating their genetic impacts on the Siberian population (*12*, *174*)), but still is closely associated with the WSHG_Botai group (|Z-scores| ranging between 3.077 and 10.796) (Fig. S6.3, Table S5b). Notably, Tarim_EBA_1 shows a tendency of an excess of allele sharing with these groups when tested against WSHG_Botai in the form of $f_4$(Mbuti.DG, Target, WSHG_Botai, Tarim_EBA_1), at insignificant levels for Novosibirsk_UpperOb and Altai_FS_Srostki_1 (|Z-scores| 1.675 and 2.062) however significant for Altai_FS_Odintsovo_1 (|Z-score| 3.913). We interpret that this may be due to a yet unsampled ancestral profile which can be more similar to the composition of Tarim_EMBA_1, however not an actual contribution from this group since it was genetically isolated in the Tarim Basin (*11*). Nevertheless, some Iron Age groups in Xinjiang (China_Xinjiang_IA_2_Jierzankale and China_Xinjiang_IA_3_Zaghunluq in our dataset) also have high ANE-related ancestry in our ADMIXTURE analyses, therefore the Tarim_EMBA_1-related ancestry may have persisted in the Xinjiang area into the Iron Age (*4*) (Fig S4). We conclude that the ancestries of the Nobosibirsk_UpperOb, Altai_FS_Odintsovo_1 and Altai_FS_Srostki_1 groups have significant contribution from an ANE-related source most closely associated with WSHG_Botai (and possibly a profile similar to Tarim_EMBA_1, yet different). When considered with the outgroup-$f_3$-statistics, a similar deep lineage could also have contributed to the ancestors of the modern Selkup and Ket groups, who are also described with ANE-related genetic affinities in other research (*8*, *58*, *59*). We acknowledge that these analyses do not rule out an undetectable contribution (undetectable in our analyses) from the other groups that did not have an excess of allele sharing with our groups when compared against WSHG_Botai.

Altai_FS_Odintsovo_2 has higher East Asian ancestry than the rest of the Upper Ob and Odintsovo associated main analysis groups on the PCA and ADMIXTURE (Fig. 2, S2, S3). After we detect shared genetic drift between Altai_FS_Odintsovo_2 and the Russian Far East in outgroup-$f_3$ results, which suggests closer relations than we detect for Nobosibirsk_UpperOb and Altai_FS_Odintsovo_1, we make a comparison in $f_4$-statistics similar to what we do for the other groups with elevated East Asian ancestries. We detect an excess of allele sharing between Altai_FS_Odintsovo_2 and Ulaanzuukh, Baikal_EN, China_LN in $f_4$(Mbuti.DG, ANA-related, Nobosibirsk_UpperOb/Altai_FS_Odintsovo_1, Altai_FS_Odintsovo_2), where the |Z-scores| range

3.204 – 8.192. When these three ANA-related groups are compared in the form of $f_4$(Mbuti.DG, Altai_FS_Odintsovo_2, ANA-related1, ANA-related2), Ulaanzuukh and Baikal_EN share significantly more alleles with Altai_FS_Odintsovo_2 than China_LN (|Z-scores|= 9.247 and 11.202 respectively), and the results favor Baikal_EN over Ulaanzuukh, however this time insignificant (|Z-score|= 1.797). The interpretation of these findings are discussed above.

Altai_FS_Srostki groups, similarly to their regional precursors (Altai_FS_Odintsovo), exhibit varying genetic drift patterns. Altai_FS_Srostki_1 and Altai_FS_Srostki_2, each show similar shared genetic drift patterns with Altai_FS_Odintsovo_1 and Altai_FS_Odintsovo_2 in the outgroup-$f_3$ tests (Fig. S5; Table S4). Meanwhile, the results for Altai_FS_Srostki_3 resemble those for Altai_Mt_Turkic_Khaganate and not the sampled Altai_FS_Odintsovo variety (Fig. S5; Table S4). $f_4$-statistics in the form of $f_4$(Mbuti.DG, ancient group, Altai_FS_Odintsovo, Altai_FS_Srostki) reveal that the comparative signals for the pairs of Altai_FS_Odintsovo_1 – Altai_FS_Srostki_1, and Altai_FS_Odintsovo_2 – Altai_FS_Srostki_2 show differences, indicating at least minor genetic changes in the region. Altai_FS_Odintsovo_1 has higher allele sharing with the ANE-related and BA Steppe-related groups than Altai_FS_Srostki_1 does in the first form $f_4$ (|Z-score| for WSHG_Botai being 3.565), however has no difference in East Asian allele sharing, suggesting that the change could be due to a different gene flow undetectable here. Altai_FS_Odintsovo_2 has more Steppe-related alleles than Altai_FS_Srostki_2 does, whereas the latter has an excess of allele sharing with Ulaanzuukh and China_LN when compared with the former (|Z-scores|= 3.054 and 3.023). $f_4$(Mbuti.DG, Altai_FS_Srostki_2, Ulaanzuukh, China_LN) shows significantly closer relationship between Altai_FS_Srostki_2 and Ulaanzuukh (|Z-score|= 9.701), suggesting a gene flow originating from the Eastern Steppe region. Moreover, Altai_FS_Srostki_3 shows tendencies in $f_4$ matching the Altai_Mt_Turkic_Khaganate group, differing from the rest of the Forest-Steppe Altai dataset as a whole, yet Altai_Mt_Turkic_Khaganate is cladal with the ANA-related groups in the form of $f_4$(Mbuti.DG, ANA-related, Altai_Mt_Turkic_Khaganate, Altai_FS_Srostki_2), indicating a higher East Asian component in Altai_Mt_Turkic_Khaganate (Fig. S6, Table S5a). Findings for a Mongol Empire period individual associated with the Karmatsky culture in Forest-Steppe Altai (1200-1400 CE) are indistinguishable from Altai_FS_Srostki_2 in $f_4$(Mbuti.DG, ancient group, Altai_FS_Srostki_2, Altai_FS_Karmatsky) (Fig. S6; Table S5).

**Supplementary Text S5: Genetic kinship analyses**

We performed kinship analyses with READ (v2) (*45*) and KIN (*46*) softwares (see Methods). We discovered first and second degree kinship connections between several samples (Fig. S8; Table S9). This allowed us to build pedigrees that connected different burials and burial sites, that could not otherwise be connected without genomic data. Pedigrees with notable findings are discussed below and presented in Figure S8, while the whole list of genetic kinship links can be found in Supplementary Table S9.

**Pedigree 1:** Out of eight total Bulan-Koby associated Mountainous Altaians presented in this study, five individuals belong to this pedigree, spanning to three generations that are connected through the paternal line (Fig. S8). All were excavated from the Choburak-I burial site (Table S1; Text S1a). Therefore, this site likely belonged to a group who viewed genetic kinship as a social cohesion factor. The Choburak-I necropolis is a fully-excavated burial site that represents the "Dyalyan" burial rite (*дяланская традиция обрядовой практики*), where the deceased were buried with advanced weaponry, armour, elaborate horse equipments and a horse "at the feet" of the deceased, associated with the local elite stratum of the Mountainous Altai region in the 4th-5th centuries CE (*175*). In

Choburak-I, three individuals in total were buried with weapons, two of whom exhibit peri-mortem injuries caused by sharp objects (*176*). Another male in the site who also has a similar peri-mortem injury, however, was not buried with weapons. This high level of military activity, and the dating of the site was interpreted to indicate that this group participated actively in the wars during the Xianbei-Rouran transition of the Eastern Steppe region (*177*).

**Pedigree 2:** This pedigree consists of some of the wealthiest burials from the Srostki period Forest-Steppe Altai region, connecting three different burial sites in the central part of the area (Fig. S8; Table S1, S9). These burials are associated with the elite contexts within the Srostki archaeological culture (*31*) (Text S1d). All individuals of this pedigree are grouped under Altai_FS_Srostki_2, which is one of the two main Srostki period groups that have substantial regional genetic continuity from the Odintsovo period, and the males have the haplogroup R-Y20747 (sublineage of R-Y20762), suggesting also a paternal continuity from the Odintsovo groups as it is different from the lineages discovered in the Kimak-period individuals (Table S1; Text S7). Three individuals from this pedigree are present in the runs of homozygosity (ROH) dataset with more than 400k SNPs covered on the 1240k dataset. Two individuals out of three (ALT140, ALT165) have ROH signals for inbreeding between 2nd-3rd cousins, which could indicate endogamy practices in the Srostki period elite (Fig. S7). Given the presence of consanguinity signals in the Odintsovo period Forest-Steppe Altai as well, and lack of these signals in contemporaneous Mountainous Altai and the steppe-related groups of the Avar period Carpathian Basin (*53*, *54*), we interpret that this practice is likely continuous from the Odintsovo period traditions in the region (Fig. S7).

### Supplementary Text S6: Identity-by-descent analyses

After imputation and identity by descent (IBD) autosomal haplotype segment sharing analyses, we built IBD networks in Gephi v0.9.2, showing the haplotype segment sharing between individual pairs (see Methods). We used two networks. At the first network, we filtered for at least one 12 cM continuous segment shared (edge) per individual (node) pair. This way, we aimed to minimize the false-positive IBD links as mentioned in the internet vignette of the ancIBD software (https://ancibd.readthedocs.io/en/latest/Intro.html). In the second network, we used a criterion of at least one stretch of continuous 20 cM shared per pair (20 cM IBD dataset). We used the 20 cM dataset to detect stronger and recent biological connections, since a 20 cM stretch would last only five meiosis events on average (*47*). We note that the samples from four recently-published studies (*53*, *54*, *57*, *169*) were not available during the preparation of our IBD dataset. Relevant information to this section are presented in Supplementary Tables S10 and S11.

#### a. *Leiden clustering and network metrics*

For the 12 cM IBD network consisting of 846 ancient genomes, we used the MultiGravity ForceAtlas2 layout algorithm (*178*), and the Leiden clustering algorithm (*96*) (see Methods). We ran Leiden between 0.01-0.1 resolution levels, increasing the resolution by 0.01 in each step. We observed the clustering patterns and changes in each threshold, and used the 0.01 resolution level that had the highest quality score of clustering. The quality score in 0.01 resolution was 0.858, 0.810 in 0.02 resolution, 0.780 in 0.03 resolution and decreased towards 0.663 in 0.1 resolution. At the 0.01 level, the inferred IBD-sharing communities (clusters) consisted of the largest numbers of individuals with the minimum number of total communities, therefore the data harbored by the clusters at this threshold was the most informative. We used this threshold for the first observation of the 12 cM dataset, which gave 28 clusters with at least five individuals, covering 89.85% of the whole dataset, while the ten largest clusters covered 68.18% of the sample set (Fig. S9; Table S10).

The Leiden algorithm detected several contemporaneous inferred-clusters that overlap on some regions and even burial sites (Fig. S10). Among the ten largest clusters, three are related to the Iron Age Eurasian Steppe populations (clusters 2, 5 and 6), two are related to the Medieval Eurasian Steppe populations (clusters 0 and 8), and one is related to the Medieval Circum-Uralic and –Medieval/Hungarian period– Carpathian Basin populations (cluster 1) (Fig. S9, S10). The Medieval and Iron Age clusters are also inferred by the algorithm to be separate, which can be attributed to the long time interval as series of recombinations lead to diminishing of the IBD connections (*47*) (Fig. S9, S10; Table S10, S11). Therefore, this separation likely does not indicate a genetic discontinuity (*179*), but on the contrary, a continuously connected steppe region through the IA and the Medieval. Contemporaneous clusters (such as 2, 5, 6 for the Iron Age and 0, 8 for the Medieval) are not completely separate entities either, but rather statistically defined groups in the meta-clusters of the Eurasian region (Fig. S9, S10; Table S10, S11). The reason for separate clustering of the mentioned contemporaneous groups may lie in the limitations of Leiden, which was designed for separation of nodes, without considering overlapping communities of IBD-sharing individuals. It should also be considered that the available published sample sets (that cover more individuals from some analysis groups and certain burial sites) likely affect the results (Table S10), and further sampling and research is needed to be able to infer the IBD clusters better. Furthermore, the differences in the endogenous DNA preservation of the ancient individuals, and the nondeterministic nature of the imputation process can result in missing the IBD links which would be detected otherwise (Methods). We focused our analyses on cluster 0 and subcluster 0_0_0 due to their relevance to the Altai region (see below). Cluster 0 separated into subclusters with increased resolution thresholds (0.02, 0.03 and onwards), where subcluster 0_0_0 (in 0.03 resolution) remained the largest subcluster of the dataset and of cluster 0.

In order to prevent sample and analytical bias in the following analyses, we kept the statistically-inferred IBD clusters as “different modules”, and centered our analyses on the main IBD cluster (cluster 0) of the newly-analysed individuals (see below). To analyse the data more objectively, we calculated network metrics (Table S10) after filtering out close kinship connections (≥1200 cM): Calculations for degree ($k$, number of total IBD links of a single node), degree centrality (calculated by $k$/n-1 where n is the number of total nodes), clustering coefficient (connectedness of the node’s neighbors to each other), eigenvector centrality (the value of the information stored within the node, high values meaning a highly influential node connected to other influential nodes) were done through Gephi; calculations for number of cliques (a subset consisting only of nodes that are all adjacent to each other, “complete” subgraphs) and the number of nodes participating in cliques were done with CFinder (see Methods). Cliques, in their nature, consist of at least three nodes, forming a complete subset. Large cliques can indicate recent shared ancestries, or a direct common ancestor in case of individuals from a kindred (*53*). The number of nodes participating in cliques and the metrics related to the cliques are the only measures without applying the 1200 cM filtering process, in order to preserve potentially informative connections. We then measured the within-module links ($k_W$, subset of k), where the modules correspond to either inferred clusters or the predefined analysis groups (see below). We used individual and average $k_W/k$ ratios (between 0 and 1) to evaluate the inner IBD links, where numbers closer to 1 suggest a more connected cluster/module. We calculated the IBD fractions of the modules to normalise the values, by dividing the number of detected links by total possible internal links similarly with previous research (*47*), which we present for both within-module links and between-module links (Fig. 4).

The network metrics of clusters 0 (Medieval Eurasian Steppe) and 1 (Medieval Circum-Ural and Hungarian period Carpathian Basin) show that these clusters have denser within cluster connectedness than the other clusters: The average clustering coefficients of these clusters (0.203 and 0.224) are higher than the other eight largest clusters (clusters 2-9 averages between 0.043-0.180 clustering coefficients), as well as and the average degrees (13.625 and 12.677 compared to 2.875-7.110), degree centralities (0.016 and 0.015 compared to 0.003-0.008) and eigenvector centralities (0.23 and 0.17 compared to 0.00-0.03). These reflect the dense sampling of the groups in clusters 0 and 1 in our study and previous studies (*37*, *180*). Average $k_W/k$ values (0.897 and 0.849 compared to 0.682-0.972) are in the same margin with the other clusters. Cluster 0 –being the largest cluster inferred in this study– contains a total of 200 ancient individuals from the Eurasian steppe region and its extensions such as the Altai and Tian Shan Mountains (Fig. S10, S11; Table S10). Hence, we interpret that the information stored in cluster 0 is well-informative and sufficient to evaluate the connections further. Individual metrics can be found in Supplementary Table S10.

An overwhelming majority (53 out of 57 imputed genomes) of the newly-analysed Altaians who could be included in the IBD dataset are grouped under cluster 0 (see Methods; Table S10). Together with these, there are also three previously-published Mountainous Altaians associated with the early and late Bulan-Koby period, and one previously-published Forest-Steppe Altaian from the Early Medieval era (Table S10; Text S2). Furthermore, all 12 individuals of the Berel_50BCE and Berel_300CE groups from adjacent Berel kurgans in Eastern Kazakhstan are also in cluster 0. This sums to 69 individuals from and around the Altai region. Notably, 72 Early Medieval (Avar period) Carpathian Basin individuals sampled from the westernmost end of the Eurasian steppe are also in this cluster, which is most likely due to the influx from the Eurasian Steppe region between the 6th-9th centuries CE, described previously (*53*, *54*, *180*, *181*). As mentioned above, the $k_W/k$ value of cluster 0 is 0.897 on average (median=0.923, min=0 [after 1200 cM filtering], max=1). Therefore, most of the connections of the cluster's nodes are within the cluster. In accordance, the clustering coefficient values (mean=0.203, median=0.190, min=0 [after 1200 cM filtering], max=1), being higher than other clusters, describe an increase in connectedness in cluster 0, since it means the connectedness of a node's neighbours (those connected to the node with an edge) to each other as well. Thus, not only the individuals of this cluster have high internal connections, but also their neighbours have so. This is explanatory of the increased eigenvector centralities compared to other clusters as mentioned before (mean=0.227, median=0.109, min=0 [after 1200 cM filtering], max=1), since more connectedness would mean more valuable information, and is supported by the presence of a high number of cliques in cluster 0. The largest "community" of cliques (k=3, cliques of at least three nodes) in cluster 0 has 591 cliques that involve 177 individuals (88.5% of the cluster's individuals and 20.9% of all individuals in the network) and 1074 IBD links (34.9% of all links in the network). This immense connectedness can be ascribed to the numerical dominance of the Altai and Carpathian Basin genomes within cluster 0, which could result in the presence of certain groups from these regions more than the others in the dataset.

Subcluster 0_0_0, subcluster of cluster 0 overlaps significantly with the newly analysed Altaian individuals. We detect the majority of the newly-analysed Altai individuals in subcluster 0_0_0 regardless of their genetic profile (51 out of 57), and we define this subcluster as the "Altai IBD cluster". It has 85 ancient individuals in total, where the rest of the individuals are from the Berel_50BCE and Berel_300CE (the relations explained below), the rest of Central Asia and the Medieval (Hunnic and Avar period) Carpathian Basin. Notably, the contemporaneous individuals from the region of modern Mongolia are lower in number in this cluster compared to those from modern Kazakhstan (also after excluding the Berel groups of Eastern Kazakhstan), where we see the Kazakhstan Kimak-Kipchak, Karluk-Karakhanid and Karakaba groups' individuals clustered in subcluster 0_0_0 (Table S10). The value of the Altai cluster data and the internal connections of its groups are observable on the IBD metrics: average within-subcluster degree ($k_W$) 16.59, average $k_W/k$ 0.81, average clustering coefficient 0.25, average eigenvector centrality 0.42. The largest "community" of cliques (k=3, cliques of at least three nodes) of cluster 0_0_0 has 394 cliques that involve 84 individuals (42.0% of the cluster 0 individuals and 9.9% of all individuals in the network) and 677 IBD links (54.7% of the links in cluster 0 and 22.0% of all links in the network).

b. ***Analysing the Altai through modules***

Our findings above indicate that the variety in the sampled dataset is representative of the Altai region and their connections in Inner Asia. After these evaluations, we investigate the links of the contextually important analysis groups of the IBD dataset, by using them as modules in a similar manner to the customary analyses' groups to increase comprehensibility (Fig. 4; Table S10). The groups chosen as modules are the newly presented and published ones from the Altai, as well as those from other regions surrounding the Altai in Inner Asia. Please see Methods on how the modules were prepared for this section. The modules of the Mountainous Altai show a tight connectedness to each other, similarly with our findings using the Leiden algorithm (see above), whereas the Forest-Steppe Altai modules have more complicated internal patterns. The links between the groups of the two regions are scarcer than the within-region links they have. The findings are detailed below.

From a group of 21 total individuals of the early and late Bulan-Koby periods (Altai_Mt_BulanKoby_Early and Altai_Mt_BulanKoby, the latter a combined group of newly and previously published individuals associated with the Bulan-Koby culture since adverse effects of the lack of UDG-treatment was not expected here) in the Mountainous Altai and the Berel_50BCE and Berel_300CE groups in the Eastern Kazakh Altai, 17 individuals build an influential subset. These 17 individuals form the twenty largest cliques with 10-11 individuals and 119 IBD links (3.9% of all links in the 12 cM dataset), while we note that these cliques include close relatives. The connections of these analysis groups are presented in Figure 4, with high numbers of between-module IBD fractions for the observed links from total possible links (38.9-87.5%). When considering the qpAdm results, where the Altai_Mt_BulanKoby analysis group can be modeled with Berel_50BCE to represent ~95% of their ancestries, and that both groups can be modeled similarly in the regional model with Berel_300BCE as a single source (Fig. 3; Table S6), this phenomenon implies the presence of a homogeneous population in the pre-Medieval era Mountainous Altai – Eastern Kazakhstan regions, continuous for around 600 years prior to the genetic shifts we describe in the Main Text. Particularly, this population contributes to the Turkic period Mountainous Altaians (represented with Altai_Mt_Turkic_Khaganate since Altai_Mt_Turkic_Early did not pass the imputation quality check) who have similarly high between-module IBD fractions with the early and late Bulan-Koby period groups combined (42.2%, 19 of 45 possible links between) and with Berel_50BCE (33.3%, 10 of 30 total possible links between). We detect decreased links between Altai_Mt_Turkic_Khaganate and Berel_300CE (16.7%, 5 of 30 possible total links).

We see varying rates of IBD-sharing in the Forest-Steppe Altai region. The Odintsovo period modules (Altai_FS_Odintsovo 1 and 2) both have high within-module links (100% and 47.2%) considering the whole dataset, but the connections between the two groups are lower in comparison (15.6%, 7 from 45 total possible links between). Altai_FS_Srostki_1 has higher rates of links with Altai_FS_Odintsovo_1 (60%, 6 out of 10 total possible links) compared to Altai_FS_Odintsovo_2

(16.7%, 3 from 18 total possible links). Contrarily, the Odintsovo period modules have similar rates of IBD-sharing with Altai_FS_Srostki_2, which counted for 31.8% for Altai_FS_Odintsovo_1 (27/85 total possible links) and 26.8% for Altai_FS_Odintsovo_2 (41/153 total possible links). The IBD signals of Altai_FS_Srostki_3 align with their non-local origin we detect in other analyses, as the links are only sporadic (0-2 detected links with the other Forest-Steppe Altai groups corresponding to 0-4% of total possible links). This group instead has more prominent links to the Berel_50BCE and Altai_Mt_BulanKoby (16.7% and 7.5% of total 35 and 40 possible links).

Between-region IBD links of the Altai groups are not as strong as the within-region links, and most of the detected links between the modules of the two Altai geographical areas are mediated by Altai_FS_Odintsovo_2 (57.5%, 88 links with the Mountainous Altai-Berel groups from 153 total between-region links), which has 18.5-55.6% of total possible links with the Mountainous Altaians. The rest of the 67 links between the other groups (modules) represent lower ratios of between-module IBD fractions (0-20% of total possible links between the module pairs), where the highest fractions for detected out of total links are of Altai_FS_Medieval_published that has one detected link with Altai_Mt_Turkic_Khaganate from five total possible (20%) and another sole link with Berel_300CE (16.7%, from six total possible links). The remaining pairs of modules have a maximum of 14.3% detected/total links. Altai_FS_Medieval_published module consists of a single individual previously $^{14}$C dated to 709-888 CE (Text S2), making it penecontemporaneous with Altai_Mt_Turkic_Khaganate in the Mountainous Altai and contemporaneous to the Srostki-period groups, notably without any detected IBD link to the latter.

#### c. ***Strong connections in the 20 cM threshold IBD dataset***

As mentioned above, the 20 cM IBD dataset was used to detect the strong IBD links between the newly-analysed Altaians and also their connections to other groups, where we considered the same modules defined on the 12 cM IBD dataset. The IBD sums presented in parentheses below are "sum_IBD>20" per individual pair, and the total IBD sums in "sum_IBD>12" are higher than the 20 cM sums in their nature (Fig. 4, S11; Table S11).

Having not detected 20 cM IBD links for the Pazyryk-period modules, we then evaluate the post-Pazyryk period findings. Altai_Mt_BulanKoby, Altai_Mt_BulanKoby_Early and the Berel groups (both Berel_50BCE and Berel_300CE) show pronounced links between several individuals except between Altai_Mt_BulanKoby_Early and Berel_300CE (ranging 19.0-62.5% of total possible links, three longest pairs sum 60.5 cM BRE009-ALT071, 57.4 cM BRE007-ALT060, 50.2 cM BRE013-ALT066), verifying strong connections between these groups in Mountainous Altai – Eastern Kazakhstan described above (Table S10, S11). These are likely more distant than 6th degree relationships, given there are less than at least three ≥20 cM stretches per pair (*47*). Both the Bulan-Koby period and the Berel individuals on this network have a long time-range (50 BCE – 450 CE and 150 BCE – 335 CE), indicating long term biological continuity in the Mountainous Altai and its surroundings, repeating the findings we describe in the 12 cM IBD dataset. RISE602, the early Bulan-Koby period individual, has three stretches of ≥20 cM IBD sharing (sum 75.9 cM) to ALT071, a late Bulan-Koby period individual. RISE602 could be an ancestor to ALT071, or could be related to an ancestor of the latter.

Connections of these groups with the Odintsovo period population in the neighbouring Forest-Steppe Altai are varying, and depend on the analysis group. Every strong IBD link from the Bulan-Koby associated modules with the Odintsovo groups is to the individuals of Altai_FS_Odintsovo_2 (15 detected ≥20 cM pairs, 18.5% of total possible and three longest pairs sum 33.5 cM ALT030-ALT056, 29.8 cM ALT011-ALT060, 28.8 cM ALT011-ALT062), lacking any links with the Altai_FS_Odintsovo_1 group (Table S10, S11). Berel_50BCE shows the same pattern of connections (5 detected ≥20 cM links, 9.3% of total possible and three longest pairs sum 48.2 cM BRE010-ALT029, 24.6 cM BRE010-ALT001, 22.3 cM BRE003-ALT029). Compared to these, the detected links between the two main Odintsovo analysis groups 1 and 2 are relatively weaker (a total of two strong links that are 4.5% of total possible, sum 21.2 cM ALT009-ALT036 and 21.0 cM ALT031-ALT042) (Table S10, S11). These results correlate with the qpAdm models that indicate relatively dissociated population histories for the Odintsovo analysis groups. We can not evaluate the Novosibirsk_UpperOb - Altai_FS_Odintsovo IBD links due to the low quality DNA yield of the Novosibirsk_UpperOb samples.

Biological continuity is observable between the Turkic period groups (Altai_Mt_Turkic_Khaganate, Altai_FS_Srostki) and their regional predecessors (Altai_Mt_BulanKoby, Altai_FS_Odintsovo). Altai_Mt_Turkic_Khaganate individuals in the Mountainous Altai have strong links to both the late Bulan-Koby period (Altai_Mt_BulanKoby) group and Berel_50BCE (three detected ≥20 cM links with both, 7.5% of total possible and sum 35.0 cM ALT056-ALT099, 27.9 cM RISE601-ALT092, 21.0 cM RISE601-ALT090 for the former; 10% of total possible and three longest pairs sum 21.6 cM BRE010-ALT092, 20.6 cM BRE003-ALT090, 20.5 cM BRE012-ALT090 for the latter), but there are no strong links detected with Berel_300CE or Altai_Mt_BulanKoby_Early. In the Forest-Steppe Altai, Altai_FS_Srostki_1 has strong links to both Odintsovo period groups (two links with Altai_FS_Odintsovo_1, 20% of total possible and 26.1 ALT043-ALT114, 21.0 cM ALT052-ALT109; whereas one link with Altai_FS_Odintsovo_2, 5.6% of total possible links and 37.7 cM ALT009-ALT109). Altai_FS_Srostki_2 group too, has detected strong links with individuals from both Altai_FS_Odintsovo_1 and Altai_FS_Odintsovo_2, but more prominent with the latter (4.7% of total possible links and three longest sum 27.2 cM ALT043-ALT165, 22.7 cM ALT048-ALT123, 22.0 cM ALT048-ALT139 for Altai_FS_Odintsovo_1; 7.8% of total possible links and three longest sum 109.7 cM ALT011-ALT147, 46.3 cM ALT018-ALT150, 36.9 cM ALT002-ALT150 for Altai_FS_Odintsovo_2) (Table S11, S12). Contrarily, Altai_FS_Srostki_3 does not exhibit any strong IBD links to the Odintsovo period samples, and neither to Altai_FS_Srostki_1 or Altai_FS_Srostki_2.

Notably, Altai_FS_Srostki groups do not exhibit any strong links to Altai_Mt_Turkic_Khaganate either. The ancestries present in Altai_Mt_Turkic_Khaganate and Altai_FS_Srostki_3, despite being similar to each other, may have arrived in the Altai with different groups given the lack of connecting IBD signals. In contrast, two published contemporaneous individuals from North Kazakhstan (Kimak-period DA93, Kipchak-period DA23) have pronounced ≥20 cM IBD links with Altai_FS_Srostki_2 (8 detected pairs where 7 are of DA93, 23.5% of total possible links and longest sums 78.5 cM between ALT118-DA93 with three ≥20 cM stretches, whereas 26.3 cM between ALT118-DA23) and an individual from Altai_FS_Srostki_3 (one link, 10% of total possible and sum 22.2 cM ALT157-DA93) (Table S11, S12). Considering the lack of strong links of DA23 and DA93 to the Odintsovo-period groups and the presence of these links with the Srostki-period groups, our results indicated a gene flow from the North Kazakhstan area that might have had an impact on the formation of the Srostki-period population of the Forest-Steppe Altai region. Therefore, these results suggest a source for the increase of East Asian ancestry and the newcomer groups, yet the scarcity of the samples from North Kazakhstan hinders the possibility of a more detailed evaluation.

**Supplementary Text S7: Y-chromosomal haplogroup data**

We investigated the Y-chromosomal haplogroup data in relation with the rest of our analyses, and found similarities between the observed genetic ancestries and the haplogroups. The Y-chromosomal haplogroups of these populations represent their varying deep ancestral connections. We evaluate our data with the public view Family Tree DNA Discover database (*74*) and ISOGG v15.73 Haplogroup Tree (2019-2020). Time to most recent common ancestor estimates (TMRCA) are taken from the Family Tree DNA Discover database, accessed on 02/09/2025, and may be subject to change with future findings. Supplementary Table S1 contains the new haplogroup data used for the evaluations here, and in Figure S12 we present the results from both the newly-analysed and the published groups.

Despite the genetic heterogeneity in the Iron Age groups of the Altai context (see above and Main Text), the detectable Y-chromosomal gene pool is limited to haplogroups Q-YP771 (Q1b1a3a~) and R-S23592 (R1a1a1b2a2a3~) in four unrelated male individuals of the Iron Age period. While the Berel_300BCE individual is genetically female, and as such does not have a Y-chromosome, the newly analysed 6th-4th c. BCE group's and the Berel_300BCE_Eastern group's males have each haplogroup (or their subclades) (*182*, *183*) (Fig. S12; Table S1). We detect a Y-chromosomal continuity in the region, and the haplogroups Q1b1a and R1a1a1 persist until the 5th c. BCE, whereas not without additional haplogroups' presence after the 1st c. BCE (Fig. S12). The newly analysed 3rd-5th c. CE Bulan-Koby period individuals namely have J-PF5041 (J2a2a~), Q-SK1944 (Q1b1a3b) and R-YP1560 (R1a1a1b2a2a3b1a~) (Table S1); whereas the previously published individuals carry Q-YP844 (sublineage of Q-L715, which is Q1a2a1 in ISOGG v15.73) and J-BY114993 (sublineage of J-PH358 [J2a2a1a1a~]) (*184*, *185*). Notably, some of these lineages correspond to ancestral or detected lineages shared with the Berel groups' individuals (Berel_50BCE, Berel_300CE) who bear Q-YP789 (sublineage of Q-L713 [Q1a2a1a~]) and Q-BZ93 (sublineage of Q-L330 [Q1b1a3]), as well as R-S10885 (R1a1a1b2a2a3c2~) (*171*, *186*, *187*). In a broader perspective, these lineages align with the Iron Age Central Steppe-related autosomal ancestry that consists of major Bronze Age Steppe and East Asian (Baikal_EBA/Khovsgol_LBA) and minor Southwest Asian contributions, as well as the additional East Asian influxes into the region during and after the Xiongnu era (*1*, *2*).

The dominant Y-chromosomal haplogroup in the sampled Turkic period Mountainous Altai individuals is J-M172 (J2), while we also observe Q-BZ427 (Q1b1a3a1b~) and R-YP1560 (R1a1a1b2a2a3b1a~) (Fig. S12; Table S1). The results for individuals who have J2 detectably lead through J2a, J2a2~, J2a2a~, J2a2a1a1~, finally reaching a specific sublineage, J-PH358 (J-PH1795, J2a2a1a1a) in one individual, while we note that the 1x detected SNP is a transition (Table S1). This haplogroup was discovered in several other individuals of Xiongnu-Medieval era Inner Asia including the published early Bulan-Koby period male (*188*). It was also found in a Medieval-Ottoman period male from Anatolia published by Lazaridis et al. in 2022, where it likely arrived with the Central Asian genetic influx after the 11th century CE that was reported in the same study (*189*).

Haplogroups J-M172 and R-YP1560 were discovered in both the Bulan-Koby and Turkic periods Mountainous Altai, which can suggest a paternal continuity, yet more sampling is needed from other regions and also from the Altai, as we report an intense East Asian gene flow in the region after the 5th century CE. The inferred TMRCA of J-PH358 (mean 274 BCE, 758 BCE - 129 CE [95% CI]) (*188*) suggests a recent common patrilineal origin for the carriers of this branch, which could originate from a founder event. In contrast, R-YP1560 has no ancient individuals currently assigned to this haplogroup in the Discover database, and the TMRCA estimate post-dates our Bulan-Koby period samples (mean 817 CE, 536 - 1048 CE [95% CI]) (*190*), therefore either the Bulan-Koby period population may represent the basal lineage for this haplogroup or the actual TMRCA could predate the current estimations. Notably, a specific sublineage of R-YP1560 (R-YP1556 > R-YP1564) is found in the modern Kyrgyz frequently and the TMRCA of YP1564 is inferred to be around 1187 CE (971 - 1363 CE [95% CI]) (*191*). This could indicate connections between the modern Kyrgyz people and the ancient Yenisei Kyrgyz of the Minusinsk Basin near the Altai region.

In the Odintsovo period Forest-Steppe Altai, the paternal lineages are consistent with the analysis groups. The individuals of Odintsovo group 1 who show a high amount of WSHG_Botai-related component have solely haplogroup R-M478 (R1b1a1a1), namely the sublineages of R-L1432 (R1b1a1a1a) and R-Y20762 (R1b1a1a1b), and the latter is shared with an individual of the Novosibirsk Upper Ob group (Table S1). R-L1432 is shared with a male associated with the Eneolithic Botai culture as well as other Neolithic-Bronze Age individuals from Western Siberia (*192*), and its subbranch R-L1433 (R1b1a1a1a, TMRCA 356 BCE - 443 CE [95% CI]), the most recent lineage we can trace to, is shared with DA87 and DA93 from Medieval North Kazakhstan (labeled as Kimak in our dataset), who have different ancestries from Odintsovo group 1 (*193*). R-Y20759 (/R-Y20768 [R1b1a1a1b] and TMRCA 1101 - 33 BCE [95% CI]) is the deepest subclade we can find for R-Y20762 in the Odintsovo period, and is shared with an individual from pre-Medieval North Kazakhstan (*194*).

R-L1433 is present in an individual of the Odintsovo group 2 as well, yet the rest of the haplogroup pool of group 2 is dominated by R-Z93 (R1a1a1b2) and its subbranch R-YP349 (/R-S23592, R1a1a1b2a2a3~ in ISOGG v15.73, TMRCA 2938 - 1589 BCE [95% CI]) where the latter we found also in the Bulan-Koby individuals (see above) (*183*). Yet, according to our observations, the individuals of Odintsovo group 2 constantly possess the ancestral allele for the subclades including that observed in the Bulan-Koby group, and since the TMRCA is distant in time, the individuals of Odintsovo group 2 likely belong to a yet-undefined Y-lineage. Notably, one individual in this group has haplogroup N-FGC28492 (/N-B482 [N2]), a deep ancestral lineage discovered in Eneolithic Baikalians and is traced to in another Botai culture male who has N-FT324, a sublineage of N-B482 undefined in ISOGG v15.73 (*195*).

Y-chromosomal haplogroups of Altai_FS_Srostki_1 and Altai_FS_Srostki_2 show a continuity of the Odintsovo period lineages, with the predominance of the sublineages of R-M478 (R1b1a1a1, discovered in all three unrelated Odintsovo group 1 males) and an occurrence of R-Z93 (R1a1a1b2) (Table S1). However, this pattern should not be interpreted as evidence that the Srostki-period population derived exclusively from the genetic profile of Odintsovo group 1, since unsampled burial sites could uncover additional variability in autosomal component proportions and Y-chromosomal haplogroup composition. Furthermore, we also demonstrate identity-by-descent autosomal haplotype links between both Odintsovo period groups and the Srostki period groups, indicating genetic continuity from a major bulk of the Odintsovo period population regardless of the differences in ancestries (Fig. 4, S11; Text S6). Furthermore, as mentioned above, R-M478 and its subbranches were discovered also in Medieval North Kazakhstan, therefore a more detailed explanation needs more sampling. Nevertheless, individuals from the upper-elite context Srostki burials those analysed in our study, discovered in our analyses to be genetically related, are in genetic group 1, and have the haplogroup R-M478 (Fig. S8; Table S1, S9; Text S1d, S5). The lineages in Altai_FS_Srostki_3 are distinct from the sampled Odintsovo groups and the rest of the Srostki groups, which is in accordance with their non-local ancestries (Fig. S12; Table S1).

**Supplementary Text S8: Freshwater reservoir effect (FRE) analysis of the bone samples**

The actual age of archaeological finds dedicated to radiocarbon dating can easily be ascertained, assuming that its carbon content was in exchange equilibrium with the ambient atmosphere reservoir. In aquatic pools, carbon may however derive from inorganic carbon forms like carbonates, having a significantly lower 14C/12C ratio. Consequently, even a terrestrial sample may inherit an old apparent age if it came into contact with aquatic carbon sources, which phenomenon is called marine or freshwater reservoir effect (FRE). While the marine reservoir effect is well known and widely considered in chronological applications (~400-500 yrs), FRE can significantly vary within lakes or rivers with different characteristics, yielding apparent ages up to 6,000 years (*196*, *197*).

One of the most frequently applied methods for determining reservoir effects is to radiocarbon date aquatic samples of known age. This method uses samples with known collection date results in accurate 14C measurements that can be used to calculate the reservoir effect for that production period. In addition, it can be established by calibrating back the age of a terrestrial sample to yield the expected 14C date of the aquatic sample (*85*, *198*). This method relies on supposing entirely closed contexts where the two types of samples must be contemporaneous and the offset between the 14C dates is expressed in a ΔR value. These values are very important in some regions, where a dietary component from either marine (*199–201*) or freshwater (*202*) systems is present. Due to the scarcity of locations where this effect could be studied, attempts have been made to develop regression and Bayesian models to calculate 14C offsets, and δ13C or δ15N values showed reasonably well correlations with it (*86*, *203*).

Nevertheless, the FRE on radiocarbon dates has been documented from some archaeological locations around the world (*199*, *204*, *205*) and on the Eurasian Steppe, including for example the north Peri-Caspian Sea region and the lower course of the Don River (*206*, *207*), the middle and lower Dnieper basin (*204*) or northeast Kazakhstan (*87*) etc. For example, Schulting et al (2014) presented a successful application of the use of stable isotope data to predict the FRE in humans for the Lake Baikal region. Stable nitrogen values were proved to be more useful over carbon in predicting the extent of the observed 14C offsets. Their linear regression model using δ15N values alone accounted for some 67% of the observed variability in the R, where values around 11-12‰ were associated with an offset of 100-200 years. From other archaeological sites across the Eurasian Steppe, the results of Svyatko et al (2022) overall indicated that the values were highly variable, yet FRE was frequently observed both for faunal and human samples. The maximum R for a human sample was 1071 ± 64 14C years (Tegiszhol, Kazakhstan) and they also managed to indicate a moderate positive linear correlation between the size of R and both δ13C and δ15N for the human samples. From the region of the Altai Mountains, kurgans from the Verkh-Uimon and Kuraika archaeological sites were studied, where the 14C ages of human and animal bones from the same context were compared to calculate the R value. For archaeological samples, a maximum difference of ~400 yrs was revealed but for modern samples, FRE was as high as ~1000 years.

In our FRE analysis five individuals from the Altai Mountains were involved, of which calibrated 14C dates showed significant offsets relative to the expected age ranges. R-modelling was performed using the OxCal v4.4.4 program (*82*, *208*) and the IntCal 2020 calibration curve (*84*). We used rounded dates for the calibrated age ranges during our calculations. Our results are shown in Figures S13.1-S13.5. In the case of ALT037 and ALT042, radiocarbon dates of 1821±17 and 1847±18 were obtained, respectively, corresponding to a calibrated age range around 100-300 CE. The expected range is 500-750 CE, thus a R of around 200-600 years can be assumed based on the modelled offsets

(Figure 1 and 2). The calibrated age range of ALT052 was obtained to be 80-220 CE, while the expected range is 300-500 CE, offering an offset range of approximately 100-350 years (Figure 3). Similar offsets of around 100-250 years can be observed in case of the remaining two individuals ALT134 and ALT168 (Figure 4 and 5). Based on the measured stable isotope results, the 13C value of three human individuals are surprisingly low, below -20‰, while each bone samples with supposed FRE possess elevated δ15N values around or higher than 12‰, indicating the possibility of significant fish or aquatic food consumption.

Nitrogen isotope ratios generally reflect the trophic level of animals and humans, with higher trophic level consumers having relatively higher δ15N values (*209*). As aquatic ecosystems generally consist of longer food chains, elevated δ15N values of aquatic organisms can be used in reconstructing the relative amounts of marine and terrestrial food sources in diets of human populations (*210–212*). The limited isotopic results, having been published in dietary studies, suggest that at least some of the inhabitants of Central Eurasia had a diet rich in animal protein, with freshwater fish perhaps supplying up to 50% of their total protein intake (*213*). Studies of Bronze Age populations also suggest that fish was exploited as a dietary resource in the Bronze Age period across the steppe and forest-steppe zones (*214–216*). From a methodical perspective, fishing is acknowledged as taking place practically everywhere in Central Eurasia, particularly during the earlier periods (*217–219*). Although fish bone is generally absent or likely underrepresented in regional archaeological deposits, fishing implements such as fishhooks, net weights, and harpoons are nevertheless widely attested (*213*).

**Table S1: Metadata and sequencing information of the newly analysed dataset.**
Information about the newly-analysed ancient individuals, included and excluded individuals and samples are both presented.

**Table S2: Published ancient and modern groups used in this study.**
Published groups and individual samples here were used in PCA, ADMIXTURE, outgroup f3-statistics, f4-statistics and qpAdm analyses. The imputed sample dataset used in the IBD analyses were given in Table S10 (IBD annotation table).

**Table S3: Outlier detection in the newly analysed dataset.**
Outlier removal based on Mahalanobis distances and chi-square distributions, on the PC1-PC2 and PC1-PC3 spaces of the PCA. P-value ≤ 0.05 was considered as outlier on each PCA space. We highlighted the outlier samples in red and presented them on gray background.

**Table S4: Outgroup $f_3$-statistics results.**
Outgroup $f_3$-statistics for the newly published groups, who were targeted for their shared drift with the modern Siberian populations. Only the results from the models with at least 100K SNPs were taken into consideration for evaluation. The results are visualised in Figure S5.

**Table S5: f4 and D-statistics results related to the main text.**
$f_4/D$-statistics comparing the newly analysed and published groups for their allele sharing and cladality patterns. Negative Z-scores indicate closer sharing with Test1, whereas positive scores

indicate closer sharing with Test 2. Results with |Z-score| ≥ 3 are deemed significant. The results are visualised in Figures S6.1-S6.3.

**Table S6: qpAdm analyses.**
qpAdm models of the analysis groups in the Altai context. Models with p-value ≥ 0.05 were considered plausible and highlighted in gray. Models with ≥ 20% standard error per any source or standard error higher than the calculated proportion of the source were considered unplausible by the program and therefore were discarded, not being presented here. Chosen models of the groups are visualised in Figure 3.

**Table S7: Admixture times inferred with DATES.**
Admixture times were inferred with DATES in generations and converted into dates by subtracting from the mean group dates assuming 29 years per generation. BCE dates are shown with negative values. Models with standard errors higher than the generation value or Z-score < 2 are shown in red.

**Table S8: Effective population size ($N_e$) estimations.**
Estimations of the groups in the Altai context with hapROH. Outlier individuals were not excluded from the combined groups. Models with negative lower bonds are presented in red. ROH signals used for the estimations can be found in Figure S7.

**Table S9: Genetic relatedness analyses.**
The KIN results were filtered for log likelihood ratios ≥ 1 to prevent false positives. Up to third degree kinship results are presented, and only the highest coverage individual from a pedigree was kept in the main analysis groups for the allele-sharing-based analyses. Pedigrees 1 and 2 are presented in Figure S9 and discussed in Supplementary Text S5.

**Table S10: Annotation table of the identity-by-descent analysis nodes.**
Information about the newly-analysed and published ancient individuals in the identity-by-descent (IBD) haplotype sharing analysis dataset. The computationally-inferred clusters and related IBD metrics are presented here.

**Table S11: IBD-sharing between individuals in the network (edges).**
Identity-by-descent (IBD) haplotype sharing results of the individuals in the IBD dataset. The edges were filtered per node pair to include at least one segment of 12 cM IBD sharing.

**Table S12: Fisher's exact test for IBD comparisons.**
Results of the Fisher's exact test analyses presented in the Main Text. Odds ratio and confidence interval (CI) results are for test order 1 compared to order 2.
