## Supplementary material for "Ancient human genomes from the Altai region reveal population continuity and shifts in the 4th-12th centuries": Figures 1-4 and Supplementary Figures S2-S12

**Figures and legends:**

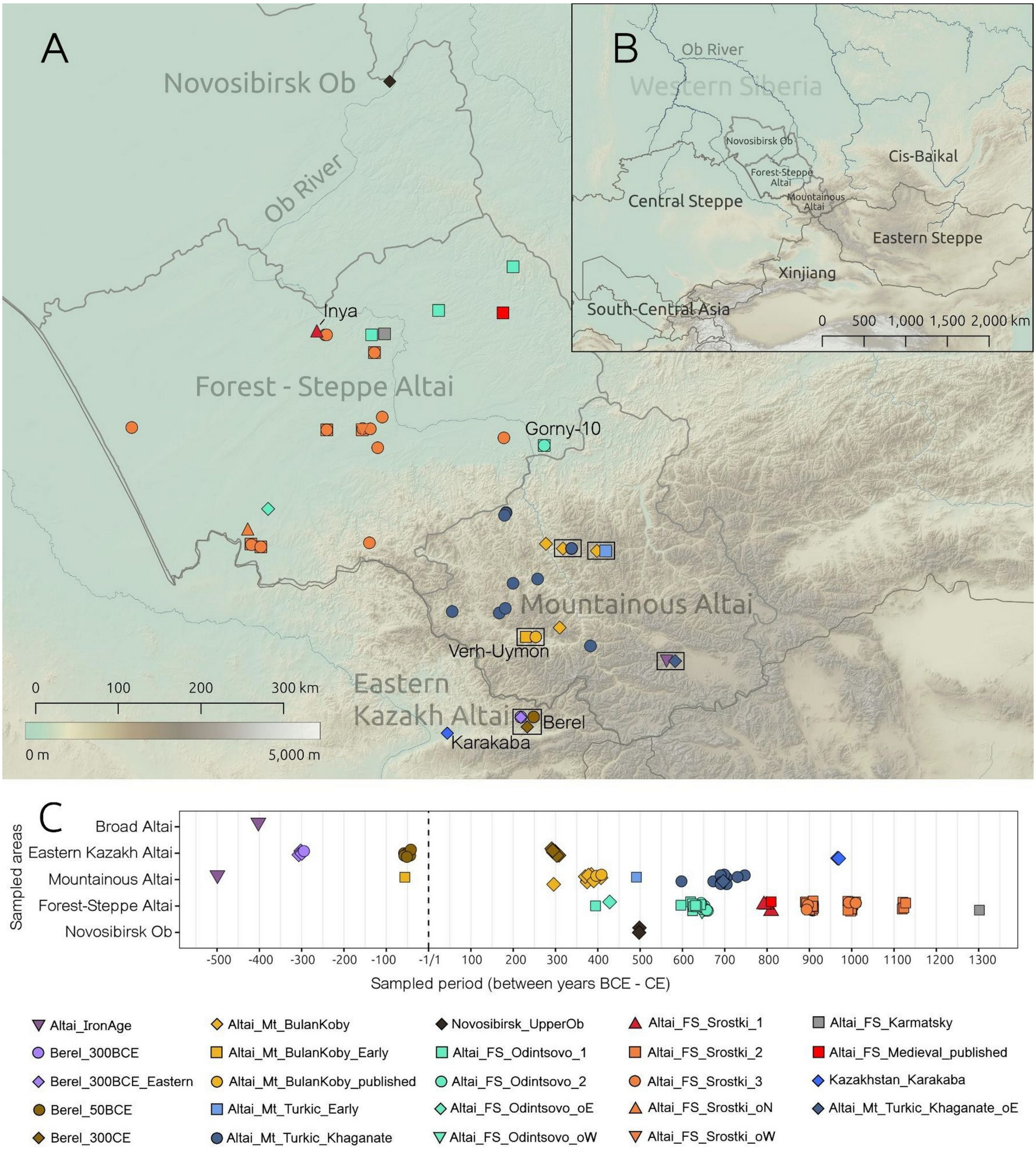


**Figure 1: Sampled burial sites and chronology of the sampled individuals.**
(A) Map of the archaeological sites where newly-reported and previously-published Altaian individuals were excavated. Different periods sampled from the same archaeological sites are jittered and shown in boxes. Sites with multiple genetic profiles from the presented periods have overlapping symbols. The symbols represented on the map are the same as those used in the legend at the bottom of the figure. Borders of internal regions of modern Russia are shown in addition to the country borders. (B) A broad-scale map of the country borders and names, and contextually-important regions. (C) Chronology of the newly reported and published ancient groups from the Altai and Upper Ob Plateau. The symbols and color scheme used are the same as those used in Fig. 2.

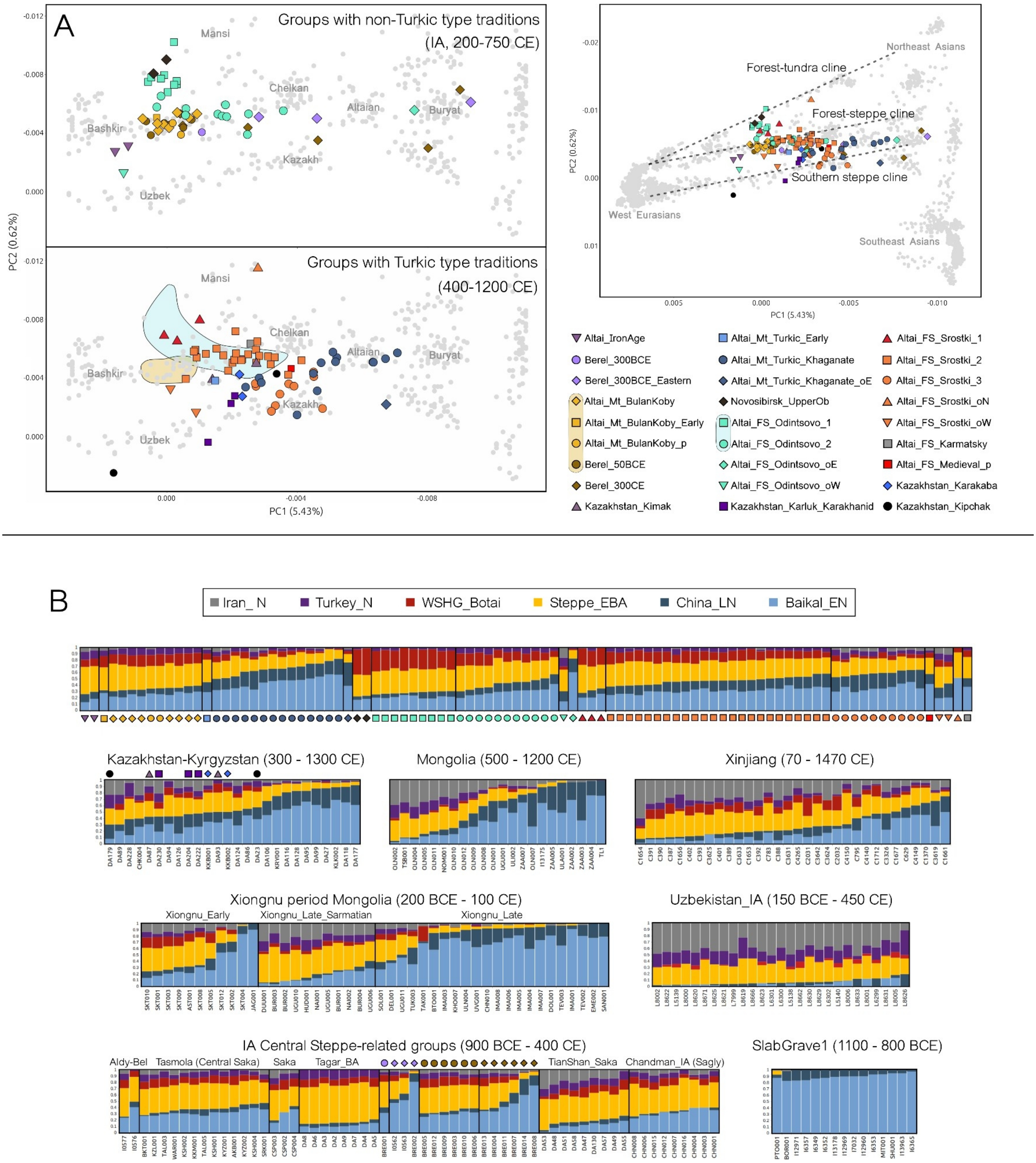


**Figure 2: PCA and ADMIXTURE analysis.**
(A) Principal Component Analysis (PCA) plot of Eurasia on the PC1-PC2 space, based on modern Human Origin dataset (8), including newly reported and published genomes. The right panel shows the complete version of the plot, and the left panels focus on the contextually-important groups from the full Eurasian plot. Ancient genotypes were projected onto the modern genomic variation (gray points). The labels on the PCA plots show broad geographical regions and modern groups. The genetic clines of modern Eurasia are respectively shown in dashed lines on the right panel. (B) K6 supervised ADMIXTURE results of the newly reported genomes, and a subset of published ancient individuals. Inner Asian groups mentioned in the main text are presented here. Individuals per each analysis group are in order according to their increasing total East Asian ancestries (Baikal_EN + China_LN). The symbols in the PCA plot legend also represent the individuals plotted in the upper row of (B).

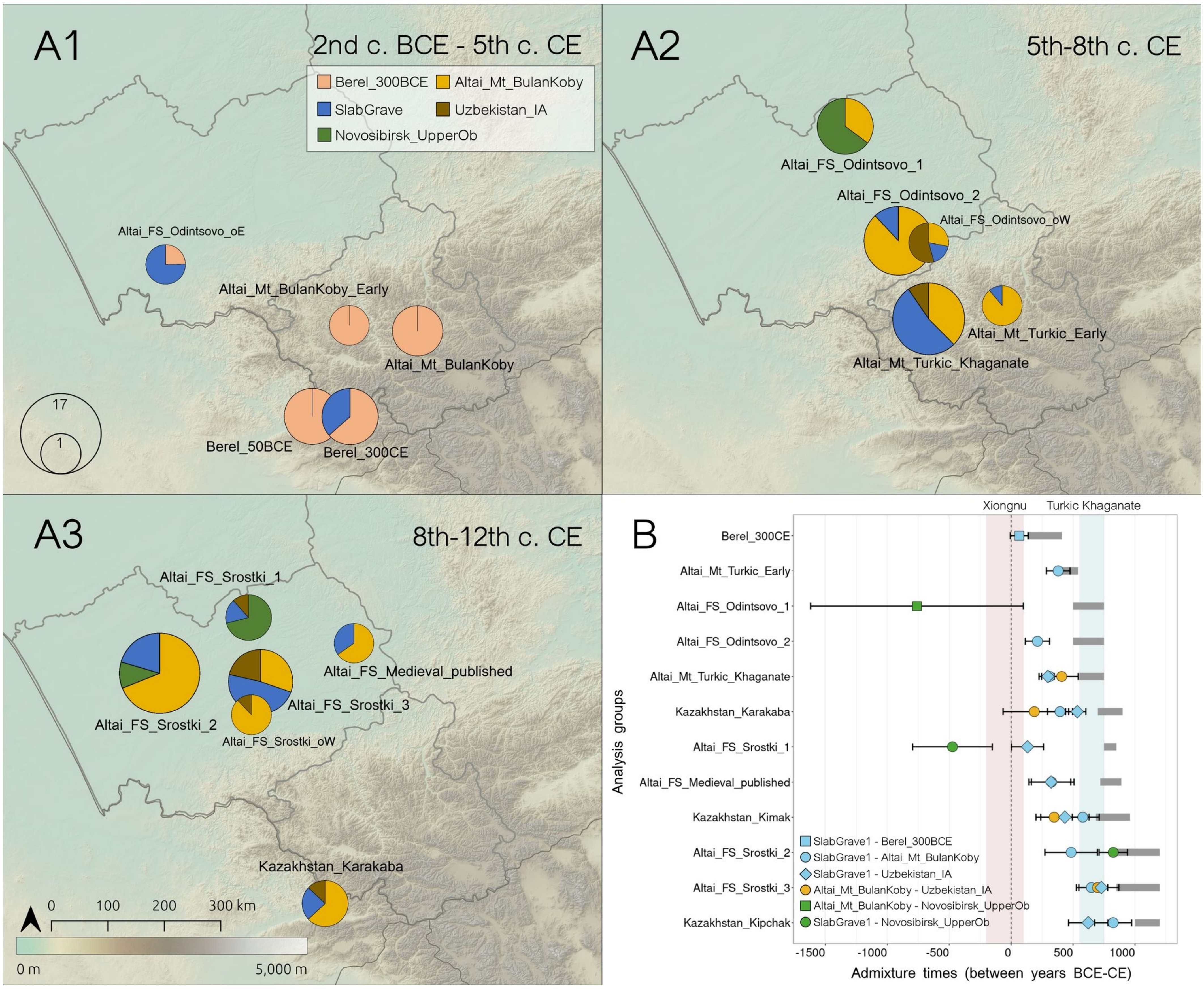


**Figure 3: Ancestry modeling and admixture times.**

(A1-3) Fitting (P≥0.05) lowest rank group-based qpAdm ancestry models of the newly-analysed Altai groups and outliers. The legend shows the designated color for the plotted source qpAdm groups. The right groups used in qpAdm analyses are named in the methods section. The circle sizes indicate the total individual number at a given analysis group. (B) Admixture times of the main analysis groups using DATES. The horizontal black bars show the mean admixture times ± standard errors (considering 29 years per generation), and the mean values are in coloured symbols which are based on the used source combinations (see legend). The horizontal gray bars show the archaeologically assigned dates of the analysis group. The results from DATES can be found in Supplementary Table S9.

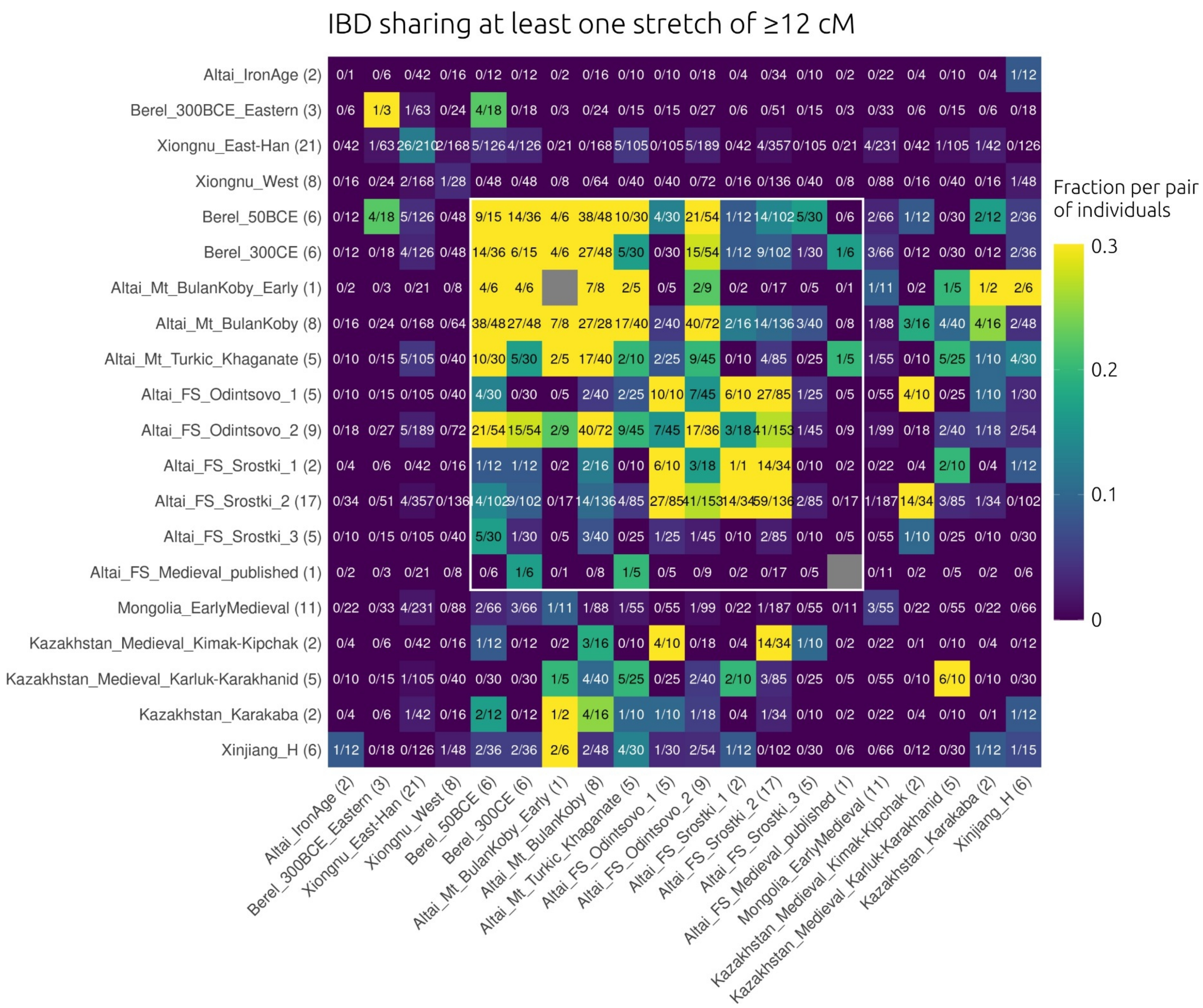


**Figure 4: IBD heatmap of the newly-analysed Altaians and other contextually-related groups.**

IBD sharing between chosen analysis groups in the 12 cM IBD dataset. Parentheses next to the group labels indicate the group sizes. The numbers in the cells represent (total detected IBD links)/(total possible pairwise links) and the colouring is based on normalization with the same fraction similar to Ringbauer et al., 2024 (*47*). A higher fraction means closer connectedness between the analysis groups. Groups of the post-IA Altai context are framed in white. The IBD analysis groups differ from the customary population genetic analysis groups (see Methods, Table S10).

**Supplementary Figures**

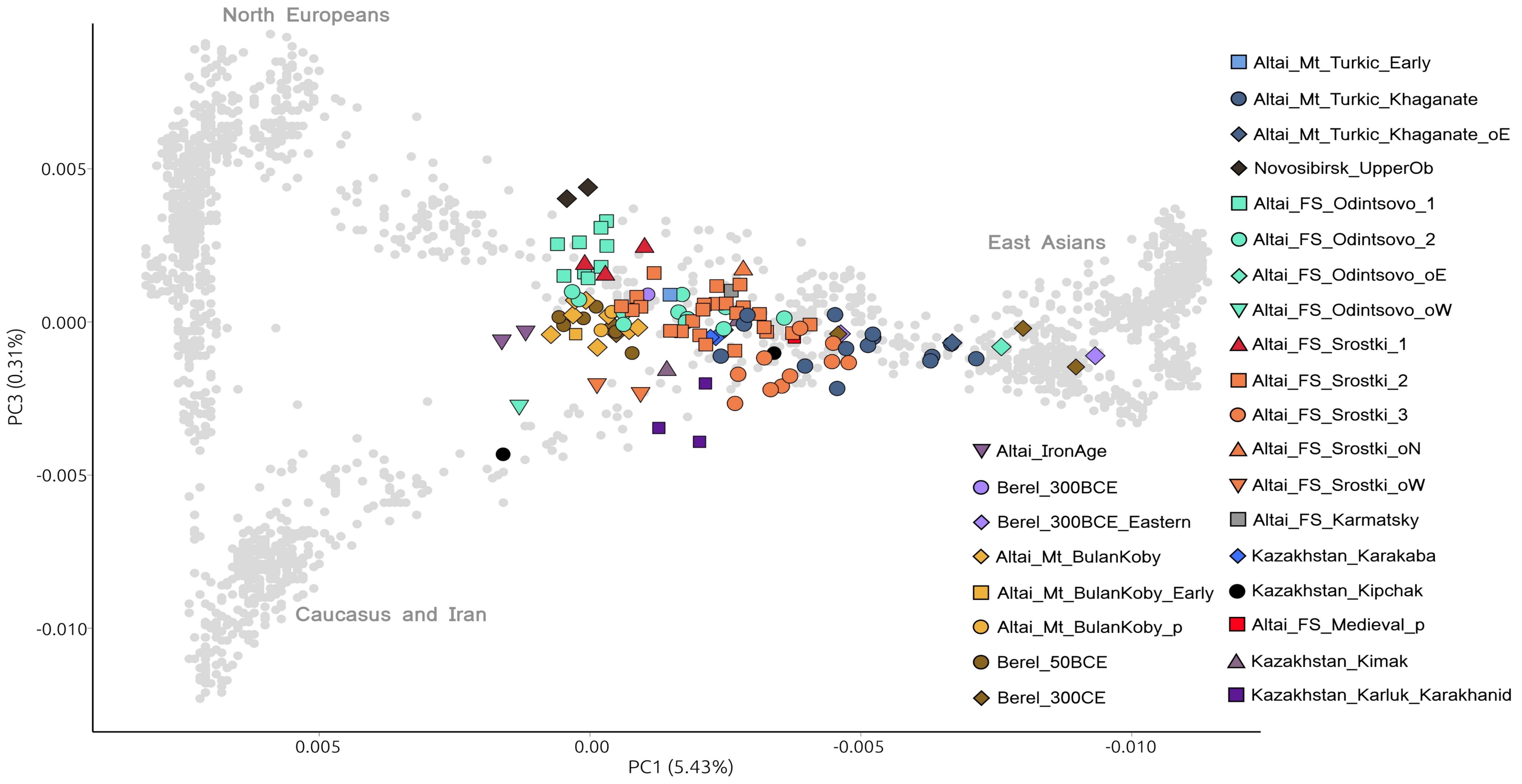


**Figure S2: PC1-PC3 variation PCA plot.**

Principal Component Analysis (PCA) plot of Eurasia on the PC1-PC3 space, based on modern Human Origin dataset, including newly reported and published genomes. Ancient genotypes were projected onto the modern genomic variation (gray points).

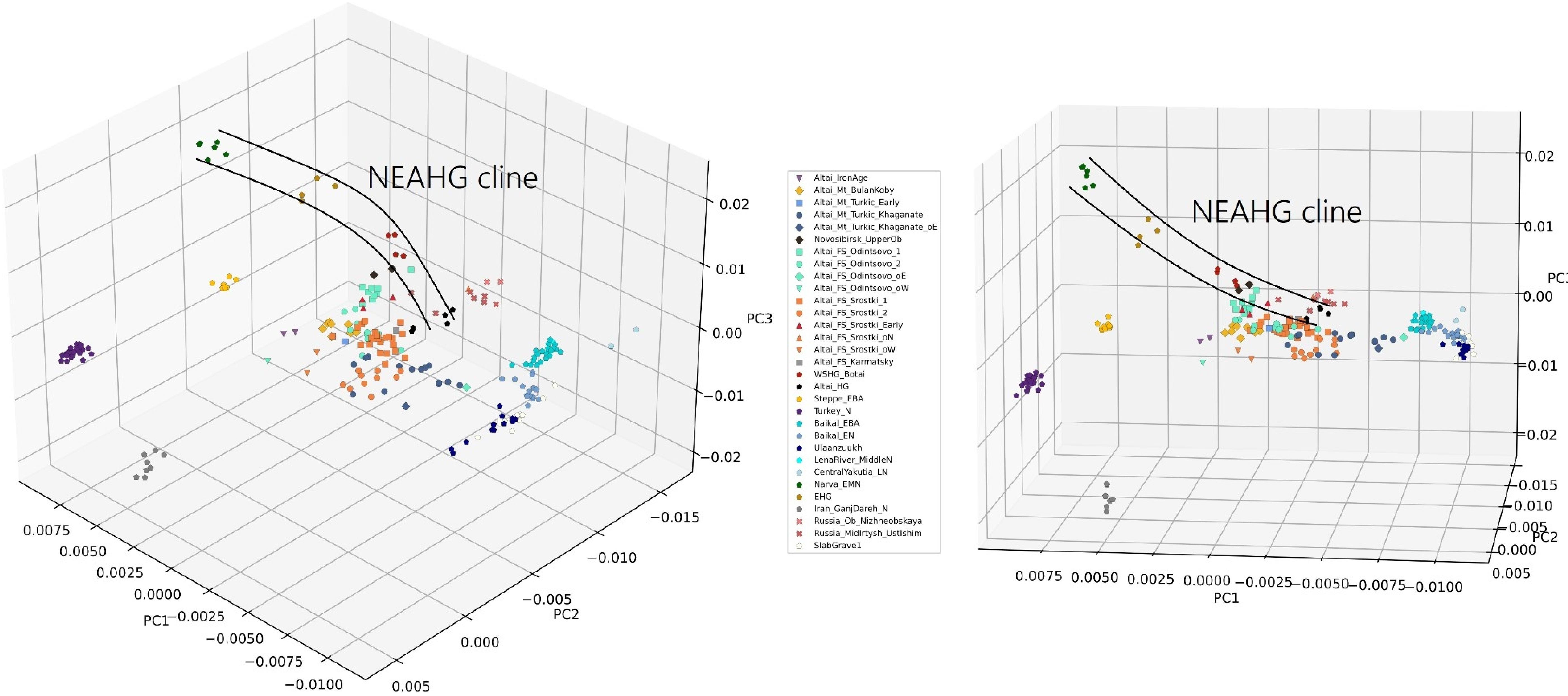


**Figure S3: 3D PCA plot of the newly published individuals with reference groups.**
3D PCA plot of the newly published dataset on the PC1 (5.43%) - PC2 (0.62%) - PC3 (0.31%) space. Some source groups, including five of six reference groups of supervised ADMIXTURE are given as references on the plot for the variety.

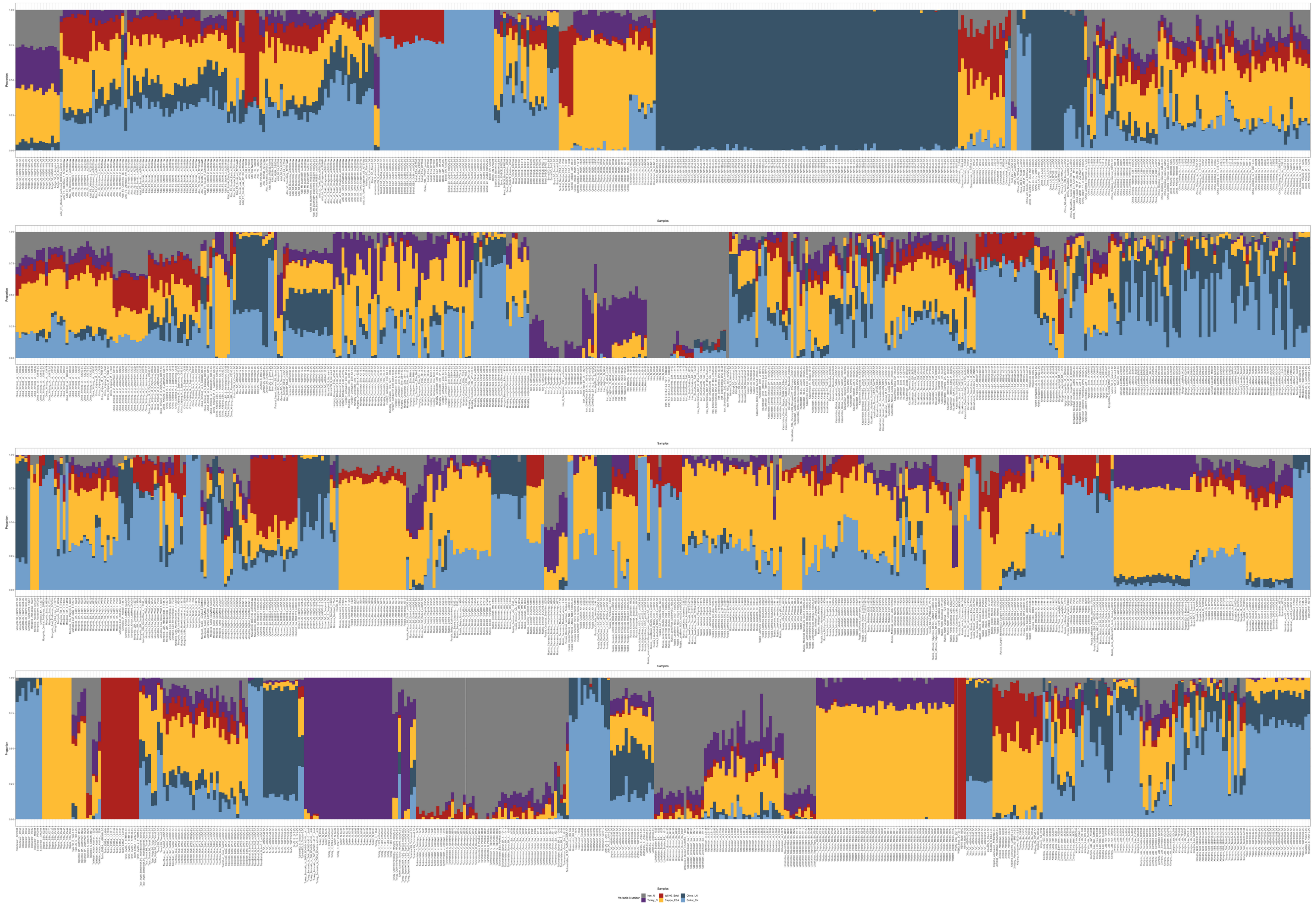

**Figure S4: ADMIXTURE plot of all samples in the dataset.**

K=6 supervised ADMIXTURE analysis results are presented for ancient and modern individuals. The whole list of the used groups can be found in Supplementary Table S2.

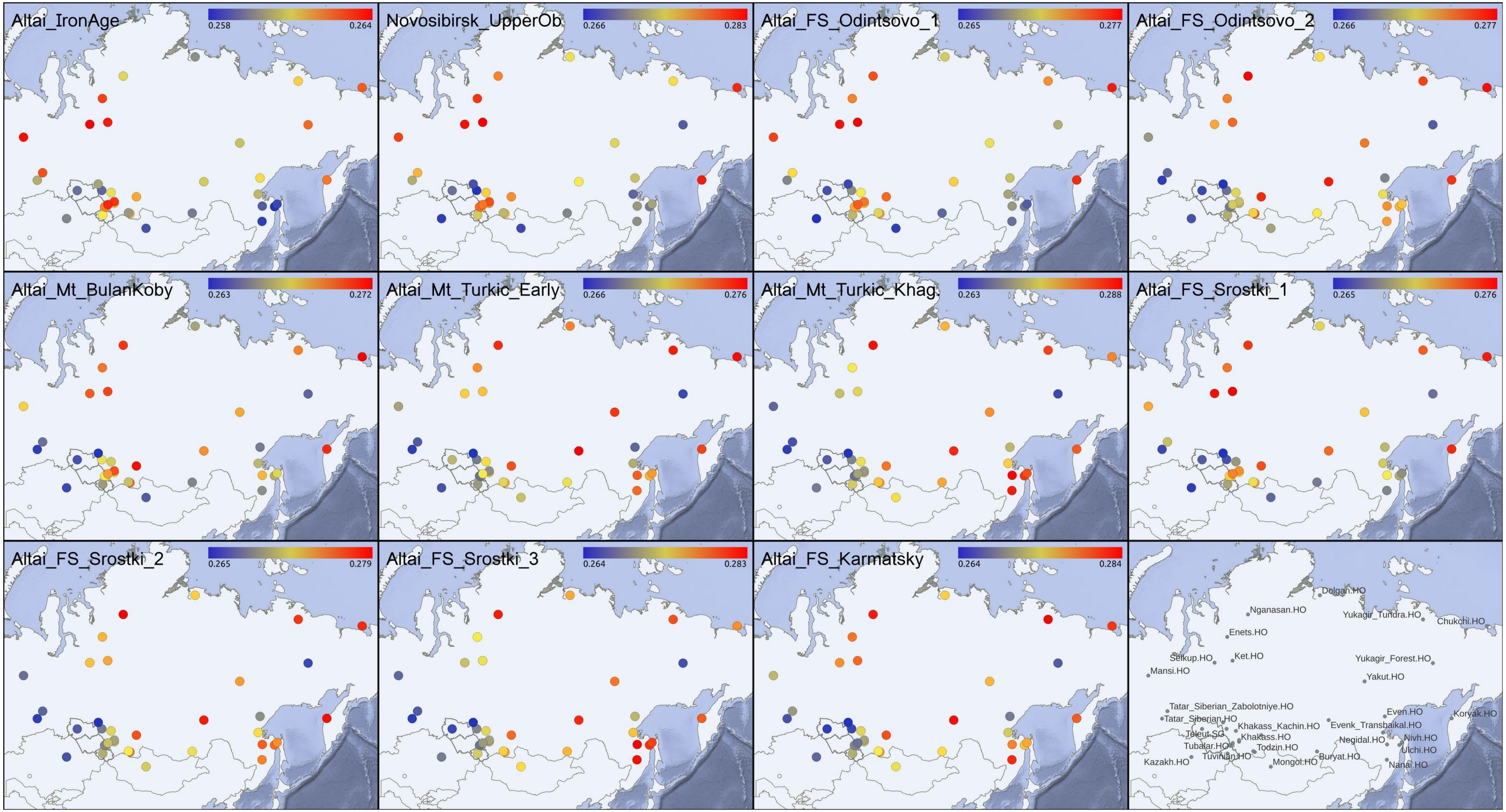


**Figure S5: Map of outgroup *f*3-statistics results for affinities to modern Siberians.**

Outgroup-$f_3$-statistics results of the newly presented ancient Altai analysis groups in the form $f_3$(x, Test; Mbuti.DG) for 35 modern Siberian populations are projected on the map, focusing on the Siberian region. The minimum and maximum values were given in the legend of each map, rounded for three decimals. Names of the projected modern groups are presented in the below-right corner. The findings are discussed in Supplementary Text S4, and the complete list of results can be found in Supplementary Table S4.

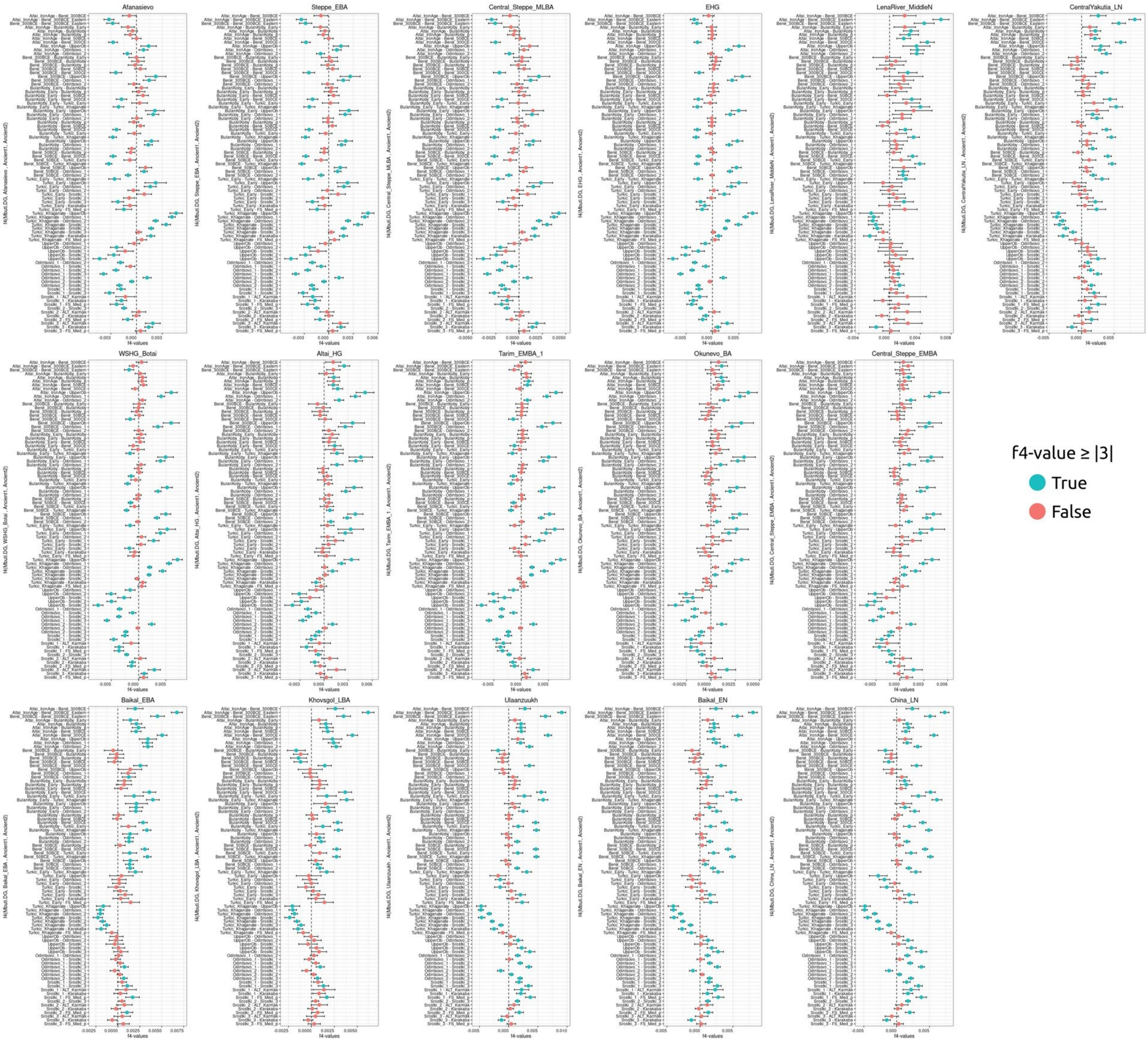


**Figure S6.1: $f_4$ comparisons of the deep ancestral populations of Inner/Central Asia for the analysis groups.**
$f_4$-statistics in the form $f_4$(Mbuti.DG, ancient group, Test1, Test2), designed to examine the cladality patterns of the deep ancestral populations, with the newly presented ancient Altai analysis groups (Test). The test combinations are sorted the same for each graph. For each deep ancestral group, positive values mean higher allele sharing with the second test group, negative values mean higher allele sharing with the first test group. Significance threshold was taken Z≥|3|. The findings are discussed in Supplementary Text S4, and the complete list of results can be found in Supplementary Table S5.

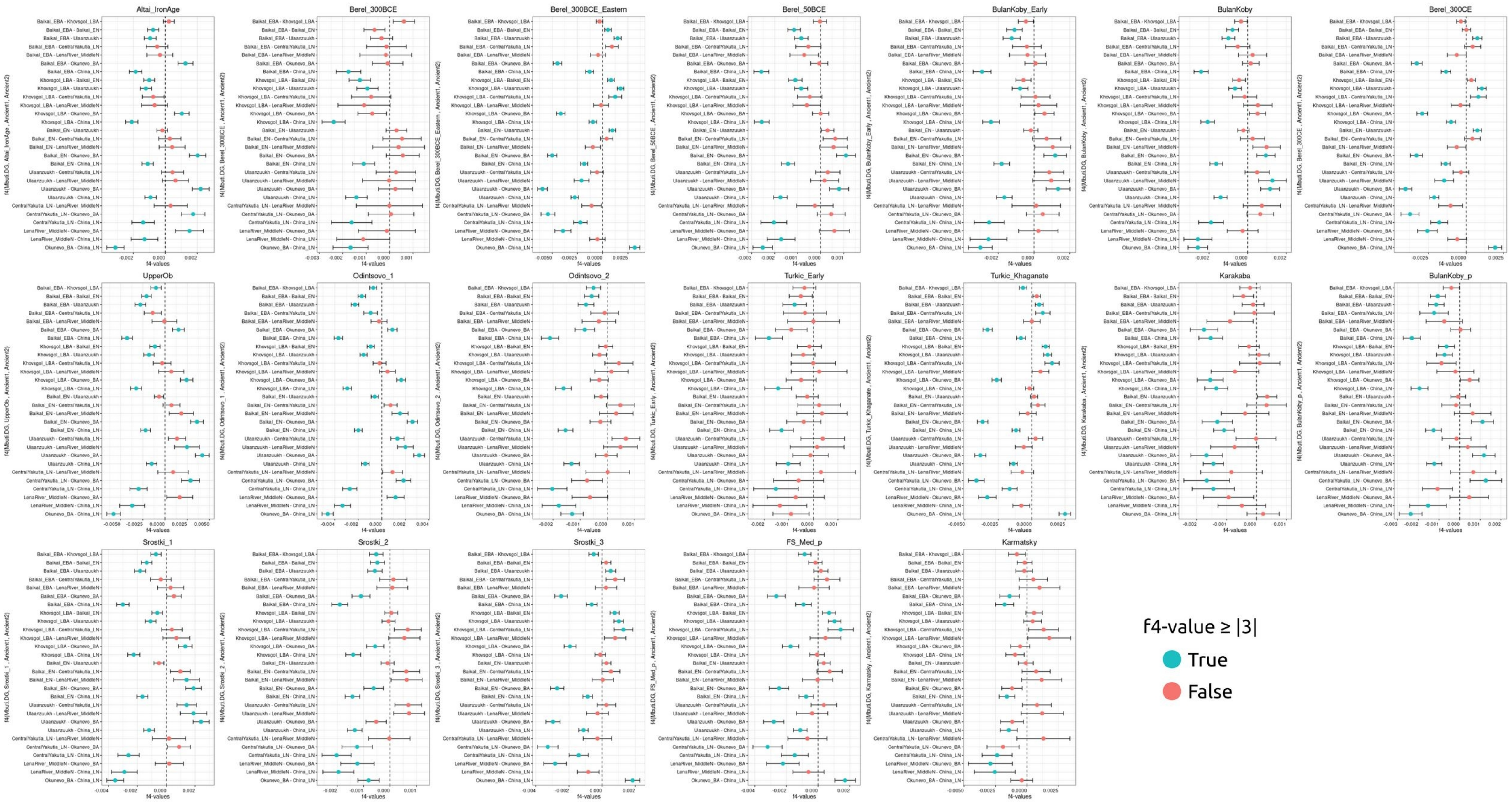


**Figure S6.2: $f_4$ comparisons of the analysis groups for deep ancestral East Asians.**

$f_4$-statistics in the form of $f_4$(Mbuti.DG, Test, ancient group1, ancient group2) to examine the cladality patterns of the newly presented ancient Altai analysis groups (Test), comparing between different deep ancestral East Asians (ancient group). The ancient group combinations are sorted the same for each graph. For each newly analysed group, positive values mean higher allele sharing with the second ancient group, negative values mean higher allele sharing with the first ancient group. Significance threshold was taken Z≥|3|. The findings were discussed in Supplementary Text S4, and the complete list of results can be found in Supplementary Table S5.

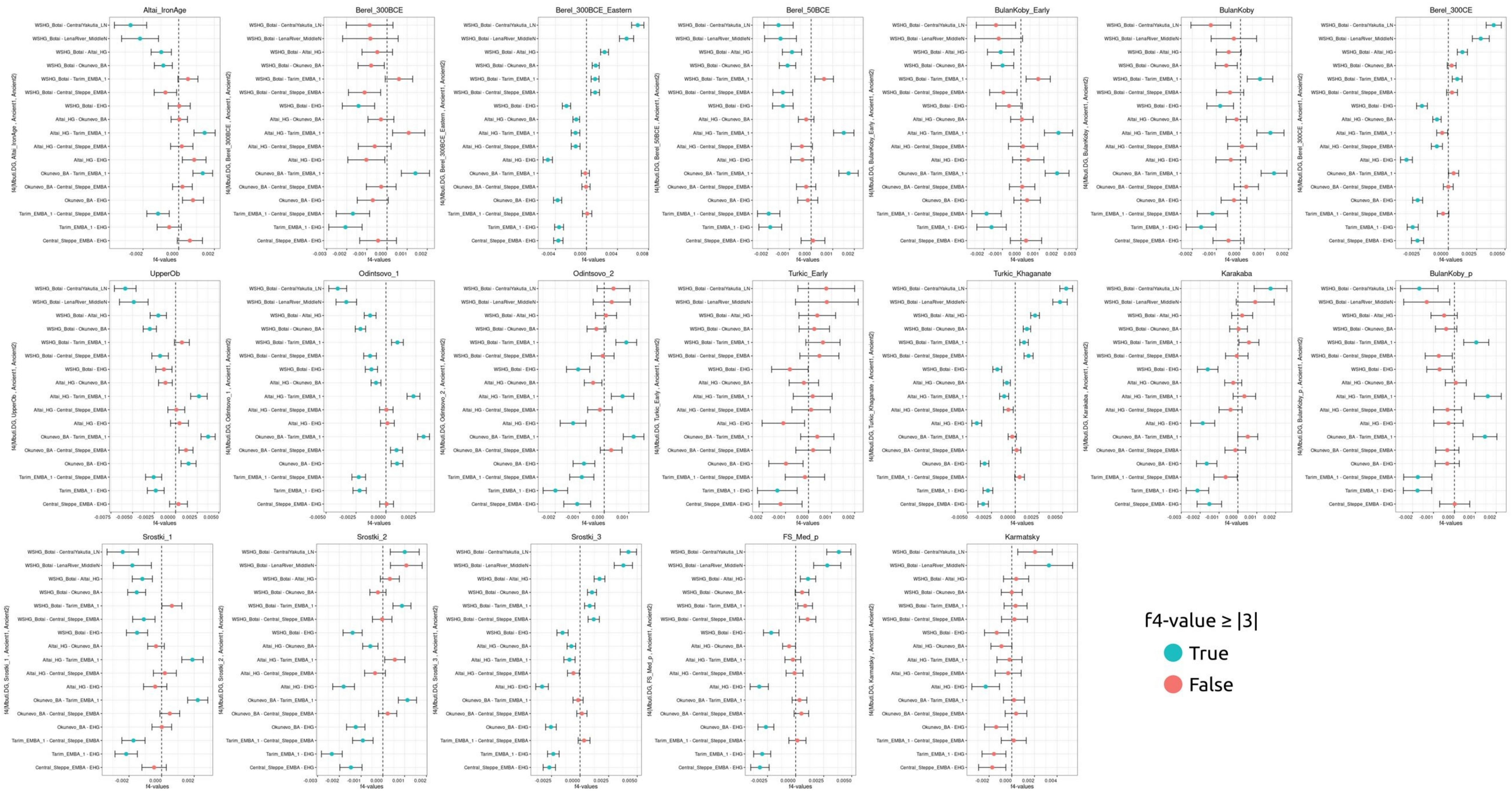


**Figure S6.3: $f_4$ comparisons of the analysis groups for deep ancestral Ancient North Eurasians.**
$f_4$-statistics in the form of $f_4$(Mbuti.DG, Test, ancient group1, ancient group2) to examine the cladality patterns of the newly presented ancient Altai analysis groups (Test), comparing between different deep ancestral Ancient North Eurasians (ancient group). The ancient group combinations are sorted the same way for each graph. For each newly analysed group, positive values mean higher allele sharing with the second ancient group, negative values mean higher allele sharing with the first ancient group. Significance threshold was taken Z≥|3|. The findings were discussed in Supplementary Text S4, and the complete list of results can be found in Supplementary Table S5.

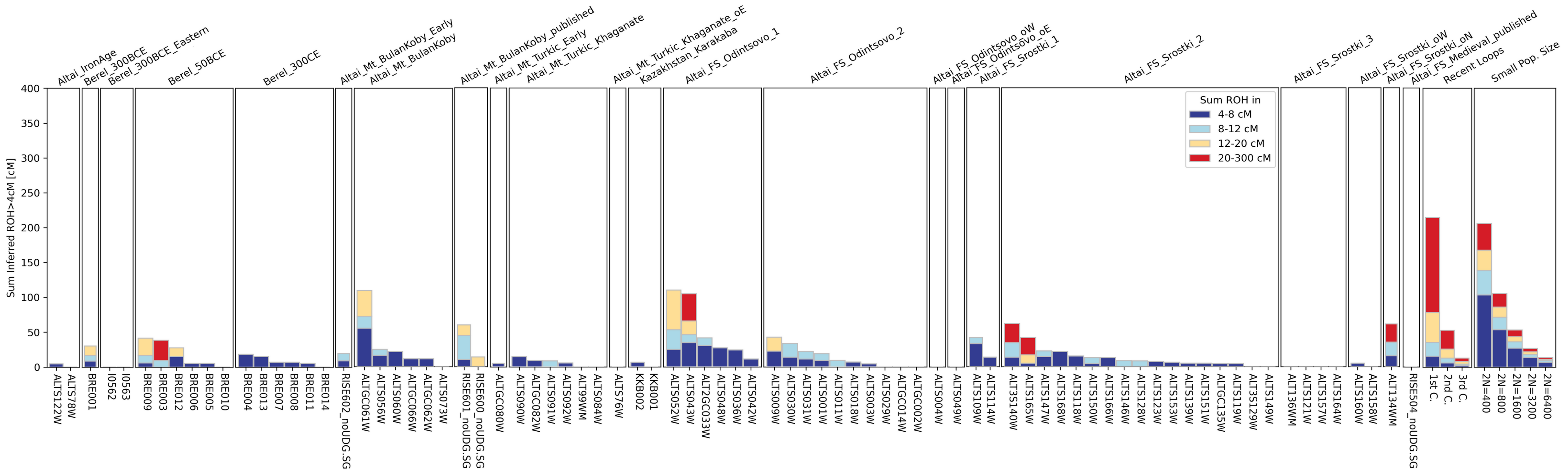


**Figure S7: hapROH multiplot of the Altai groups.**

Runs of homozygosity (ROH) results of the newly-presented and previously published Altai individuals analysed in this study. Long homozygous runs are indicative of parental relatedness, while signals for shorter runs indicate lower effective population sizes (*Ne*). The *Ne* estimations can be found in Supplementary Table S8.

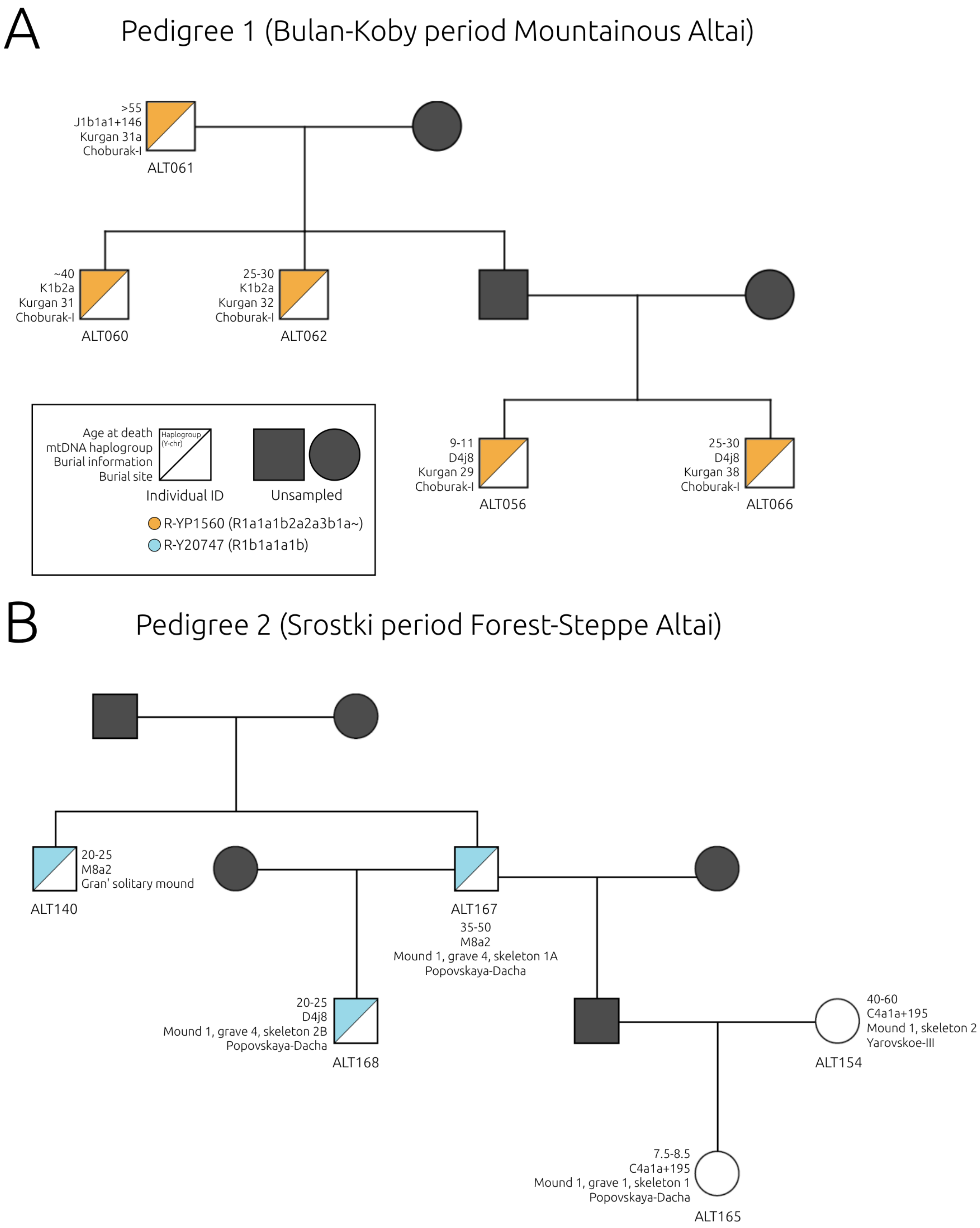


**Figure S8: Two pedigrees discovered in the newly-analysed dataset.**
Pedigrees reconstructed using the kinship estimates. (A) Pedigree 1 from the Bulan-Koby culture and (B) Pedigree 2 from the Srostki culture both covered three generations. Y-chromosomal haplogroups are given with colors matching the legend, on the top-left half of the male (square) symbols. Dark gray symbols show individuals lacking from the analysed dataset. The full list of kinship estimates can be found in Supplementary Table S9.

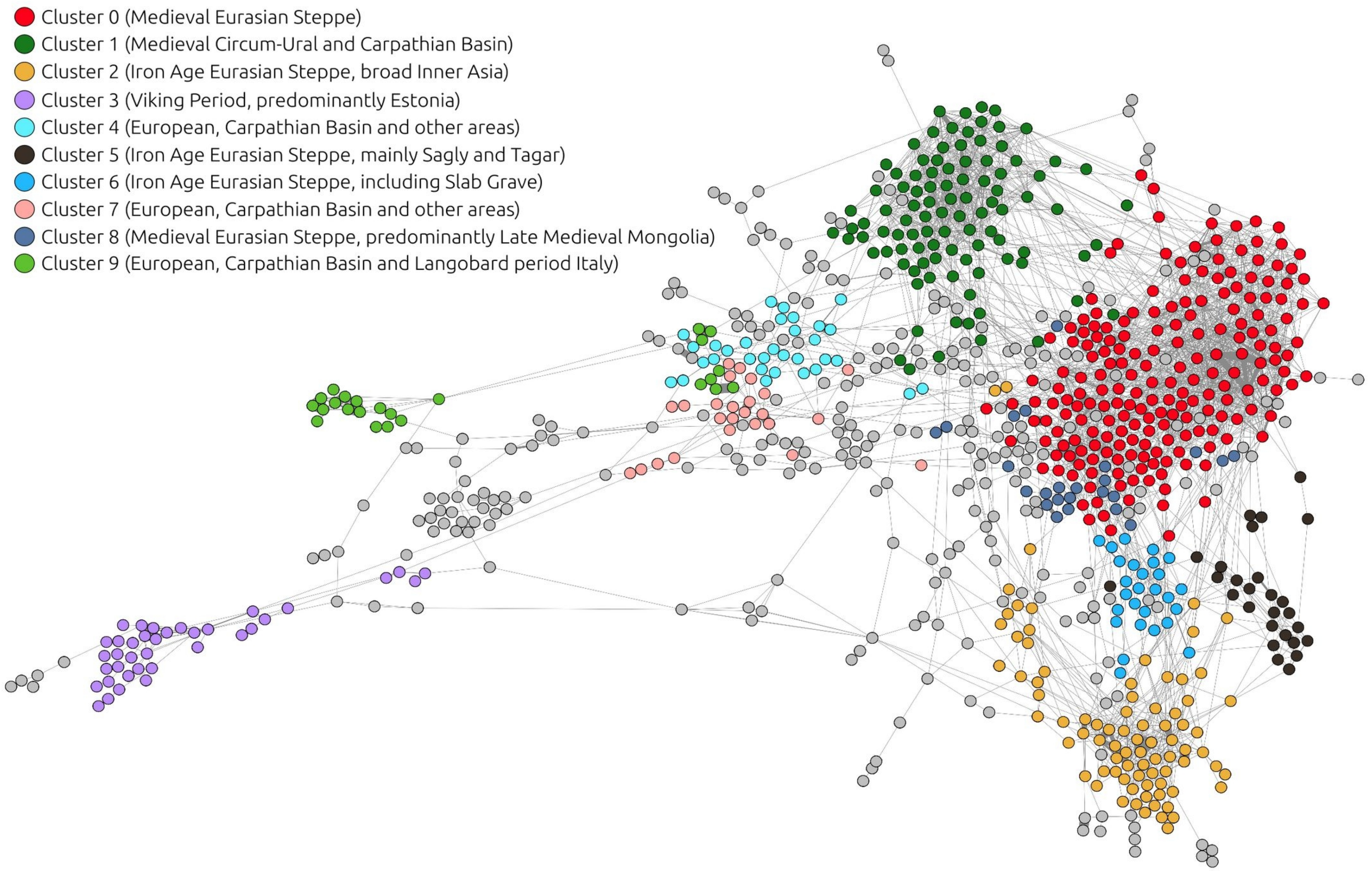


**Figure S9: IBD graph of ancient individuals and IBD clusters.**

IBD-sharing network graph of 846 connected ancient Eurasian individuals, dated to between the Iron Age and pre-modern eras. Ten largest IBD sharing clusters are visualized in color. The network is constructed using minimum sum≥1x12 cM IBD connections per individual pair. For the layout algorithm and clustering, see Methods. The full list and information of the individuals (nodes), and the IBD-links (edges) between these individuals are given in Supplementary Tables S10 and S11.

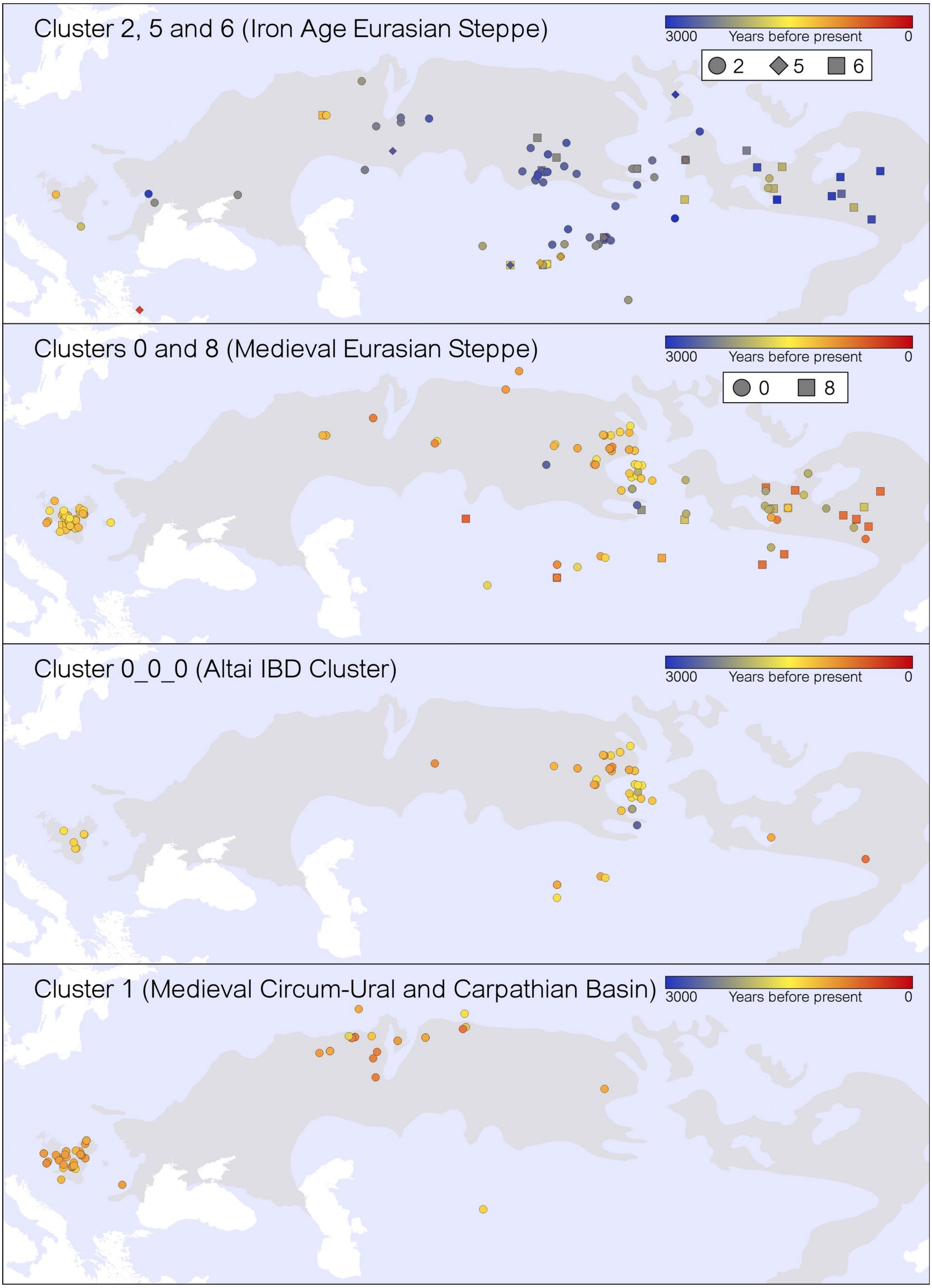


**Figure S10: IBD clusters on the Eurasian map.**
IBD clusters related to the steppe and Altai context are projected on the map. The steppe and forest-steppe regions are coloured on top of the base map. Contemporaneous clusters are also observed to be overlapping on some regions, including some shared burial sites.

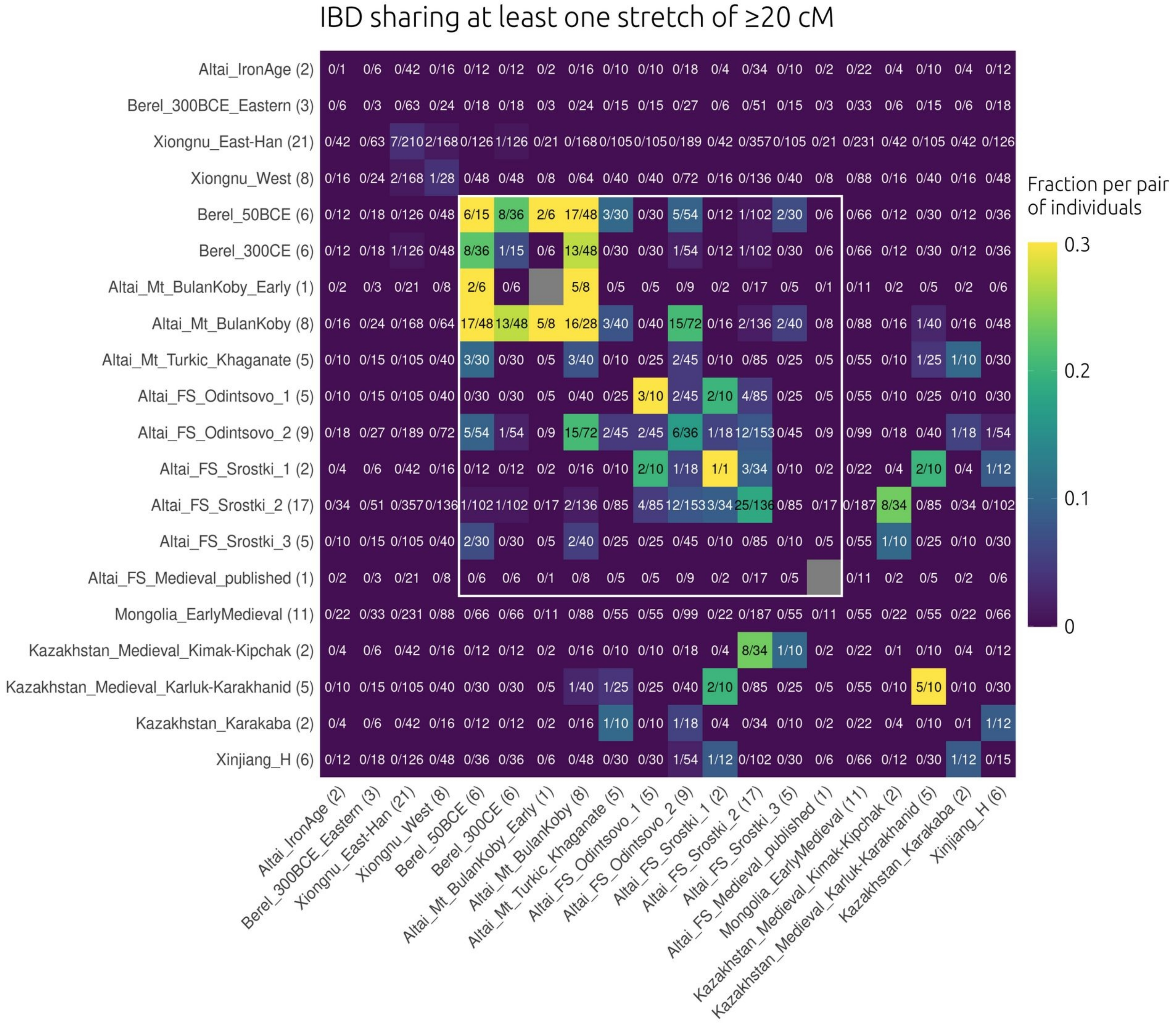


**Figure S11: IBD heatmap graph (≥1x20 cM).**
IBD sharing between chosen analysis groups in the 20 cM IBD dataset. Parentheses next to the group labels indicate the group sizes. The numbers in the cells represent (total detected IBD links)/(total possible pairwise links) and the colouring is based on normalization with the same fraction similar to Ringbauer et al. 2024. A higher fraction means closer connectedness between the analysis groups. Groups of the post-IA Altai context are framed in white. The IBD analysis groups differ from the customary population genetic analysis groups (see Methods, Table S10).

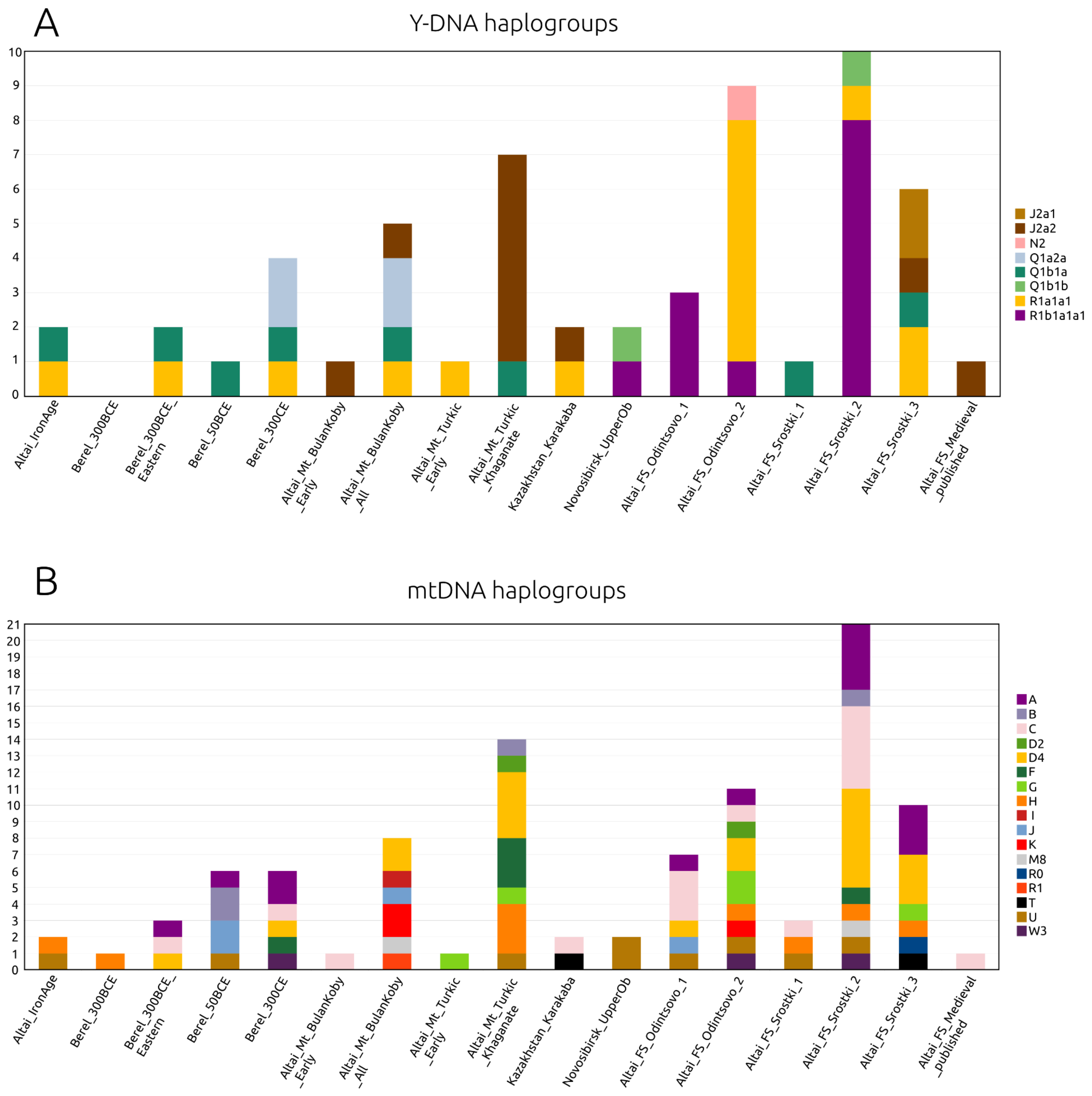


**Figure S12: Uniparental haplogroups of the Altai groups.**
Distribution of (A) Y-chromosomal haplogroups and (B) mitochondrial haplogroups across the ancient Altaian groups. Individuals were grouped based on the analysis groups as listed in Supplementary Tables S1 and S2.

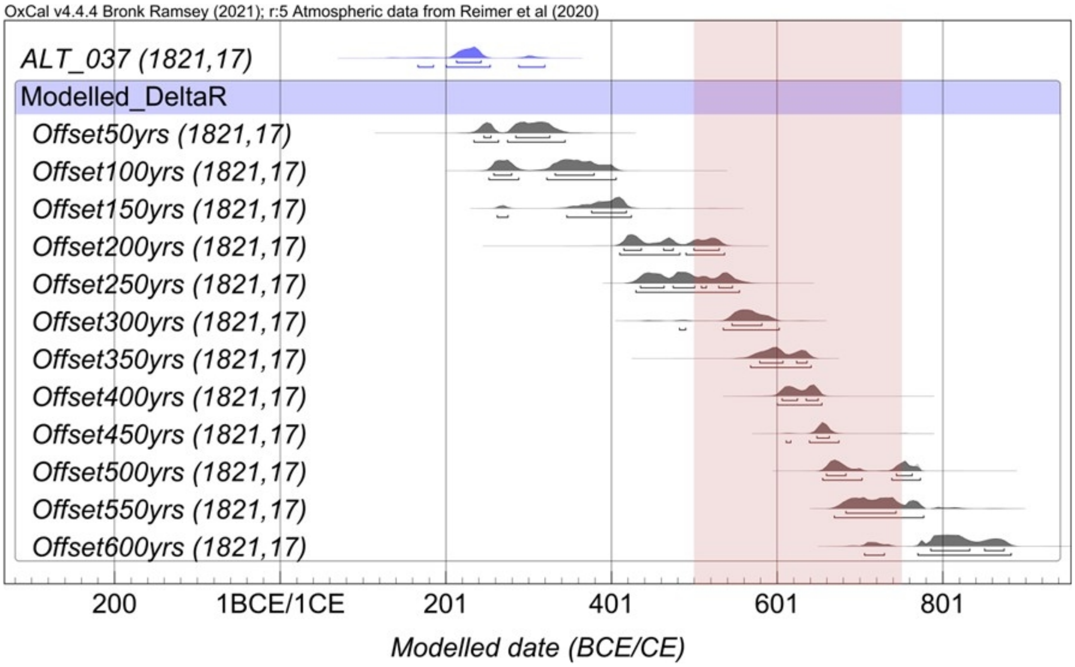


**Figure S13.1:** ΔR modeling through comparison of the measured [14]C date for ALT037 (blue) and the expected age period of 500-750 CE (marked with the reddish area).

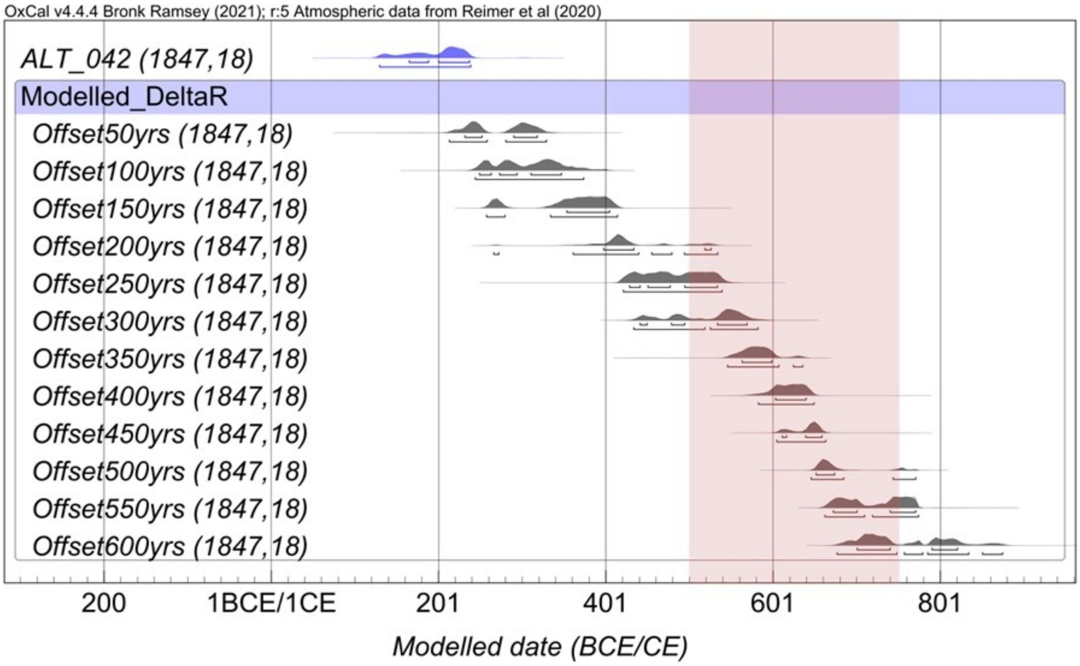


**Figure S13.2:** ΔR modeling through comparison of the measured [14]C date for ALT042 (blue) and the expected age period of 500-750 CE (marked with the reddish area).

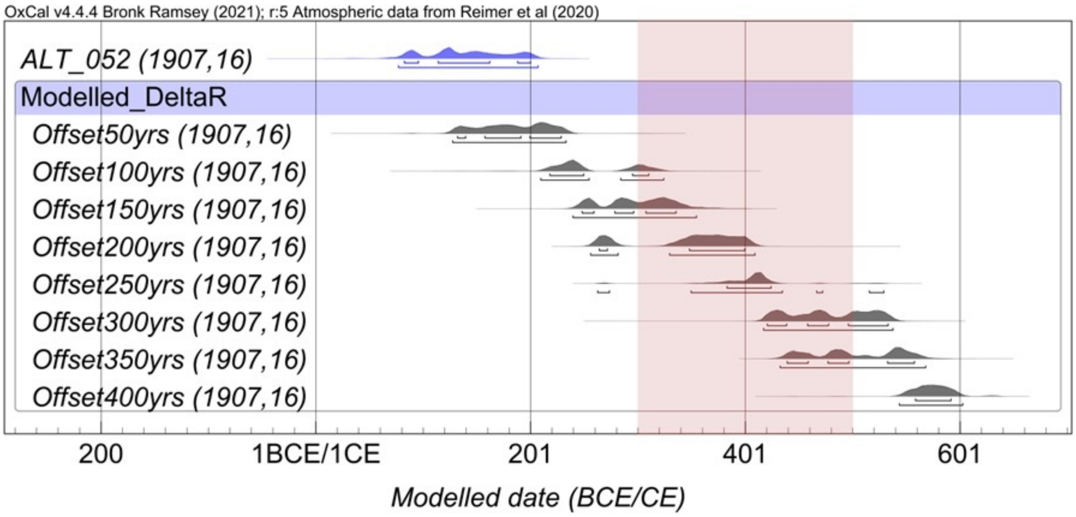


**Figure S13.3:** ΔR modeling through comparison of the measured $^{14}C$ date for ALT052 (blue) and the expected age period of 300-500 CE (marked with the reddish area).

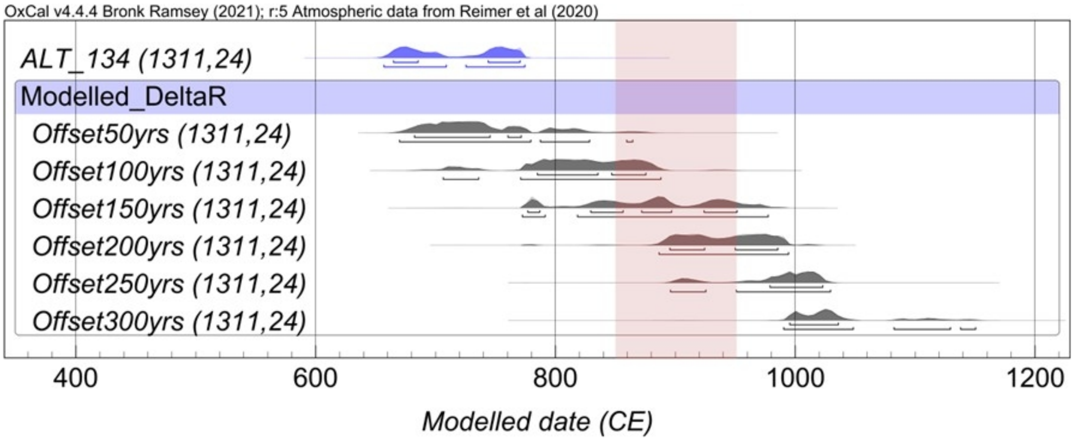


**Figure S13.4:** ΔR modeling through comparison of the measured $^{14}C$ date for ALT0134 (blue) and the expected age period of 850-950 CE (marked with the reddish area).

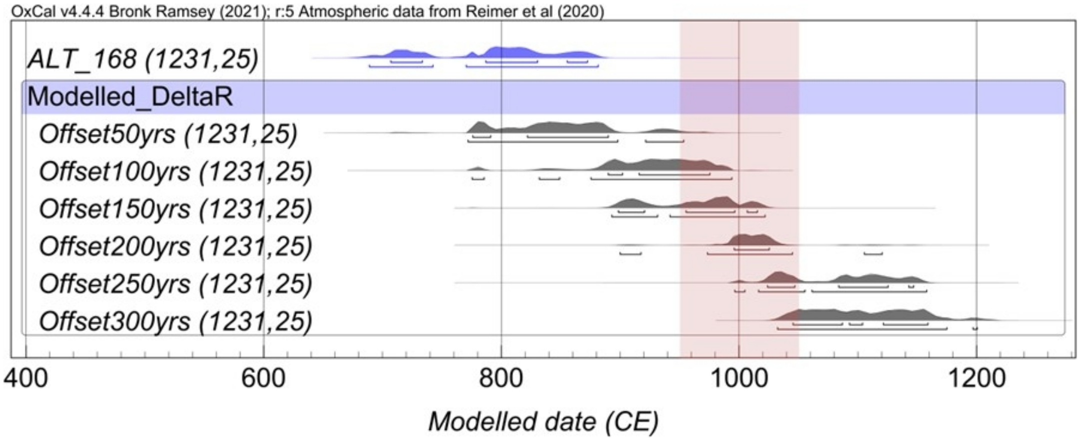


**Figure S13.5:** ΔR modeling through comparison of the measured $^{14}C$ date for ALT168 (blue) and the expected age period of 950-1050 CE (marked with the reddish area).
